# Kinetic and structural characterization of the self-labeling protein tags HaloTag7, SNAP-tag and CLIP-tag

**DOI:** 10.1101/2021.04.13.439540

**Authors:** Jonas Wilhelm, Stefanie Kühn, Miroslaw Tarnawski, Guillaume Gotthard, Jana Tünnermann, Timo Tänzer, Julie Karpenko, Nicole Mertes, Lin Xue, Ulrike Uhrig, Jochen Reinstein, Julien Hiblot, Kai Johnsson

## Abstract

The self-labeling protein tags (SLPs) HaloTag7, SNAP-tag and CLIP-tag allow the covalent labeling of fusion proteins with synthetic molecules for applications in bioimaging and biotechnology. To guide the selection of an SLP-substrate pair and provide guidelines for the design of substrates, we report a systematic and comparative study on the labeling kinetics and substrate specificities of HaloTag7, SNAP-tag and CLIP-tag. HaloTag7 reaches almost diffusion-limited labeling rates with certain rhodamine substrates, which are more than two orders of magnitude higher than those of SNAP-tag for the corresponding substrates. SNAP-tag labeling rates however are less affected by the structure of the label than those of HaloTag7, which vary over six orders of magnitude for commonly employed substrates. Solving the crystal structures of HaloTag7 and SNAP-tag labeled with fluorescent substrates allowed us to rationalize their substrate preferences. We also demonstrate how these insights can be exploited to design substrates with improved labeling kinetics.

## Introduction

Modern high-resolution fluorescence imaging techniques require the specific labeling of proteins with appropriate fluorescent probes. Self-labeling protein tags (SLPs) have been shown to offer a straightforward way to achieve this goal as they undergo a specific and irreversible reaction with synthetic substrates such as fluorophores (1). SLPs are furthermore employed in various other applications such as *in vitro* biophysical studies (2, 3), the generation of semisynthetic biosensors (4–7) and yeast three-hybrid screenings (8). The three most popular SLPs are HaloTag7 (HT7) (9), SNAP-tag (SNAP) (10) and CLIP-tag (CLIP) (11) (**Fig. 1**).

HT7 was engineered from a bacterial dehalogenase (DhaA from *Rhodococcus sp*.), an enzyme able to hydrolyze halogenated alkanes (12). Inactivating the second catalytic step of its enzymatic reaction (mutation H272N in HT7) abolished the hydrolysis of the ester formed with an active site aspartate residue and created an SLP. HT7 reacts specifically with chloroalkane-PEG (CA) molecules resulting in covalent bonding of the alkane chain to the catalytic aspartate and release of a chloride ion (**Fig. 1A**). HT7 was further engineered for increased stability and efficient labeling kinetics toward CA-fluorophore substrates (13).

SNAP was engineered from the human *O^6^*-alkylguanine-DNA alkyltransferase (hAGT), a protein involved in the repair of alkylated DNA by transferring alkyl moieties to its reactive cysteine (14). SNAP was engineered to efficiently react with benzylguanine (BG) derivatives as substrates (**Fig. 1B**) and to reduce its DNA binding properties (10). SNAP irreversibly transfers the benzyl moiety of the substrate to its reactive cysteine, leading to the release of guanine. SNAP also accepts substrates in which the guanine is replaced by a chloropyrimidine (CP) (**Fig. 1B**), reported to possess higher cell permeability (15). Later, CLIP was engineered from SNAP as an orthogonal SLP system, accepting benzylcytosine (BC) derivatives as substrates (11) (**Fig. 1B**).

Even though it has become clear over the last years that the nature of the transferred label can have a significant impact on the reaction kinetics (9, 16, 17), no systematic study has been reported so far that addresses the influence of the transferred label on the SLP labeling kinetics. The structural reasons for the differences in labeling rates are poorly understood. Furthermore, the reaction kinetics of SLPs are usually characterized as a single step-reaction under pseudo-first order reaction conditions, *i.e*. in large excess of one of the reactants (Model 1, **Fig. 1C**). However, the reaction mechanism of SLPs is more complex and should be characterized by a multi-step kinetic model comprising reversible substrate binding (k_1_), unbinding (k_−1_) and irreversible covalent reaction (k_2_) (Model 2, **Fig. 1C**). Here, we report an in-depth characterization of the reaction kinetics of HT7, SNAP and CLIP with different substrates, identifying those structural features of labels that control labeling rates for the different tags. We complement these kinetic studies by reporting crystal structures of HT7 and SNAP covalently labeled with rhodamine-based fluorophores, providing a detailed understanding of their substrate preferences. Our results will (i) facilitate the use of SLPs in various applications, (ii) aid in the SLP engineering and (iii) help in the design of improved labeling substrates.

**Figure 1:**
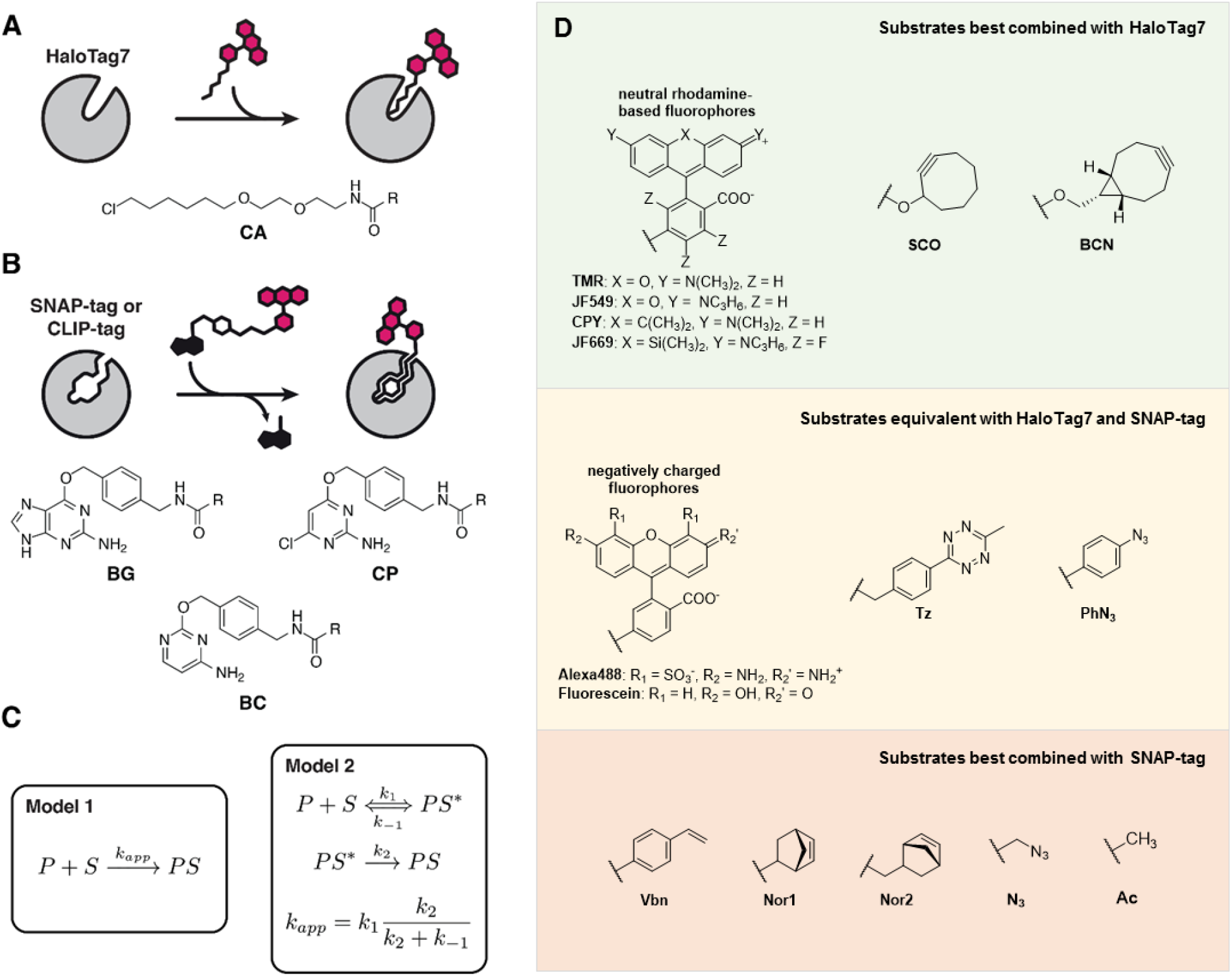
Self-labeling reaction, substrates and kinetic models. **A**. Scheme of HT7 labeling reaction with fluorophore substrates. The chemical structure of HT7 substrates (CA) is depicted below. R represents the functional moiety to be linked to HT7. **B.** Scheme of SNAP(f) / CLIP(f) labeling reaction with fluorophore substrates. The chemical structures of SNAP/CLIP substrates (BG/CP/BC) are depicted bellow. R represents the functional moiety to be linked to the SLP. **C**. Models employed to describe the SLP kinetics in this study. **D.** Chemical structure of different SLP substrate substituents. Substrates are organized by their preferential use with HT7 or SNAP.

## Results

### HaloTag7 kinetic characterization

Fluorophores represent the most popular class of labels employed with SLPs. We characterized HT7 labeling kinetics with different CA-fluorophore substrates, namely CA-TMR, CA-JF549, CA-LIVE580, CA-CPY, CA-JF669 and CA-Alexa488 (**Fig. 1D & S1**) by tracking fluorescence anisotropy change over time at different reactant concentrations. The very high labeling speed of HT7 towards most rhodamine-based CA substrates required a stopped-flow setup to precisely measure the labeling kinetics. Data were fitted to the kinetic model 2 (**Fig. 1C**), which described the reaction kinetics of most rhodamine-based HT7 substrates and allowed to determine the three kinetic parameters (k_1_, k_−1_ & k_2_) independently (**Fig. 2A-C, S2 & Table S1**). Data fitted to the simplified model 1 resulted in a poorer fit, since curves show a clear biphasic character, indicating that model 2 should be preferred to describe these fast labeling kinetics (**Fig. S3**). It should be noted that fitting the data for the faster reacting substrates to model 1 would lead to a significant overestimation of the labeling speed (**Fig. S4 & Table S2**). The slower labeling reaction with CA-Alexa488 allowed to perform measurements in a microplate reader. However, fitting model 2 to this data does not allow to determine the kinetic parameters (k_1_, k_−1_ & k_2_) independently. Hence the data was fitted using the kinetic model 1 (**Fig. S5**). The kinetic model 1 yields the apparent second-order rate constant k_app_ which describes the labeling reaction at reactant concentrations far below the K_d_ at which the substrate binding site is not saturated and the labeling rate depends linearly on the reactant concentrations. To compare the labeling rate constants of substrates analyzed through different kinetic models (**Fig. 2D & Table 1**), k_app_ can also be calculated from the individual rate constants obtained with kinetic model 2 (**Fig. 1C**).

**Figure 2:**
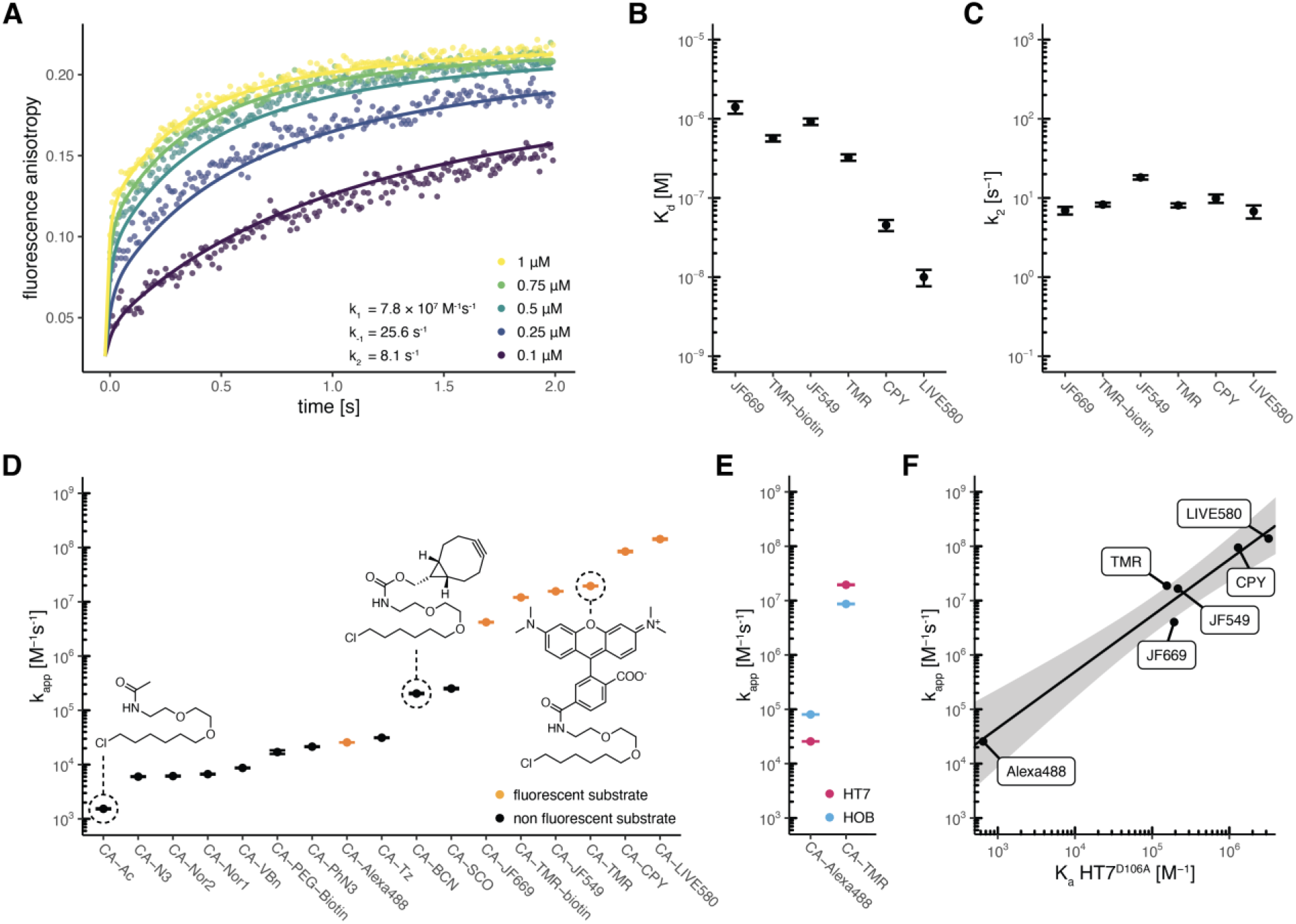
Characterization of HaloTag7 labeling kinetics. **A**. Fluorescence anisotropy traces (points) and fitted curves of HT7 labeling with CA-TMR in 1:1 stoichiometry at the indicated concentrations. Kinetics were recorded by following fluorescence anisotropy over time using a stopped flow device. Reactions were started by mixing equal volumes of HT7 and CA-TMR. Data were fitted to the kinetic model 2 (lines). **B**. HT7 affinities (K_d_) for different fluorophore substrates calculated from the kinetic parameters (k_−1_/k_1_). **C**. HT7 reactivity (k_2_) for different fluorophore substrates obtained from fluorescence anisotropy kinetics. The minimal differences in k_2_ illustrate that labeling kinetics are mostly influenced by differences in K_d_. **D**. Apparent second order labeling rate constants (kapp) of HT7 with different substrates. Rate constants span over six order of magnitude. Non-negatively charged fluorophore substrates reach the fastest labeling kinetics. **E**. Comparison of k_app_ between HT7 and HOB for CA-TMR and CA-Alexa488 labeling highlighting the preference of HOB for the negatively charged substrate CA-Alexa488. **F**. Correlation between HT7 apparent second order rate constant (k_app_) and affinity (K_a_ = 1/K_d_) for different fluorophore substrates. Affinities were obtained with the catalytically inactive variant HT7^D106A^. Log transformed values were fitted to a linear model (black line, log(k_app_) = log(K_a_) x 1.042 + 1.544). The grey area represents the 95% confidence bands (the area in which the true regression line lies with 95% confidence).

**Table 1:**
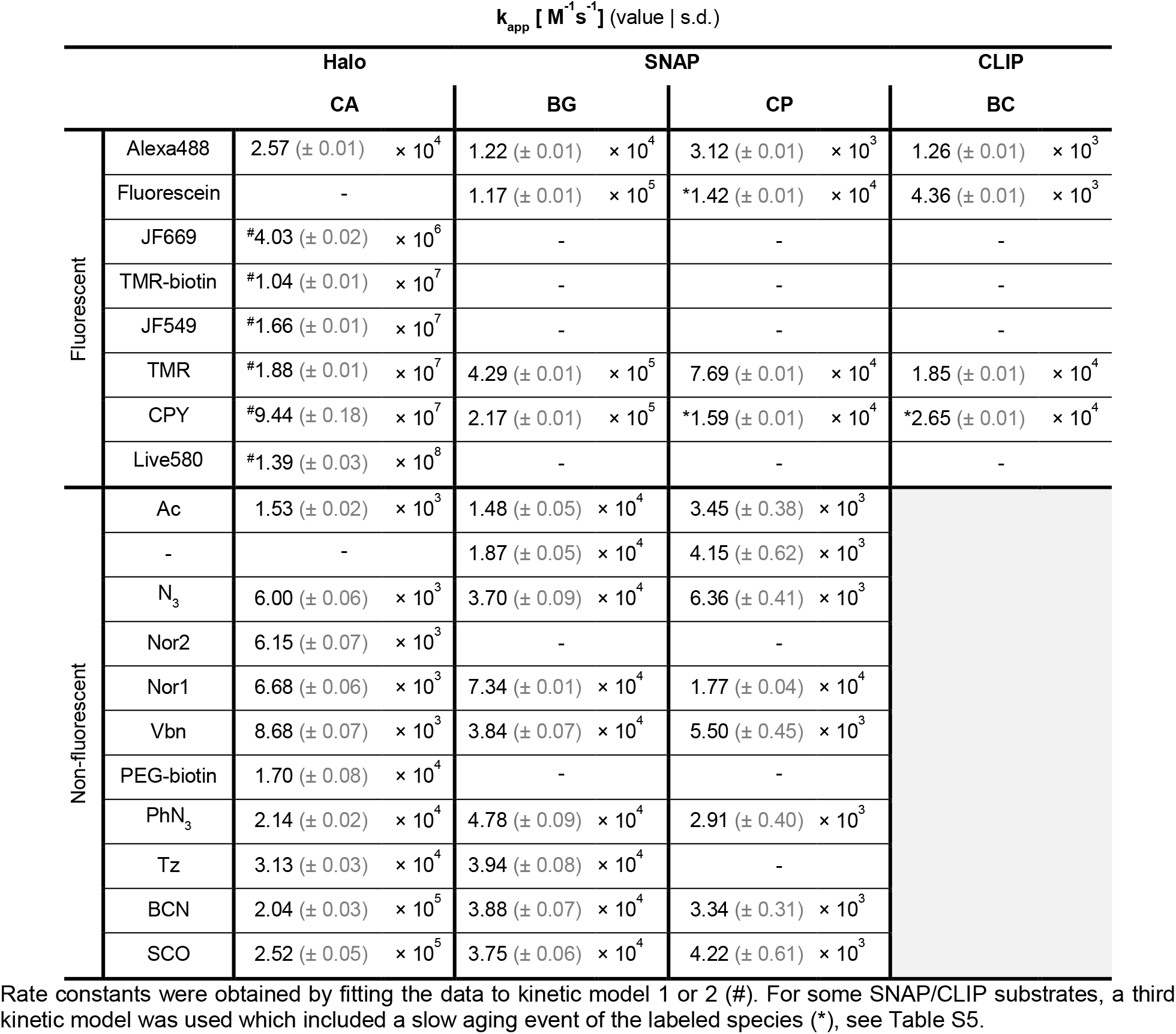
Apparent labeling rate constants (k_app_) for different HT7, SNAP and CLIP substrates.

### HaloTag7 reaches fast kinetics with fluorophore substrates

Among the tested fluorophore substrates, CA-LIVE580 turned out to be the fastest substrate for HT7 with a k_app_ of 1.39 ± 0.03 ×10^8^ M^−1^s^−1^, reaching an almost diffusion-limited labeling rate, and a calculated K_d_ (= k_−1_/k_1_) of 9.99 nM (7.64 to 12.35 nM 95% CI). All other rhodamine-based substrates showed efficient labeling kinetics as well (10^6^ < k_app_ < 10^9^ M^−1^s^−1^) with the exception of the negatively charged CA-Alexa488 (k_app_ = 2.57 ± 0.01 × 10^4^ M^−1^s^−1^) (**Table 1 & Fig. 2D**). The HT7 variant HOB (halo-based oligonucleotide binder) (18) features several positively charged surface mutations close to the substrate binding site, which were introduced to increase the labeling rates with chloroalkanes attached to oligonucleotides. We hypothesized that HOB may have increased labeling kinetics with the negatively charged CA-Alexa488. Indeed, HOB shows a 3.13 ± 0.01 fold increase in k_app_ compared to HT7 with CA-Alexa488, while a decrease in k_app_ was observed with CA-TMR (2.09 ± 0.01 fold) (**Fig. 1E, S5 & Table S3**). This suggests that kinetics of negatively charged substrates might suffer from charge repulsions at the HT7 surface.

### HaloTag7 labeling kinetics correlate with substrate affinity

For the substrates whose labeling kinetics followed model 2, we observed that k_1_ and k_2_ values were rather constant among the different HT7 fluorophore substrates, while larger differences were observed for the dissociation rate constant k_−1_ (**Fig. S6 & Table S1**). The substrate preference of HT7 seems therefore mainly driven by the substrate affinity 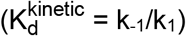 (**Fig. 2B**). After binding, the deeply buried CA moiety might adapt a similar conformation for all substrates, potentially explaining the minor effects of the substituent on the catalytic step (k_2_) (**Fig. 2C**). The trend observed for the K_d_ values calculated from the kinetic parameters was confirmed by measuring the affinity of the catalytically dead variant HT7^D106A^ for the same CA-fluorophore substrates using fluorescence polarization (**Fig. S6F & S7**). The 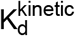 correlates with 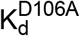 (**Fig. S6E**) and as a consequence the association constant 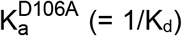 correlates with k_app_ (**Fig. 2F**). Hence, the 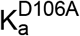 can be used to estimate the k_app_ for fluorescent HT7 substrates.

### HaloTag7 reacts slower with non-fluorophore substrates

In order to determine k_app_ for non-fluorescent CA substrates, we developed a competitive kinetic assay in which the non-fluorescent CA substrates compete with CA-Alexa488 for protein labeling. Non-fluorescent substrates were significantly slower than zwitterionic rhodamine substrates (10^3^ < k_app_ < 10^6^ M^−1^s^−1^), highlighting the strong preference of HT7 for the rhodamine core structure. Larger alkynes (*e.g*. SCO, BCN) and aromatic structures (*e.g*. Tz, PhN3, VBn) were preferred over alkenes (Nor) and small moieties (Ac, N_3_) (**Fig. 2D, S8 & Table 1**).

### HaloTag7 substrate design

Overall, HT7 can reach labeling kinetics near the diffusion limit but its apparent rate constants span over six orders of magnitude, depending on the nature of the label (**Fig. 2D**). HT7 exhibits a strong preference for rhodamine derivatives, with the exception of negatively charged rhodamines. It is noteworthy that the substrate with the slowest labeling rate carries the smallest label, *i.e*. an acetate group (CA-Ac). The preference for rhodamines can be exploited to increase labeling rates of poor substrates. As an example, the commercially available CA-PEG-biotin substrate presents slow reaction kinetics (k_app_ = 1.70 ± 0.08 × 10^4^ M^−1^s^−1^, **Table 1 & Fig. S8**), but synthesizing a CA-TMR-biotin ligand led to an over 500 fold increase in labeling kinetics (k_app_ = 1.04 ± 0.01 ×10^7^ M^−1^s^−1^, **Table 1, S1 & Fig. S2**), greatly facilitating biotinylation of HT7 fusion proteins. This strategy to improve labeling rates of HT7 ligands should be applicable to various other labels.

### Structural analysis of rhodamine-bound HaloTag

In order to better understand the substrate preference of HT7 for rhodamine-based CA substrates, we solved the X-ray structure of TMR- (PDB ID 6Y7A) and CPY-bound HT7 (PDB ID 6Y7B) at 1.4 Å and 3.1 Å resolution, respectively (**Fig. 3A, 3C, S9 & Table S4**). Additionally, the TMR-bound structure of HOB was obtained at 1.5 Å resolution (PDB ID 6ZCC) (**Fig. 3H, S9 & Table S4**). These structures present the same α/β hydrolase fold of the superfamily with minimal deviation from already available HT7 X-ray structures (19–22) (**Fig. S9C**). In addition to the conventional α/β hydrolase topology, HT7 features an extra capping domain made of six α-helices (Hlx4 to 9) which partially cover the catalytic site and form an entry channel for the CA substrate. After reaction, the PEG-alkane ligand is buried in the protein, while the xanthene moiety of the dye lays on the distorted α-helix 8 (Hlx8) in a conformation partially constrained by the crystal packing (**Fig. 3A, 3C & S9D**). A recently published HT7-TMR X-ray structure (PDB ID 6U32) shows the fluorophore bound in two alternative conformations (23). In one conformation, the fluorophore lays on Hlx8 similar to what we report here and in the other, it lays on the Hlx7-turn-Hlx8 motif (**Fig. S10**). This second conformation is incompatible with our HT7-TMR structure due to steric clashes caused by the crystal packing. The alkane-fluoro-phore is positioned by the Hlx6-turn-Hlx7-turn-Hlx8 motif of the HT7 capping domain from which T172^Hlx8^ and, to a lesser extent, T148^Hlx6^ form hydrogen bonds with the oxygen and the nitrogen of the amide bond linking PEG-alkane and fluorophore (**Fig. 3A & 3C**). CA-TMR and CA-CPY have similar conformations in both structures (**Fig. S11A**) with only minor differences in their torsion angles (**Fig. 3E**). In comparison to TMR, one of the additional methyl groups of CPY is forming van-der-Waals interactions at the protein surface, potentially explaining the increased affinity of CA-CPY relative to CA-TMR.

**Figure 3:**
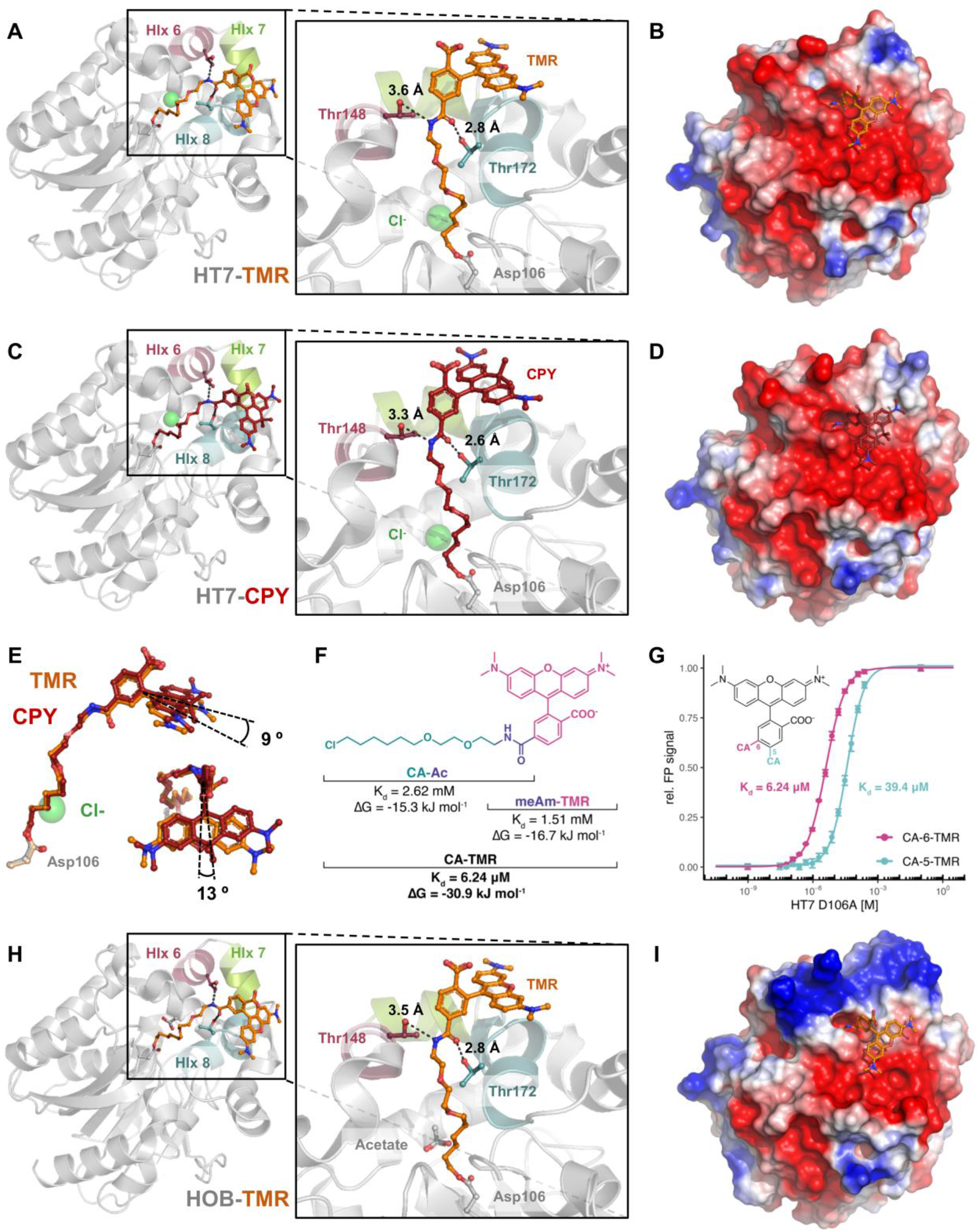
Structure-function analysis of HaloTag7 substrate interactions. Crystal structures of HT7-TMR (PDB ID 6Y7A, **A**), HT7-CPY (PDB ID 6Y7B, chain A, **C**) and HOB-TMR (PDB ID 6ZCC, **H**). Proteins are represented as grey cartoons, the fluorophore substrates and residues as sticks. Putative hydrogen bonds are represented as black dashed lines with annotated distances. Electrostatic potentials at protein surfaces (**B, D** & **I**, respectively) are drawn at −2.0 (red) to 2.0 (blue) kJ/mol/e and were obtained using the APBS software with standard parameters. **E**. Comparison of the TMR and CPY conformation on HT7. **F**. HT7 affinities (K_d_) and free binding energies (ΔG) for different TMR substrate substructures. **G**. Comparison of HT7 affinity for CA-6-TMR and CA-5-TMR.

### Fluorophore and CA core contribute both to HaloTag7 substrate affinity

To characterize the contributions of rhodamine structures and the CA core to the overall affinity of HT7 substrates, we measured affinities of the catalytically dead variant HT7^D106A^ for the acetylated chloroalkane (CA-Ac) and *N*-methylamide-fluorophores (meAm-TMR/CPY). Although the acetylated chloroalkane should form hydrogen bonds to the protein (via T148/T172) and is well buried in the cavity, we observed a rather low affinity (K_d_) of 2.62 mM (2.44 to 2.72 mM CI 95%, **Fig. 3F & S12**), which is consistent with the low apparent labeling rate constant of CA-Ac (**Fig. 2D**). The protein binds the meAm-TMR fluorophore with a slightly higher affinity (K_d_ = 1.51 mM, 1.40 to 1.64 mM CI 95%) (**Fig. 3F & S12**). The free binding energies for both fragments calculated from the K_d_ values (CA-Ac: −15.3 kJ.mol^−1^ and meAm-TMR: −16.7 kJ.mol^−1^) are thus comparable and almost sum up to the calculated free binding energy of the full CA-TMR substrate (30.9 kJ mol^−1^, K_d_ = 6.24 μM), *i.e*. no synergistic effect in binding is observed (24). Similar results were obtained for meAm-CPY (**Fig. S12**). The CA-fluorophore binding is thus driven by interactions with both the CA core and the fluorophore, explaining the high impact of fluorophore structure changes on the overall labeling kinetics.

The importance of substrate geometry was interrogated by synthesizing CA-fluorophore substrates linked via the 5 position of the rhodamine benzyl ring instead of the usual 6 position (**Fig. 3G**). According to the observed conformations in the presented crystal structures, these 5-substrates should not be able to interact with Hlx8 after HT7 binding since the xanthene would be turned 60° away from the protein surface. HT7^D106A^ showed reduced affinities towards these substrates compared to the 6-substituted rhodamine substrates (6.31 fold and 22.7 fold decrease for CA-TMR and CA-CPY, respectively) (**Fig. 3G**). This result emphasizes the importance of the interaction between the xanthene ring and the Hlx8.

### HaloTag7 surface charge impacts substrate recognition

HOB comprises four surface mutations compared to HT7 close to the substrate entry channel but opposite to the TMR binding site (**Fig. S11B**). These mutations lead to faster labeling rates with negatively charged CA substrates relative to HT7. Only minor differences can be observed between the crystal structures of HOB and HT7 labeled with CA-TMR (**Fig. 3H & S11B**). Since the HOB mutations replace mostly negative by positively charged residues, we analyzed the electrostatic potential of both proteins. While HT7 features an overall negatively charged surface around the substrate entry channel (**Fig. 3B & 3D**), HOB shows a positively charged patch opposite to the fluorophore binding site (**Fig. 3I**). Hence, a putative electrostatic steering effect (25) could explain the altered substrate preference of HOB despite that its positive charges are on the opposite side of the fluorophore binding site.

### Kinetic characterization of SNAP-tag

SNAP labeling kinetics were characterized for both BG- and CP-fluorophore substrates (*i.e*. TMR, CPY, Alexa488 and Fluorescein) (**Fig. 1D & S1**), by following fluorescence polarization changes during the labeling reaction at different protein concentrations in a plate reader assay. The kinetic model 2 did not allow to determine the kinetic parameters (k_1_, k_−1_ & k_2_) independently. Hence, data were fitted to model 1 in order to obtain apparent second order rate constants (k_app_) of the labeling reactions (**Table 1 & Fig. S13**). SNAP’s apparent labeling rate constants are ranging between 10^4^ and 10^6^ M^−1^s^−1^ for BG-fluorophore substrates (**Fig. 4A**), among which BG-TMR presents the fastest labeling rate (kapp = 4.29 ± 0.01 × 10^5^ M^−1^s^−1^) (**Table 1**). CP substrates show 4 – 14 times slower reaction kinetics than the corresponding BG substrates (10^3^ < k_app_ < 10^5^ M^−1^s^−1^) (**Fig. 4A**). Some CP substrates (CPY and Fluorescein) exhibit a slow additional phase of fluorescence polarization increase or decrease after labeling that might be due to a slow conformational change of the labeled protein. In order to fit these traces, the kinetic model 1 was extended by adding a step that occurs after labeling. The rate constants of this additional process (k_3_) ranged between 10^−2^ and 10^−3^ s^−1^ (**Fig. S13 & Table S5**). SNAP labeling with BG-TMR and CP-TMR was further investigated by measuring stopped flow fluorescence anisotropy kinetics at higher protein concentrations (**Fig. S14 & Table S6**). Fitting the data to the kinetic model 2 allowed to estimate the kinetic parameters k_1_, k_−1_ and k_2_ independently and to calculate K_d_ values (**Fig. S14C**). The calculated k_app_ for both substrates were similar to the k_app_ determined via the plate reader assay using model 1 (**Fig. S14C**). CP-TMR presents similar k_1_ and k_2_ as BG-TMR, while k_−1_ is significantly higher for CP-TMR (8.8 fold), indicating that both substrates feature the same chemical reactivity but differ in their affinity towards SNAP.

**Figure 4:**
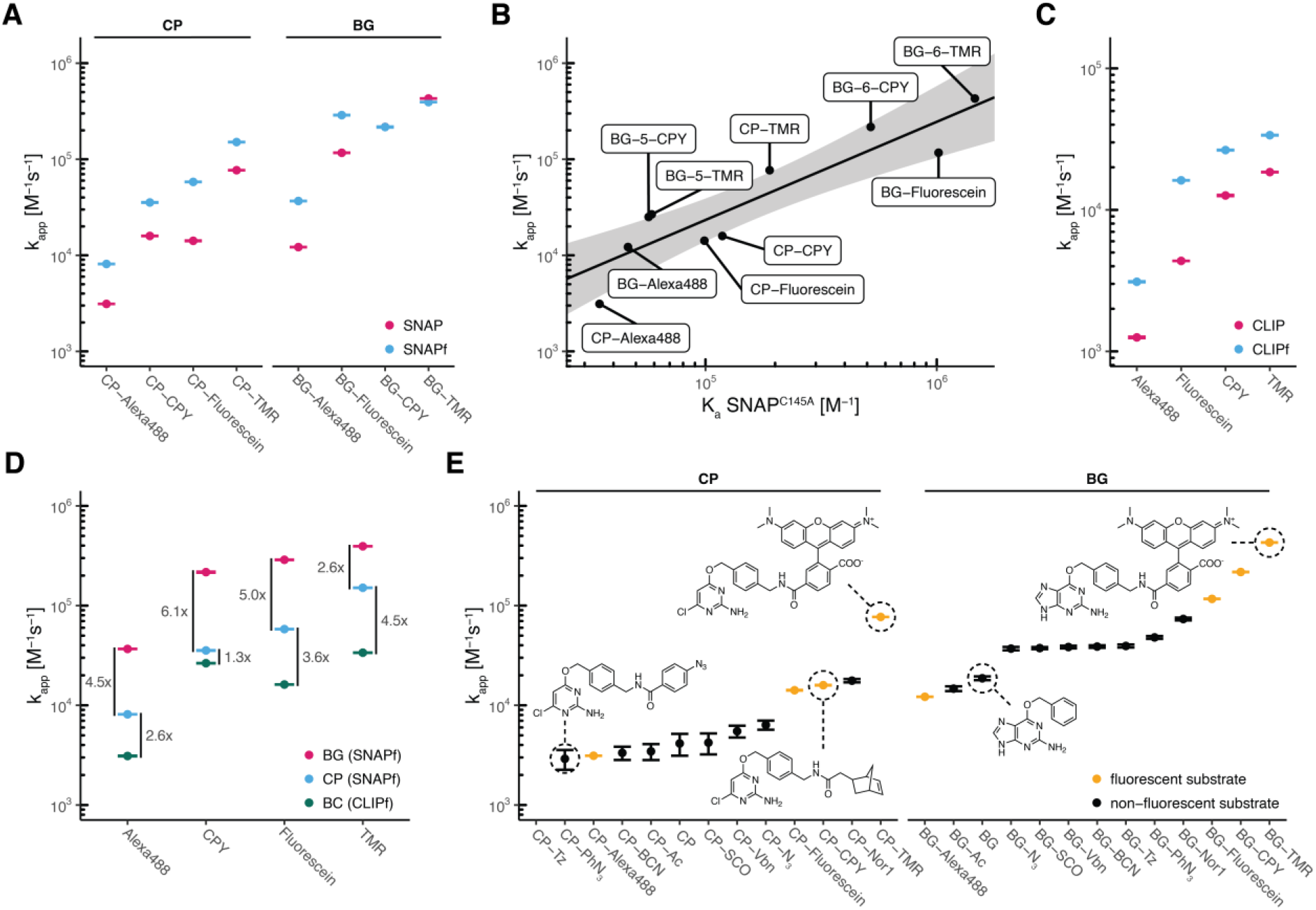
Characterization of SNAP- and CLIP-tag labeling kinetics. **A**. Comparison of labeling kinetics (k_app_) between SNAP and SNAPf. **B**. Correlation between SNAP apparent second order rate constant (k_app_) and affinity (K_a_ = 1/K_d_) for different fluorophore substrates. Affinities were obtained for the catalytically inactive variant SNAP^C145A^. Log transformed values were fitted to a linear model (black line, log(k_app_) = log(K_a_) * 1.0217 - 0.7407). The grey area represents the 95% confidence bands (the area in which the true regression line lies with 95% confidence). **C**. Comparison of labeling kinetics (k_app_) between CLIP and CLIPf. **D**. Comparison of labeling kinetics (k_app_) between SNAPf and CLIPf. **E**. Apparent second order labeling rate constants (k_app_) of SNAP with different substrates. Kinetics span over three orders of magnitude (two orders of magnitude within each substrate class BG/CP). BG-based, non-negatively charged fluorophore substrates reach the fastest labeling kinetics.

### SNAP-tag labeling kinetics correlate with substrate affinity

To confirm the previous finding, affinities for different fluorescent substrates were measured using the catalytically inactive mutant SNAP^C145A^ (**Fig. S15**). A strong preference for BG-TMR over CP-TMR was observed with almost one order of magnitude difference in 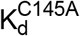. SNAP^C145A^ presents a 3 fold lower 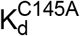 for BG-TMR (0.68 μM; 0.63 to 0.75 μM CI 95%) than calculated from stopped-flow experiments using active SNAP. SNAP^C145A^ showed similar affinities as for BG-TMR towards various xanthene-based fluorophores such as BG-MaP555, BG-JF549 and BG-fluorescein (**Fig. S15**), indicating that modifications of the rhodamine structure seem not to affect the affinity of the protein as much as observed for HT7 substrates. However, SNAP^C145A^ has very low affinity for sulfonated fluorophore substrates such as BG-Alexa488 (21.6 μM; 20.5 to 22.9 μM CI 95%) or BG-sulfo-Cy3/5 (Cy3, 68.1 μM; 63.8 to 72.7 μM CI 95%) (**Fig. S15**). A good correlation between 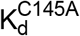 and k_app_ was observed for the tested fluorophore substrates (**Fig. 4B**), highlighting again the importance of high affinity for a quick labeling reaction. As for HT7, we attempt to decipher SNAP substrate recognition by measuring its affinity towards BG-Ac and meAm-TMR. While no affinity could be measured for meAm-TMR, SNAP^C145A^ presented a relatively high affinity for BG-Ac (88.0 μM; 88.6 to 91.5 μM CI 95%) and CP-Ac (201 μM; 192 to 212 μM CI 95%) compared to HT7 affinity for CA-Ac (**Fig. S16**), which could explain the promiscuity of SNAP.

### Kinetic characterization of CLIP-tag and SNAP-tag variants

The mutant SNAPf (SNAP^E30R^) is a SNAP variant with faster labeling rates for BG-Alexa488, BG-TMR, BG-Atto549 and BG-AlexaFluor647 (26) (**Fig. 4A**, **Fig. S17**). Fluorescence polarization kinetics of SNAPf revealed a 2 to 4 fold k_app_ increase compared to SNAP for most BG- and CP-fluorophore substrates (**Fig. 4A, S7, S18 & Table S7**). Nevertheless, no increase in labeling kinetics was observed for the best SNAP substrates BG-TMR and BG-CPY (**Fig. 4A**). CLIP (11) and CLIPf (CLIP^E30R^) (26) are or-thogonal variants of SNAP accepting BC instead of BG substrates (**Fig. S17**). Labeling kinetics of CLIP and CLIPf (**Table S7 & Fig. S19**) yielded apparent second order rate constants (kapp) ranging from 10^3^ to 10^5^ M^−1^s^−1^ with a 2 to 4 fold increase for CLIPf compared to CLIP (**Fig. 4C**). The fastest labeling kinetics were achieved with CLIPf and BC-TMR showing a k_app_ of 3.37 ± 0.01 × 10^4^ M^−1^s^−1^. However, CLIPf is significantly slower than SNAPf (**Fig. 4D**).

### Cross-reactivity of SNAP- and CLIP-tag substrates

SNAP and CLIP originate from hAGT (10, 11) (**Fig. S17**), which can potentially react with SNAP and CLIP substrates. We therefore measured the labeling activity of hAGT for the corresponding TMR-based substrates (**Fig. S20**). BG/CP-TMR labeling of hAGT is 130 / 20 times slower than the labeling of SNAP (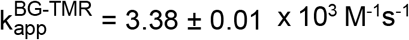; 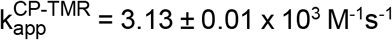) (**Table 2**). Interestingly, hAGT shows no preference for BG over CP substrates. BC-TMR reaction with hAGT is 25’000 times slower than with CLIP (k_app_ = 0.70 ± 0.01 M^−1^s^−1^) (**Table 2**). Our results suggest that CLIP should be preferred over SNAP in cases where cross-reactivity of substrates with endogenous hAGT is a concern.

**Table 2:** Labeling kinetics (k_app_) of hAGT, SNAP and CLIP with TMR substrates.

| $k_{app} [\text{M}^{-1}\text{s}^{-1}]$ (value \| s.d.) | | | |
| --- | --- | --- | --- |
|  | hAGT | SNAP | CLIP |
| BG-TMR | $3.38 (\pm 0.01) \times 10^3$ | $4.29 (\pm 0.01) \times 10^5$ | $8.26 (\pm 0.05) \times 10^1$ |
| CP-TMR | $3.13 (\pm 0.01) \times 10^3$ | $7.69 (\pm 0.01) \times 10^4$ | $7.22 (\pm 0.04) \times 10^0$ |
| BC-TMR | $6.25 (\pm 0.01) \times 10^{-1}$ | $3.20 (\pm 0.02) \times 10^2$ | $1.85 (\pm 0.01) \times 10^4$ |

CLIP development was motivated by the perspective to use both SLPs together for multicolor labeling. However, the cross-reactivities of the fastest reacting SNAP and CLIP rhodamine sub-strates have not yet been determined. Hence, we measured cross-reactivity of BG/CP-TMR with CLIP and BC-TMR with SNAP (**Table 2**). SNAP reacts more than 1000 times slower with BC-TMR (SNAP 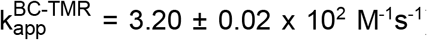) than with BG-TMR despite the noticeable affinity of SNAP^C145A^ for BC-Ac (416 μM, 408 to 421 μM CI 95%) which is only 5 times lower than for BG-Ac (**Fig. S16**). On the other hand, CLIP reacts 100 times slower with BG-TMR (CLIP 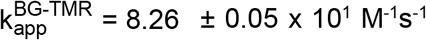) than with BC-TMR. These data are in agreement with values previously reported for fluorescein substrates (11). Since both proteins show residual reactivity towards their non-respective substrates, simultaneous co-labeling of both proteins or prior SNAP labeling is advisable to minimize cross-reactions.

### SNAP-tag is a promiscuous SLP

Labeling kinetics of non-fluorescent SNAP substrates were characterized by competition kinetics against BG-Alexa488 (**Fig. S21**). Non-fluorescent BG substrates (10^4^ < k_app_ < 10^5^ M^−1^s^−1^) were preferred over CP substrates (10^3^ < k_app_ < 10^4^ M^−1^s^−1^) (**Fig. 4E & Table 1**). In general, SNAP kinetics with non-fluorescent substrates were slower than with fluorescent substrates with the exception of the negatively charged Alexa488. However, in comparison to HT7, the labeling rates of SNAP show much less dependence on the nature of the label (**Fig. 4E & Table 1)**.

### Structural analysis of TMR-bound SNAP-tag

To better understand the preference of SNAP for TMR substrates, the X-ray structure of SNAP labeled with TMR was solved at 2.3 Å resolution (PDB ID 6Y8P) (**Fig. 5A, S22 & Table S4**). The structure shows the same α/β topology with two domains as observed for hAGT and other SNAP structures (27, 28). The active site is very similar to the benzylated SNAP structure (PDB ID 3L00) (28), despite the presence of an alternative cysteine conformation (**Fig. S22C**). The TMR moiety strongly participates in the crystal packing, engaging in interactions with the neighboring xanthene ring and protein in a sandwich-like topology (**Fig. S22D**). As a consequence, and in contrast to HT7-TMR, SNAP does not interact with the bound fluorophore in the present X-ray structure.

We next evaluated the relative preference for 6-versus 5-carboxy isomers of TMR and CPY substrates by studying their labeling rates (**Fig. S23 & Table S8**) and affinities (**Fig. S15**) for SNAP, SNAPf and their dead variants. SNAP and SNAPf showed 10 times slower reaction rates with 5-fluorophores (k_app_ ≈ 10^4^ – 10^5^ M^−1^s^−1^) compared to the corresponding 6-fluorophores (k_app_ ≥ 10^5^ M^−1^s^−1^). These differences were even more pronounced for the affinities, which were up to 25 fold higher for the 6-carboxy isomers.

In the crystal structure of TMR-labeled SNAP, a structural ethylene glycol forms hydrogen bonds with both the backbone carbonyl oxygen of I31 and the carbonyl oxygen of the amide linking the benzyl to the fluorophore (**Fig. 5A**). This benzyl-fluorophore amide is also forming a hydrogen bond to the backbone carbonyl oxygen of the catalytically important E159 residue via its Nα atom. Comparison with the BG-bound SNAP^C145A^ structure (PDB ID 3KZZ, **Fig. 5B**) suggests that, after catalytic reaction, the E159 side chain flips inside the BG binding cavity, resulting in a reorientation of its backbone carbonyl oxygen that can then interact with the amide of the substrate (**Fig. 5B**).

**Figure 5:**
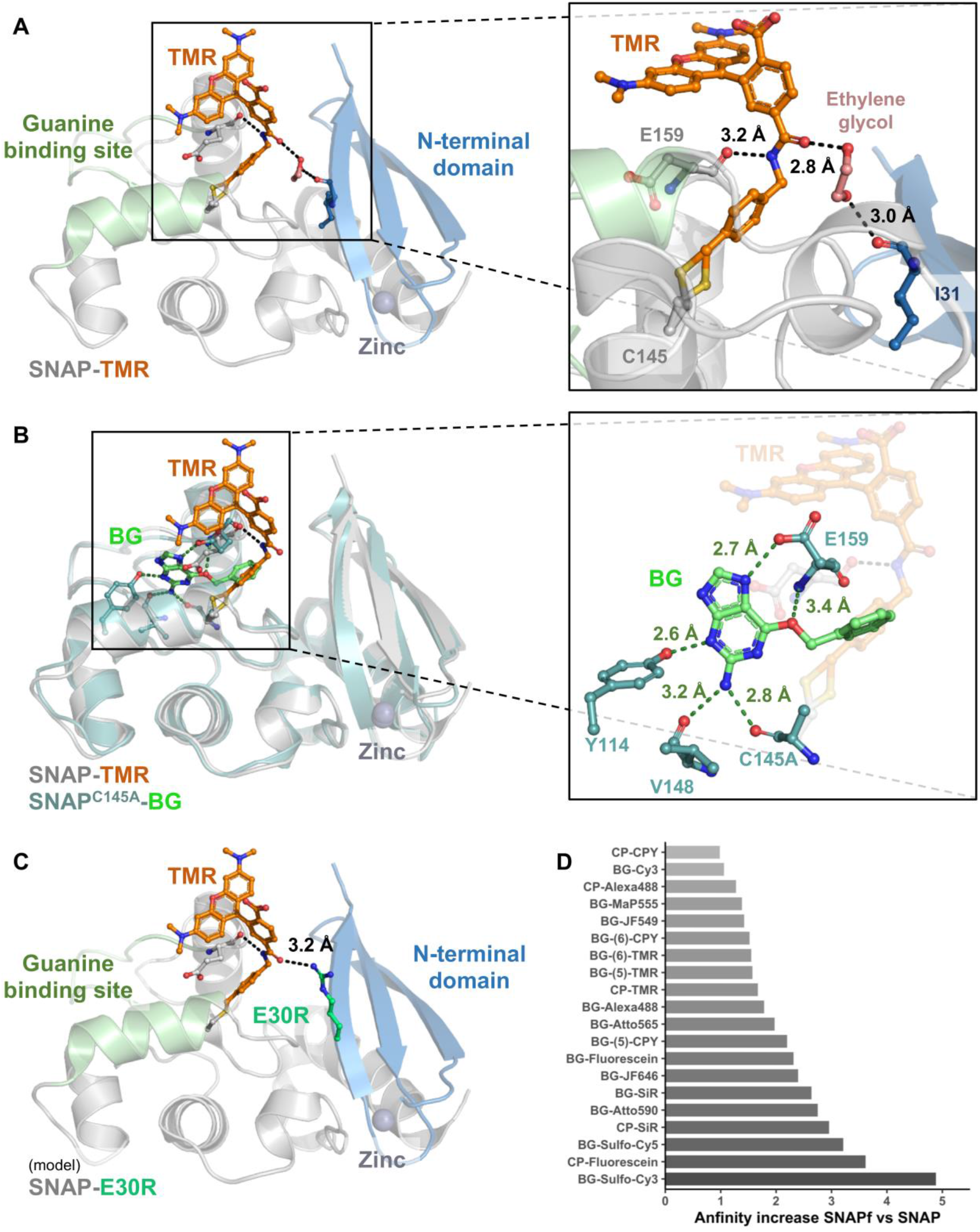
Structure-function analysis of SNAP-tag fluorophore substrate interactions. **A**. Crystal structure of SNAP labeled with a TMR substrate. **B**. Structural comparison between SNAP-TMR and the BG bound variant of SNAP^C145A^. **C**. Modeling of the E30R mutation in the SNAP-TMR crystal structure. SNAP is represented as cartoon, the fluorophore substrate and residues as sticks. Putative hydrogen bonds and corresponding distances are indicated by black dashes. **D**. Affinity increase between SNAP^C145A^ and SNAPfC^145A^ for different fluorophore substrates. Number in brackets indicate different linkage of the fluorophore benzyl group to BG.

### SNAPf has a higher affinity for its substrates

We modeled the SNAPf mutation E30R in the structure of TMR-labeled SNAP to gain a better understanding of how it affects the labeling kinetics (**Fig. 5C**). Results suggest that an arginine in position 30 could interact with the carbonyl oxygen of the amide group in the label via a moderate hydrogen bond (3.2 Å), replacing the hydrogen bond observed with the ethylene glycol in the crystal structure. This could lead to an increased affinity for the substrate or a better substrate positioning resulting in a quicker labeling. To probe this hypothesis, the affinities of SNAP^C145A^ and SNAPfC^145A^ were compared side by side for various fluorophore substrates (**Fig S16**). Among the 21 fluorophore substrates tested, only five did not show a significant increase in affinity (*i.e*. above 50%) and nine showed more than a 2 fold affinity increase (**Fig. 5D**). As observed for SNAP, SNAPf^C145A^ substrate affinities correlate well with the corresponding k_app_ values for SNAPf (**Fig. S24**). It is noteworthy to mention that negatively charged substrates such as BG-sulfo-Cy3 show the strongest increase in the protein affinities and labeling rates when comparing SNAP to SNAPf. This could be due to the exchange of the negatively charged glutamic acid by a positively charged arginine resulting in a potential electrostatic steering effect as mentioned for HT7 (25).

### Comparison between SNAP-tag and SsOGT-H^5^

Recently, an homologue of hAGT from an extremophile archaea was converted to an SLP (*Ss*OGT-H^5^) by introducing mutations that have been shown to increase the reactivity of SNAP (29). Its crystal structure labeled with SNAP-Vista Green^®^ (SVG, *i.e*. BG-5-fluorescein) (30) shows a different fluorophore conformation, constrained by the crystal packing (**Fig. S25**). Interestingly, the *Ss*OGT-H^5^-SVG structure was obtained with a fluorophore connected via the 5-carboxy isomer of the fluorophore and presents a substrate conformation that could not exist in the SNAP structure due to steric clashes (**Fig. S25A**). We compared the kinetics of SNAP and *Ss*OGT-H^5^ (**Fig. S26 & Table S9**) toward the substrates BG-TMR (5- and 6-substituted) and BG-6-Alexa488 at 37°C. In contrast to SNAP, *Ss*OGT-H^5^ showed a preference for BG-5-TMR (k_app_ = 1.45 ± 0.92 x 10^2^ M^−1^s^−1^) over BG-6-TMR (k_app_ = 6.78 ± 0.67 x 10^1^ M^−1^s^−1^). Furthermore, the negatively charged BG-6-Alexa488 (kapp = 1.24 ± 0.01 x 10^2^ M^−1^s^−1^) presents kinetics in the same range as BG-5-TMR, highlighting a different substrate preference between SNAP and *Ss*OGT-H^5^. For all substrates, *Ss*OGT-H^5^ presents kinetics 100 times slower than SNAP or CLIP, making it less suitable for labeling applications at physiological temperatures.

## Discussion

We provide here a systematic comparison of the labeling kinetics of HT7, SNAP and CLIP towards a large panel of substrates. A structure-function relationship analysis complements this comparison, thereby yielding insights into the origins of the different substrate specificities of HT7 and SNAP. The data should assist scientists in choosing SLP-substrate pairs for specific purposes.

**Figure 6:**
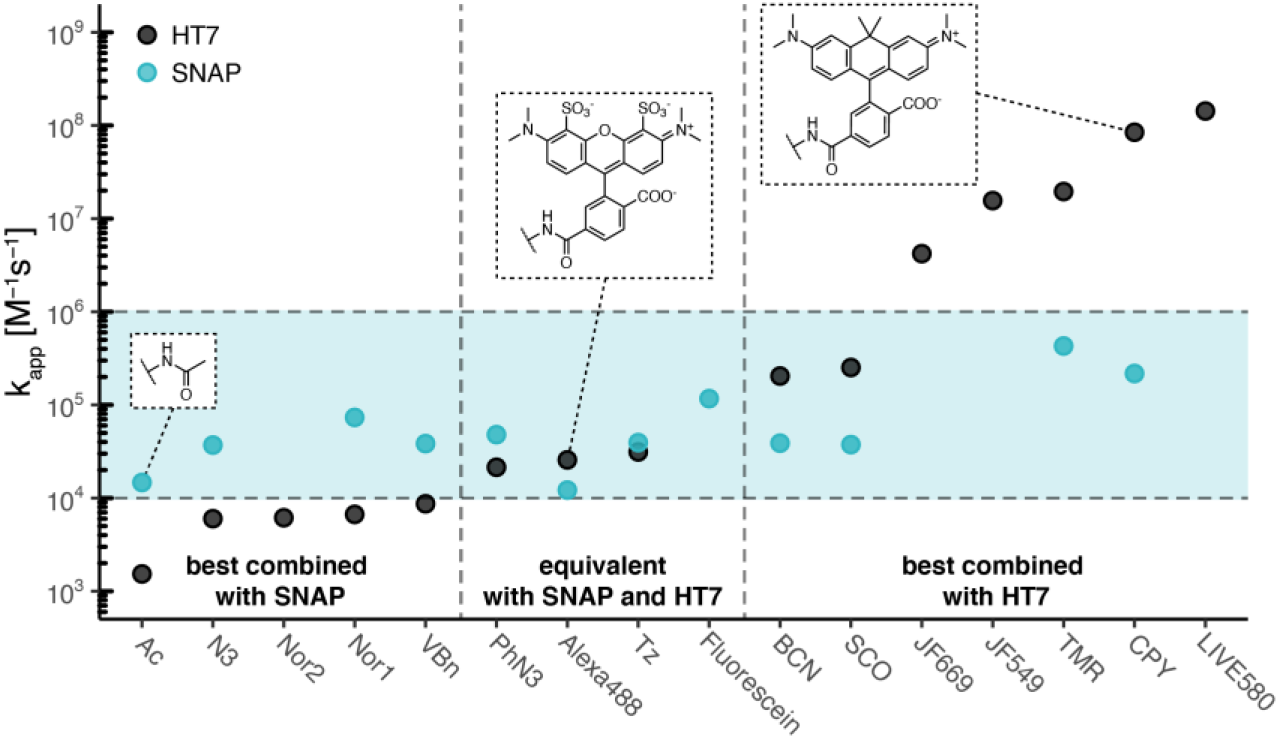
Labeling kinetics comparison between SNAP-tag and HaloTag7. Apparent labeling rate constants (kapp) of HT7 span over six orders of magnitude while rate constants of SNAP span only over two orders of magnitude (BG-substrates). The blue area highlights the span of SNAP apparent labeling rate constants. Depending on the application, some substrates should preferentially be employed with HT7 or SNAP to ensure quick labeling.

The direct comparison of SNAP and HT7 reveals that HT7 features significantly higher labeling rates with various fluorescent rhodamine derivatives (**Fig 6 & Table 1**). These differences in reactivity can be explained by specific interactions of the rhodamine’s xanthene ring with selected surface residues of HT7. The high reactivity of HT7 towards rhodamines is important as rhodamines up to now represent the most relevant class of cell-permeable fluorophores for live-cell imaging. The interactions between rhodamines and HT7 also help to explain why some rhoda-mine-based HT7 substrates tend to have improved spectroscopic properties and are more fluorogenic than the corresponding SNAP or CLIP substrates (16). Most rhodamine-based fluorophores exist in an equilibrium between spirocyclic non-fluorescent and zwitterionic fluorescent forms. While in solution the spirocyclic form might be favored, labeling reaction with an SLP switches this equilibrium toward the zwitterionic form, leading to a fluorescence intensity increase (31). This property is of particular interest in wash-free live-cell fluorescence microscopy since it leads to higher signal over background (26, 32–35) and can also be exploited for sensor design (23, 36). Furthermore, the dynamic equilibrium between the spirocyclic non-fluorescent and zwitterionic fluorescent form is crucial for cell permeability (33). The mechanism underlying the equilibrium shift from the spirocyclic non-fluorescent to the zwitterionic fluorescent form is not fully understood yet but our results indicate that the planar, zwitterionic form of rhodamines (*e.g*. TMR and CPY) features energetically favorable interactions with HT7 surface, thus potentially favoring this state of the fluorophore when labeled to the protein.

While HT7 reacts quicker with most rhodamine-based fluorophore substrates than SNAP, the dif-ferences become much less pronounced or reversed for negatively charged substrates. For ex-ample, SNAP reacts faster with Alexa488 than HT7 and the reactivity for most other non-fluorescent substrates tends to be higher for SNAP as well (**Fig 6 & Table 1**). It is interesting to hypothesize about the origin of the substrate specificity differences between SNAP and HT7. Most likely, these differences are, at a least partially, a consequence of the substrates used in the engineering of the tags. For HT7, TMR was used in most screening assays (9, 13) and, as a result, HT7 shows a specificity for zwitterionic rhodamines. In contrast, different substrates such as BG-fluorescein (37), BG-Cy3 (38) as well as affinity reagents such as BG-biotin (37) were used in SNAP screening and selection assays. As a consequence, SNAP is more promiscuous than HT7. Differences in labeling speed of both SLPs are mostly driven by differences in substrate affinity: an overall correlation between affinity and rate constants was observed for both proteins that was more pronounced for HT7. Indeed, HT7 presents a very low affinity toward the *e.g*. unsubstituted CA-Ac substrate highlighting that HT7 affinity toward substrates is highly driven by the substituent and so are the kinetics. We show here how the low reactivity of HT7, for example towards CA-PEG-biotin, can be overcome by designing substrates in which the label of interest is attached to a CA-TMR core and anticipate that such strategy could be expanded to other substituents.

A key property of SLP substrates for live-cell applications that we have not addressed in this study is their cell permeability. Generally speaking, the CA core is less polar than BG, CP and BC. The permeability of HT7 substrates therefore can be expected to be higher than the corresponding SNAP-tag substrates. However, this question will have to be more systematically addressed in future studies.

For future engineering of SLPs, it would be particularly interesting to increase the affinity of SNAP and CLIP towards rhodamine-based substrates. Given the importance of these fluorophores for live-cell fluorescence (super-resolution) microscopy (1), additional tags that display labeling kinetics towards rhodamines similar to those of HT7 would be highly welcomed. Our results suggest that increasing the reactivity towards these dyes might come with the risk of reducing the activity towards other substrates, thereby limiting the flexibility of such tags. However, given the importance of SLPs and rhodamine-based probes for live-cell imaging, the generation of such specialized tags is warranted.

## Materials and Methods

### Labeling substrates and chemical synthesis

Labeling substrates for HaloTag, SNAP-tag and CLIP-tag were synthesized according to literature procedures (10, 11, 15, 32–34, 39–45); purchased from Promega Corp. (Madison, WI, USA), Abberior GmbH (Göttingen, Germany), Santa Cruz Biotechnology Inc. (Dallas, TX, USA) and NEB Inc. (Ipswitch, MA, USA); were kind gifts from Dr. L. Lavis (Janelia research campus, USA) and Dr. A.D.N. Butkevich (MPI for Medical Research, Germany) or were synthesized according to the procedure available in the supplementary information.

### Cloning, protein expression and purification

SNAP, SNAPf, SNAP^cx^, CLIP, CLIPf, HT7 and HOB were cloned in a pET51b(+) vector (Novagen) for production in *Escherichia coli*, featuring an N-terminal His_10_ tag and a Tobacco Etch Virus (TEV) cleavage site. *Ss*OGT-H^5^ and hAGT were cloned in the same plasmid featuring an N-terminal StrepTag-II and an enterokinase cleavage site together with a C-terminal His_10_ tag. Cloning was performed by Gibson assembly (46) using E.cloni 10G cells (Lucigen) and point mutations were performed using the Q5 site-directed mutagenesis kit (NEB). Proteins were expressed in *E. coli* strain BL21(DE3)-pLysS (Novagen). Lysogeny broth (LB) (47) cultures were grown at 37°C to optical density at 600 nm (OD_600nm_) of 0.8. Transgene expression was induced by the addition of 0.5 mM isopropyl-β-D-thiogalactopyranoside (IPTG) and cells were grown at 17°C overnight in the presence of 1 mM MgCl2. Cells were harvested by centrifugation and lysed by sonication.

For N-terminally His-tagged proteins, the cell lysate was cleared by centrifugation (75 000g, 4° C, 10 min) before affinity-tag purification using a HisTrap FF crude column (Cytiva, Marlborough, MA, USA) and an ÄktaPure FPLC (Cytiva). Buffer was exchanged using a HiPrep 26/10 Desalting column (Cytiva) to HEPES 50 mM, NaCl 50 mM pH 7.3 (*i.e*. activity buffer). Proteins were concentrated using Ultra-15 mL centrifugal filter devices (Amicon, Merck KGaA, Darmstadt, Germany) with a molecular weight cut-off (MWCO) smaller than the protein size to a final concentration of 500 μM. Proteins were aliquoted and stored at −80°C after flash freezing in liquid nitrogen. Double-tagged proteins, after similar cell lysis and clearing, were purified using HisPur Ni-NTA Superflow Agarose (Thermo Fisher Scientific, Waltham, MA, USA) by batch incubation followed by washing and elution steps on a polypropylene column (Qiagen). Proteins were subsequently purified using a StrepTrap HP column (Cytiva) on an ÄktaPure FPLC. Proteins were then concentrated using Ultra-5 mL centrifugal filter devices with a MWCO smaller than the protein size and conserved in glycerol 45 % (w/v) at −20°C.

Correct size and purity of proteins were assessed by SDS-PAGE and liquid chromatography-mass spectrometry (LC-MS) analysis.

### Affinity of HT7 and HOB towards CA substrates

Binding affinities of HT7^D106A^ or HOB^D106A^ to chloroalkane (CA) substrates were determined by fluorescence polarization (FP, equation **1**) measurements using a microplate reader (Spark20M^®^, Tecan Group AG, Männedorf, Switzerland). The fluorescent substrates (10 nM) were titrated against different protein concentrations (0 – 250 μM) in activity buffer supplemented with 0.5 g/L BSA. Assays were performed in black low-volume non-binding 384-well plates (Corning Inc., Corning, NY, USA) with a final volume of 20 μL. All measurements were performed in triplicates at 37°C, filter settings are listed in **Table 3**. Obtained FP values were averaged and fitted to a single site binding model (equation **2**) to estimate K_d_ values for each fluorescent substrate. The FP value of each dye fully reacted with the native HT7 was used to improve fitting of the curves upper plateau by adding an extra data point at protein concentration of 0.1 M.

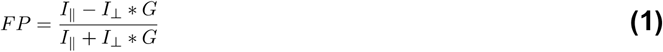

with FP: fluorescence polarization, I_║_: fluorescence intensity parallel to the excitation light polarization, I_⊥_: fluorescence intensity perpendicular to the excitation light polarization and G: grating factor (*G* = *I*_║_/*I*_⊥_).

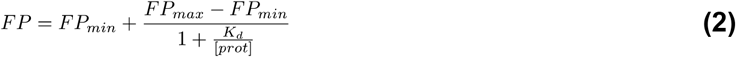

xs
with FP_min_: fluorescence polarization of the free fluorophore (lower plateau), FP_max_: maximal fluorescence polarization of fully bound fluorophore (upper plateau), K_d_: dissociation constant and [prot] = protein concentration.

**Table 3:** Filter settings used in FP measurements.

| Fluorophore | Excitation filter (BW) | Emission filter (BW) |
| --- | --- | --- |
| Alexa488, Fluorescein, Oregon green, JF503, 500R | 485 (20) nm | 535 (25) nm |
| TMR, JF549, JF525, TMR-az-F2, TMR-CN, TMR-SCH3, TMR-SNH2, MaP555, 510R, 515R, 580CP, Atto565, Atto590, (sulfo-)Cy3 | 535 (25) nm | 595 (35) nm |
| CPY, SiR, LIVE580, JF608, JF646, JF669, (sulfo-)Cy5 | 620 (20) nm | 680 (30) nm |

### Affinity of SNAP and SNAPf towards BG and CP substrates

Binding affinities of SNAP^C145A^ and SNAPf^C145A^ toward BG-Alexa488, CP-Alexa488, BG-Fluorescein, CP-Fluorescein, BG-MAP555, BG-JF549, BG-TMR(6), BG-TMR(5), CP-TMR, BG-CPY(6), BG-CPY(5), CP-CPY, BG-SiR, CP-SiR, BG-JF646, BG-Atto565, BG-Atto590, BG-sulfo-Cy3, BG-Cy3, BG-sulfo-Cy5, BG-Cy5 were determined by fluorescence polarization analogous to HT7 affinities towards CA substrates described above with the following changes: fluorescent substrates were titrated at a final concentration of 50 nM against protein concentrations ranging from (0 – 250 μM) at room temperature using 0.1 g/L BSA and 1 mM DTT (SNAP-FP buffer). The FP value of each dye fully reacted with the native SNAP/SNAPf was used to improve fitting of the upper plateau of the curves by adding an extra data point at protein concentration of 0.005 M.

### Affinity of HT7 towards methyl-amide fluorophores

Binding affinities of HT7 towards methylamide fluorophores were determined by fluorescence polarization analogous to CA substrates described above with following changes: fluorescent substrates were used at a final concentration of 50 nM and measurements were performed at room temperature.

### Affinity ofHT7^D106A^ towards CA-Ac via FP competition assay

Binding affinity of HT7^D106A^ towards CA-Ac was determined by a fluorescence polarization competition assay against CA-TMR. 5 μM protein and 50 nM CA-TMR were titrated against CA-Ac concentrations ranging from 80 μM to 10 mM in activity buffer supplemented with 0.5 g/L BSA. Assays were performed in low-volume non-binding black 384-well plates (Corning Inc.) with a final volume of 20 μL using a microplate reader (Spark20M^®^, Tecan). All measurements were performed in triplicates at 37°C, filter settings are listed in **Table 3**. Obtained FP values were averaged and fitted to a 4 parameter logistic curve (equation **3**) to estimate the I50 value. The lower plateau was fixed to the measured FP value of the free dye to improve the fit. The dissociation constant of CA-Ac was calculated as described by Rossi and Taylor (2011) (48).

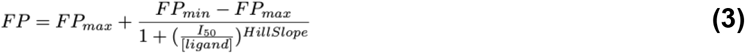

with FP_min_: fluorescence polarization of the free fluorophore (lower plateau), FP_max_: maximal fluorescence polarization of fully bound fluorophore (upper plateau), I_50_: half maximal effective concentration, HillSlope: hill slope and [ligand]: ligand concentration.

### Affinity of SNAPC^145A^ towards non-fluorescent substrates via FP competition assay

Binding affinities of SNAP^C145A^ towards BG, CP, BG-Ac, CP-Ac and BC-Ac to were obtained as previously described for HT7 by titrating 5 μM protein and 50 nM CP-TMR against non-fluorescent substrate concentrations ranging from 150 nM to 1.5 mM. Experimental conditions and data analysis were identical despite that 1 mM DTT was added to the buffer and the assay was performed at room temperature.

### Calculation of free binding energy from K_d_

Free binding energies were calculated from K_d_ values according to equation 4:

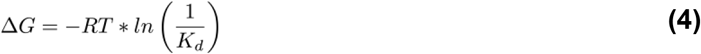

with Δ*G*: free binding energy, R: universal gas constant, T: temperature and K_d_: dissociation constant.

### HT7 and HOB labeling kinetics via stopped-flow

Labeling kinetics of HT7 with CA-TMR, CA-JF549, CA-CPY, CA-LIVE580 and CA-JF669 and labeling kinetics of HOB with CA-TMR were measured by recording fluorescence anisotropy changes over time using a BioLogic SFM-400 stopped-flow instrument (BioLogic Science Instruments, Claix, France) in single mixing configuration at 37°C. Monochromator wavelengths for excitation and long pass filters used for detection are listed in **Table 4**. HT7 protein and substrates in activity buffer were mixed in a 1:1 stoichiometry in order to reach recordable speed of these fast reactions and increase information content of the traces. Concentrations were varied from 0.125 μM to 1 μM. The anisotropy of the free substrate was measured to obtain a baseline.

**Table 4:** Monochromator excitation wavelengths and filters used for stopped-flow measurements

| Fluorophore | Excitation wavelength [nm] | Emission filter [nm] |
| --- | --- | --- |
| TMR / JF549 | 555 | 570 Long Path |
| CPY | 610 | 630 Long Path |
| LIVE580 | 603 | 630 Long Path |
| JF669 | 669 | 690 Long Path |

The dead time of the instrument was measured according to the manufacturer protocol (BioLogic Technical note #53) by recording the fluorescence decay during the pseudo-first order reaction of *N*-acetyl-L-tryptophanamide with a large excess of *N*-bromosuccinimide and fitting the data to the first order reaction rate law.

### SNAP labeling kinetics via stopped-flow

Labeling kinetics of SNAP with BG-TMR were measured via stopped-flow analogous to HT7 kinetics described above but final substrate concentration was fixed at 2 μM and the protein concentration was varied from 1.875 μM to 50 μM. The activity buffer was supplemented with 1 mM DTT.

### HT7 and HOB labeling kinetics via microplate reader

Labeling kinetics of HT7 and HOB with CA-Alexa488 were measured by recording FP over time using a microplate reader (Spark20M^®^, Tecan). The final concentration of fluorophore substrate remained constant (50 nM) with varying protein concentrations (200 nM – 256 μM) in activity buffer supplemented with 0.5 g/L of BSA. Labeling reactions were started by adding the fluorophore substrate using either multichannel pipets or the injector module of the plate reader. Assays were performed in black non-binding flat bottom 96-well plates (Corning Inc.) with a final reaction volume of 200 μL. All measurements were performed in triplicates at 37°C with filter settings listed in **Table 3**. The FP of the free substrate was measured to obtain a baseline.

### HT7-tag competitive labeling kinetics

Competitive kinetics were measured by recording FP over time using a microplate reader (Spark20M^®^, Tecan). The final concentration of CA-Alexa488 (50 nM) and HT7 protein (200 nM) remained constant with varying concentrations of non-fluorescent substrates (0 – 1 μM) in activity buffer supplemented with 0.5 g/L of BSA. Assays were performed in black non-binding flat bottom 96-well plates with a final reaction volume of 200 μL. Labeling reactions were started by adding the HT7 protein to wells containing CA-Alexa488 and non-fluorescent substrates using an electronic 96 channel pipettor (Integra Bioscience Corp., Hudson, NH, USA). All measurements were performed in triplicates at 37°C with filter settings listed in **Table 3**. The FP of free CA-Alexa488 was measured to obtain a baseline.

### SNAP and CLIP labeling kinetics via microplate reader

Labeling kinetics of SNAP and CLIP substrates were measured by recording FP over time using a microplate reader analogously to HT7 labeling kinetics described above with the following changes: fluorescent substrate concentration was fixed to 20 nM and protein concentrations were varied from 15 nM to 900 nM. Measurements were performed in SNAP-FP buffer. Kinetics with substrates that showed adsorption to plastic were recorded in a black quartz 96-well plate (Hellma GmbH, Müllheim, Germany).

### SNAP competitive labeling kinetics

Competitive kinetics were measured by recording FP over time using a microplate reader analogous to HT7 competition kinetics described above using 100 nM of BG-Alexa488 as fluorescent substrate in SNAP-FP buffer.

### Analysis of stopped-flow data

Kinetic stopped-flow data was pre-processed using a custom R script (49, 50). Recorded pre-trigger time points were removed and time points were adjusted to start at t = 0. Values from replicates were averaged. The anisotropy of the free dye was calculated by averaging anisotropy values of the baseline measurements. Pre-processed data was fit to a kinetic model (**5, 6**) described by the differential equations **7**–**10** using the DynaFit software (51). The anisotropy of the free dye and the mixing delay of the stopped-flow machine were set as fixed offset and delay parameters in DynaFit. It was assumed that the protein substrate complex and the reacted product are contributing equally to the anisotropy signal. Hence, the response for both species was set equal in DynaFit and fitted together with the kinetic constants. Standard deviations (normal distribution verified) and confidence intervals of fitted parameters were estimated with the Monte Carlo method (52) with standard settings (*N* = 1000, 5% worst fits discarded). In case of SNAP kinetics with BG-TMR, the substrate concentration was fitted by DynaFit in order to rule out quantification errors of the BG quenched fluorophore. Accurate fitting of the concentration was ensured by including conditions in which protein is limiting and no maximum FP value was reached. Data points and predictions based on the fitted models were plotted using R. Fluorescence intensity changes upon protein binding were verified to be minimal (< 12 %) and hence not noticeably biasing the fluorescence anisotropy.

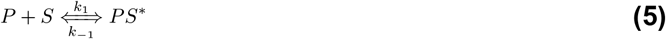

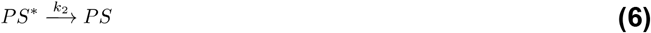

with P: SLP protein, S: SLP substrate, PS*: protein substrate complex and PS: protein substrate conjugate.

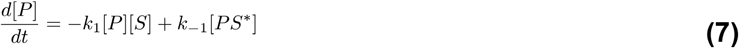

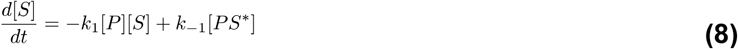

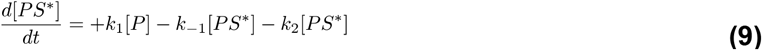

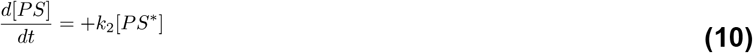

The derived parameters K_d_ (dissociation constant) and k_app_ (apparent first order reaction rate) were calculated using the following equations:

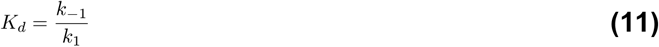

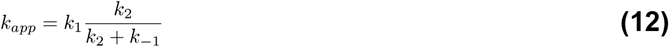

### Analysis of kinetic microplate reader data

Kinetic data from microplate reader assays was fitted to a simplified kinetic model (**13**) described by the differential equations **14**–**16** using DynaFit. Dead time of the measurements and baseline FP value were put in as fixed parameters. Standard deviations (normal distribution verified) and confidence intervals of fitted parameters were estimated with the Monte Carlo method with standard settings (*N* = 1000, 5% worst fits discarded). In case of BG, CP and BC kinetics, the substrate concentration was fitted by DynaFit in order to rule out quantification errors of the BG, CP or BC fluorophores. Accurate fitting of the concentration was ensured by including conditions in which protein is limiting and no maximum FP value was reached. Data points and predictions based on the fitted models were plotted using R.

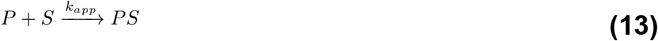

with P: SLP protein, S: SLP substrate and PS: protein substrate conjugate.

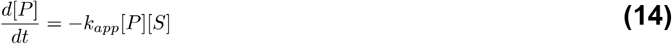

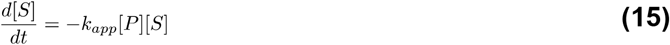

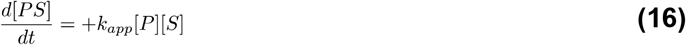

In some cases, a slow second phase (k_3_) was observed in the kinetic data that could not be described by the simplified model **13**. This data was fit to an expanded model that includes a potential conformational change in a second step (**17, 18**).

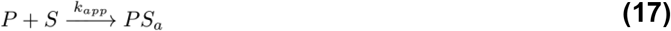

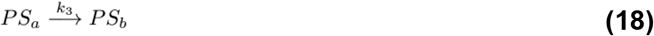

with P: SLP protein, S: SLP substrate, PS_a_: protein substrate conjugate state A and PS_b_: protein substrate conjugate state B.

### Analysis of competition kinetics

Data was fitted to a simplified kinetic competition model (**19, 20**) described by the differential equations **21**–**25** using DynaFit. Dead time of the measurements and baseline FP value were put in as fixed parameters. Standard deviations (normal distribution verified) and confidence intervals of fitted parameters were estimated with the Monte Carlo method with standard settings (*N* = 1000, 5% worst fits discarded).

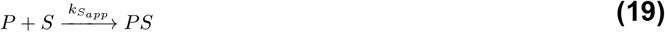

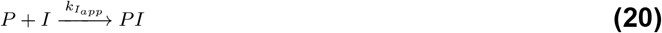

with P: SLP protein, S: fluorescent SLP substrate, I: non-fluorescent SLP substrate (inhibitor), PS: protein fluorescent substrate conjugate and PI: protein non-fluorescent substrate conjugate.

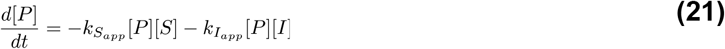

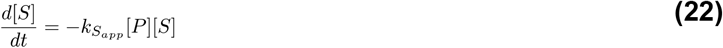

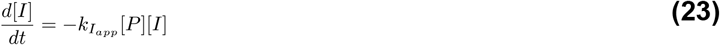

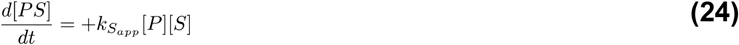

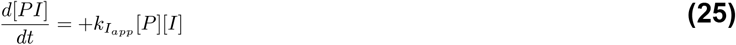

### Protein crystallization

For crystallization trials, protein purification tags were removed by overnight cleavage with TEV protease at 30°C as previously described (53). Cleaved proteins were purified by affinity-tag purification using a HisTrap FF crude column (Cytiva) on an ÄktäPure FPLC, collecting the flow-through. Proteins were further separated by size exclusion chromatography (HiLoad 26/600 Superdex 75, Cytiva) and concentrated using Ultra-4 or 15 mL centrifugal filter devices (Amicon, Merck). Correct size and high purity were verified via SDS-PAGE and LC-MS analysis. Protein labeling was performed in activity buffer, overnight at RT using fluorophore substrates at 10 μM (CA-TMR/CA-CPY and BG-TMR for HT7/HOB and SNAP, respectively) in presence of 5 μM (3 mg) of protein. After concentration to about 200 μL, excess of fluorophore substrate was removed by buffer exchange using Illustra microspin G-25 columns (Cytiva) according to the manufacturer instructions. Protein labeling was verified by SDS-PAGE fluorescence scan and LC-MS analysis. Protein concentrations were adjusted between 10 and 20 mg/mL and submitted to crystallization trials using different commercial screens mixing in 200 nL final volume protein solution:crystallization solution (1:1) using a Mosquito robot (TTP Labtech).

### HT7 crystal structures

Crystallization was performed at 20°C using the vapor-diffusion method. Crystals of HT7 labeled with a chloroalkane-PEG-tetramethylrhodamine (CA-TMR) fluorophore substrate were grown by mixing equal volumes of protein solution at 20 mg/ml in 50 mM HEPES pH 7.3, 50 mM sodium chloride and a reservoir solution containing 0.1 M MES pH 6.0, 1.0 M lithium chloride and 15% (m/v) PEG 6000. The crystals were briefly washed in cryoprotectant solution consisting of the reservoir solution with glycerol added to a final concentration of 20% (v/v), prior to flash-cooling in liquid nitrogen. Crystals of HT7 labeled with a chloroalkane-PEG-carbopyronine (CA-CPY) fluorophore substrate were obtained by mixing equal volumes of protein solution at 15 mg/ml in 50 mM HEPES pH 7.3, 50 mM sodium chloride and precipitant solution containing 0.1 M Bicine pH 9.0 and 1.7 M ammonium sulfate. The crystals were briefly washed in cryoprotectant solution consisting of the reservoir solution supplemented with 20% (v/v) ethylene glycol before flash-cooling in liquid nitrogen. Crystals of HT7-based Oligonucleotide Binder (HOB) labeled with a CA-TMR fluorophore substrate were grown by mixing equal volumes of protein solution at 9.0 mg/ml in 50 mM HEPES pH 7.3, 50 mM sodium chloride and a reservoir solution composed of 0.2 M calcium acetate and 20% (m/v) PEG 3350. Prior to flash-cooling in liquid nitrogen, the crystals were stepwise transferred into a reservoir solution with PEG 3350 concentration increased to 30 and 40% (m/v).

Single crystal X-ray diffraction data was collected at 100 K on the X10SA beamline at the SLS (PSI, Villigen, Switzerland). All data were processed with XDS (54). The structures of HT7 labeled with TMR was determined by molecular replacement (MR) using Phaser (55) and PDB ID 5UY1 coordinates as a search model. The structure of HT7 labeled with CPY and HOB labeled with TMR were subsequently determined by molecular replacement using HT7-TMR as a search model. Geometrical restraints for TMR and CPY were generated using Grade server (56). The final models were optimized in iterative cycles of manual rebuilding using Coot (57) and refinement using Refmac5 (58) and phenix.refine (59). Data collection and refinement statistics are summarized in **Table S4**, model quality was validated with MolProbity (60) as implemented in PHENIX.

### SNAP crystal structure

SNAP-TMR crystals were obtained on the crystallography platform of EPFL using the SNAP^cx^-tag construct that features the sequence of SNAP identical to available SNAP crystal structures (PDB ID 3L00, 3KZZ and 3KZY). Previously crystallized SNAP features the mutation P179R involved in the crystal packing suggesting its important role for crystallization (28). Crystals were obtained in different conditions including in 100 mM Sodium HEPES pH 7.5, 25% PEG 8000 from the PEG suite screen (Qiagen) after 48 hours at 18°C. Single crystals were fished and placed in a cryoprotectant solution (containing the crystallization solution supplemented with 20% (v/v) glycerol) before being flash frozen in liquid nitrogen. Single crystal X-ray diffraction data was collected on the ID29 beamline at the ESRF (Grenoble, France). Integration, scaling, molecular replacement (using PDB ID 3L00 as starting model) and refinement were performed as explained for HT7. Refinement statistics can be found in **Table S4**.

### *SNAPf* in silico *modeling*

The glutamic acid in position 30 of the SNAP-TMR structure (PDB ID 6Y8P) was modeled as an arginine using the mutate function using the software SYBYL-X1.3 (Tripos Int., USA). A side-chain conformation for the arginine was selected from the rotamer source library of Lovell and minimized with few steps with no steric clashes and no direct contact with another positive charges as criteria.

## Data availability and analysis

Atomic coordinates and structure factors were deposited in the Protein Data Bank (PDB) under accession codes 6Y7A (HT7-TMR), 6ZCC (HOB-TMR), 6Y7B (HT7-CPY) and 6Y8P (SNAP-TMR). Analysis was conducted on PyMOL (61). OMIT maps were generated using Phenix (62). Root mean square deviations (RMSDs) were obtained using the cealign command from PyMOL. Electrostatic potentials were generated using the adaptive poisson–boltzmann solver (APBS) (63) as PyMOL plugin including the PDB2PQR software (64). Plasmids from this study are available at Addgene (167266-167275).

## Acknowledgements

The authors thank Ilme Schlichting for X-ray data collection. HaloTag diffraction data were collected at the Swiss Light Source, beamline X10SA, of the Paul Scherrer Institute, Villigen, Switzerland. The authors thank Florence Pojer for supporting the SNAP-TMR crystallization on the EPFL platform. The ESRF is acknowledged for access to beamlines and facilities for molecular biology via its in-house research programme. The authors thank Andrea Bergner (MPIMF) and Bettina Mathes (MPIMF) for providing proteins and fluorophore substrates, respectively. The authors thank Dr. L. Lavis (HHMI, Ashburn, VA, USA) and Dr. A.D.N. Butkevich (MPI-MF, Heidelberg, Germany) for providing HaloTag substrates. We thank all members of the Johnsson lab for critical discussions. This work was supported by the Ecole Polytechnique Federale de Lausanne (EPFL), the Max Planck Society and the Deutsche Forschungsgemeinschaft (DFG, German Research Foundation), SFB 1129.

## Author contributions

J.W., S.K., J.R., J.H. and K.J. designed the experiments.

J.W., S.K., J.H. and J.T. performed the biochemistry experiments.

J.H., T.T., G.G. crystalized and solved the SNAP-TMR structure.

M.T. and J.H. crystalized and solved the HaloTag structures.

U.U. performed the structural modeling work.

S.K., J.W., J.K., N.M. and L.X. synthesized the compounds used in the study.

K.J., J.H., J.W. and S.K. wrote the manuscript with input from all authors.

## Competing financial information

K.J. is inventor on patents filed by MPG and EPFL on fluorophores and labeling technologies.

## Additional information

Further information and requests for resources and reagents should be directed to and will be fulfilled by Julien Hiblot and Kai Johnsson.

## Supplementary information

### Chemical Synthesis

#### General information

All chemical reagents and (anhydrous) solvents for synthesis were purchased from commercial suppliers (Merck KGaA, Darmstadt, Germany; Honeywell, Charlotte, NC, USA; TCI, Tokyo, Japan; Thermo Fisher Scientific, Waltham, MA, USA; SiChem, Bremen, Germany) and were used without further purification or distillation. Anhydrous solvents were handled under argon atmosphere. SLP substrates were purchased from commercial sources, synthesized according to published procedures or gifts from colleagues. Details are given in **Material Table**.

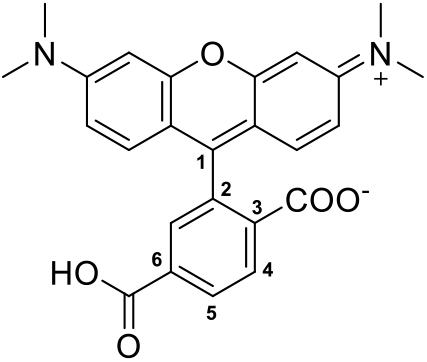

^1^H- and ^13^C-NMR spectra were recorded in deuterated solvents on a Bruker (Bruker Corp., Billerica, MA, USA) DPX400 (400 MHz for ^1^H, 101 MHz for ^13^C, respectively) or on a Bruker AVANCE III HD 400 (400 MHz for ^1^H, 101 MHz for ^13^C, respectively) equipped with a CryoProbe. Chemical shifts (δ) are reported in ppm referenced to the residual solvent peaks of DMSO-*d_6_* (*δ*_H_ = 2.50 ppm, *δ*_C_ = 39.52 ppm), acetone-*d_6_* (*δ*_H_ = 2.05 ppm, *δ*_C_(CH_3_) = 29.84 ppm, *δ*_C_(CO) = 206.26 ppm) or CDCl_3_ (*δ*_H_ = 7.26 ppm, *δ*_C_ = 77.16 ppm). Coupling constants *J* are reported in Hz and corresponding multiplicities are abbreviates as follows: s = singlet, d = doublet, t = triplet, q = quartet, p = pentet, m = multiplet and br = broad.

Reaction progress was monitored by thin layer chromatography (TLC) (Silica gel 60G F_254_ on TLC glass plates) in appropriate solvents. Reaction spots were visualized under UV lamp (254 nm or 366 nm) and/or by staining solutions. LC-MS was performed on a Shimadzu MS2020 (Shimadzu Corp., Kyoto, Japan) connected to a Nexera UHPLC system equipped with a Waters (Waters Crop., Milford, MA, USA) ACQUITY UPLC BEH C18 (1.7 μm, 2.1×50 mm) column. Buffer A: 0.1% formic acid in H_2_O, Buffer B: acetonitrile. Measurements were done with an analytical gradient from 10% to 90% B over 6 min or from 1% to 90% B over 10 min.

Normal phase flash chromatography was performed on self-packed silica gel (60 M, 0.04 - 0.063 mm, Macherey-Nagel GmbH & Co. KG, Düren, Germany) columns or by using an Isolera One system (Biotage Sweden AB, Uppsala, Sweden) using pre-packed silica gel columns (ultra pure silica gel 12 g or 25 g). Solvent compositions are reported individually in parentheses.

Preparative reversed phase high-performance liquid chromatography (RP-HPLC) was conducted using a Waters SunFire™ Prep C18 OBDTM column (10 × 150 mm, 5 μm pore size, 4 mL/min. flow rate) or an Ascentis (Merck KGaA, Darmstadt, Germany) C18 column (10 × 250 mm, 5 μm pore size, 8 mL/min. flow rate) on either a Waters Alliance e2695 separation module connected to a 2998 PDA detector or a Dionex system equipped with an UVD (170 U, UV-Vis detector). Solvent A: 0.1%TFA in H_2_O, Solvent B: acetonitrile.

High resolution mass spectra (HRMS) were measured by the MS-service of the EPF Lausanne (SSMI) on a Waters Xevo^®^ G2-S Q-Tof spectrometer (Waters, Milford, MA, USA) with electron spray ionization (ESI) or by the MS-facility of the Max Planck Institute for Medical Research on a Bruker maXis IITM ETD.

**Material Table:**
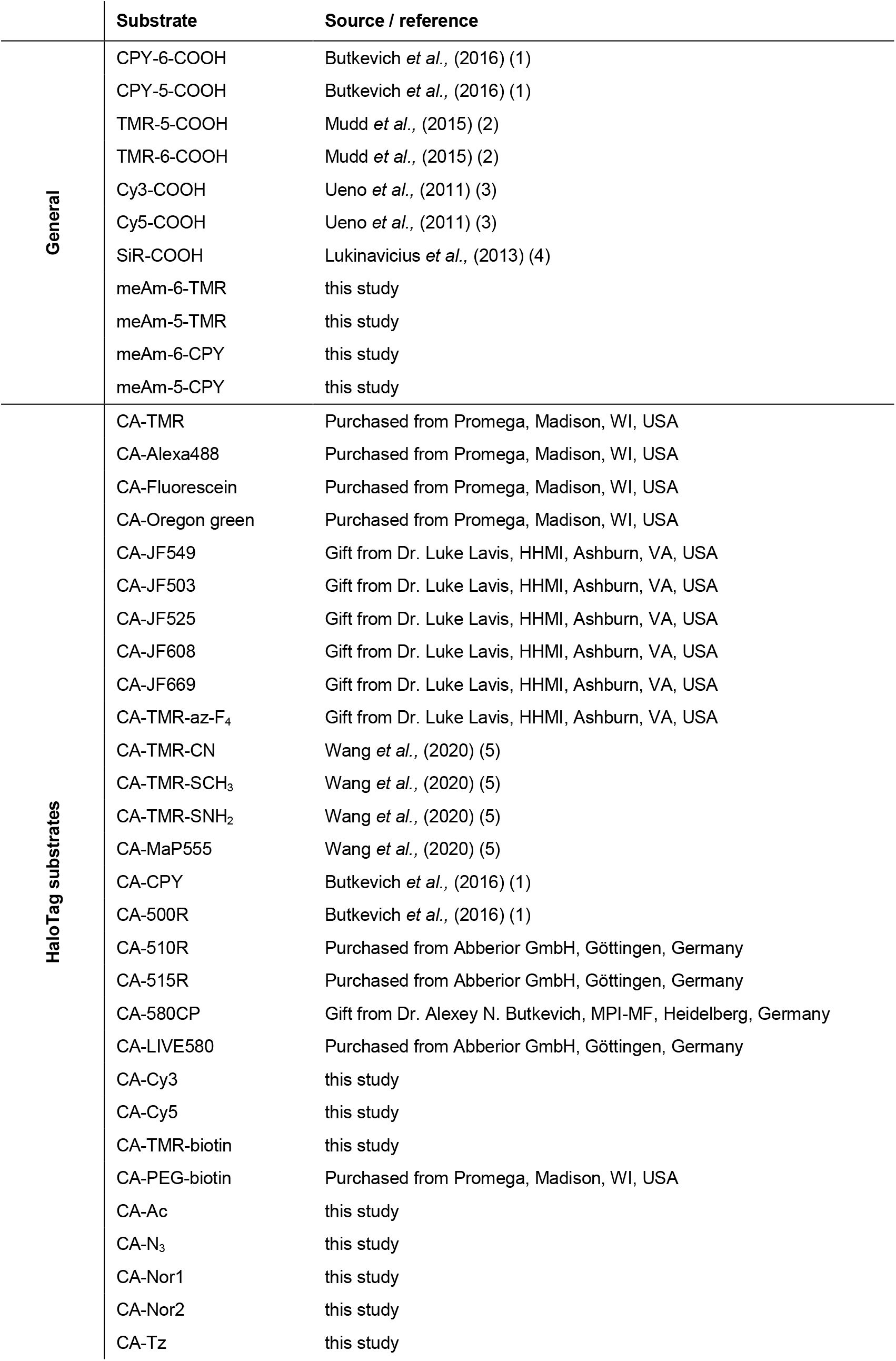
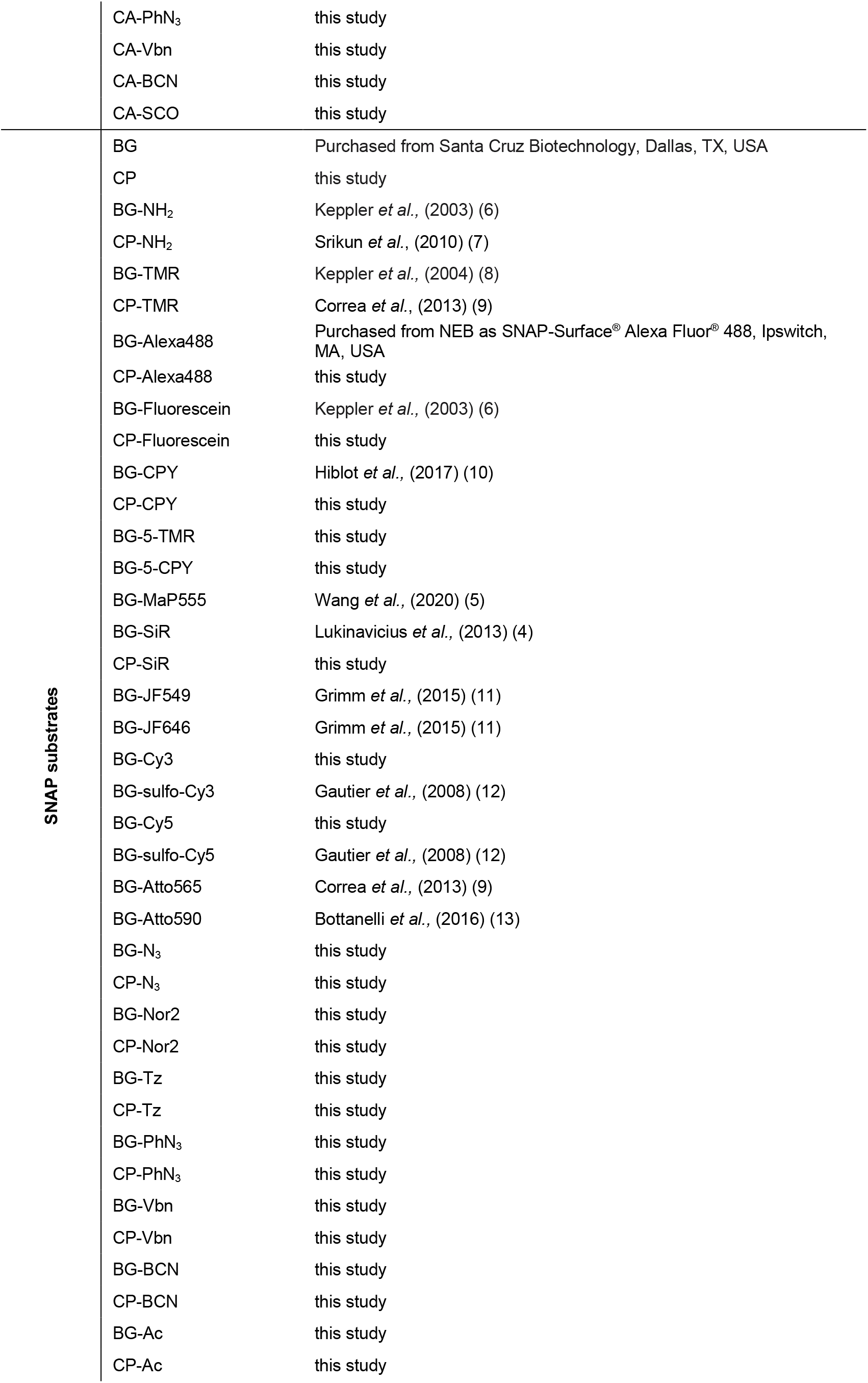
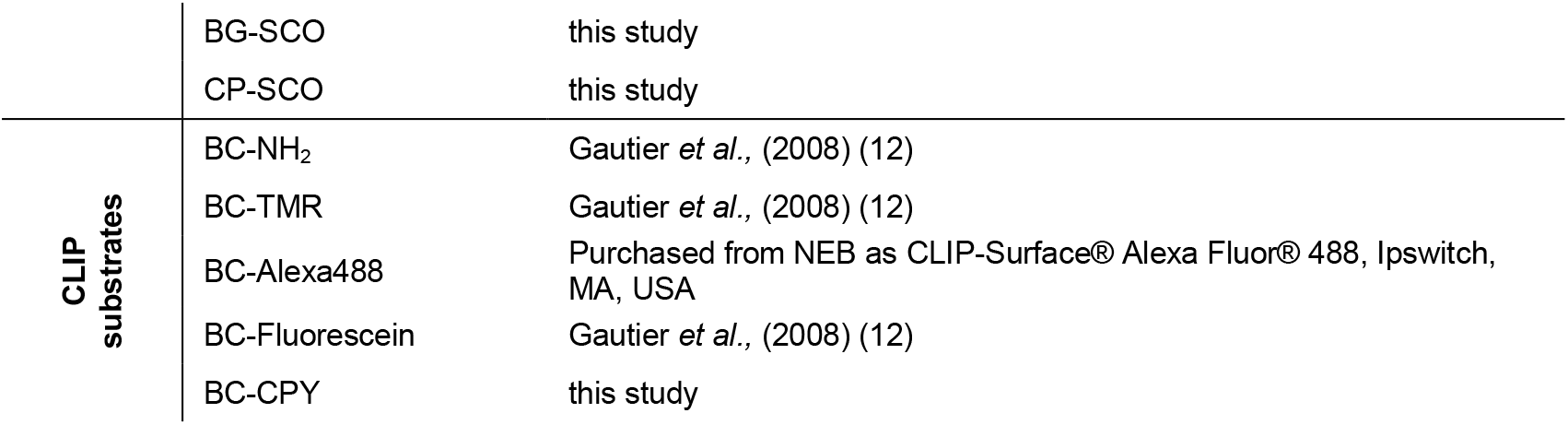
Substrate and chemical source used in the study

### Chemical Synthesis

#### 1.1 Synthesis of substrate amines

##### 1.1.1 2-(2-((6-chlorohexyl)oxy)ethoxy)ethan-1-amine (CA-NH_2_)

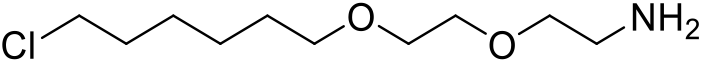

CA-NH_2_ was synthesized according to the procedure from Zhang *et al*. 2006 (14).

##### 1.1.2 6-((4-(aminomethyl)benzyl)oxy)-9*H*-purin-2-amine (BG-NH_2_)

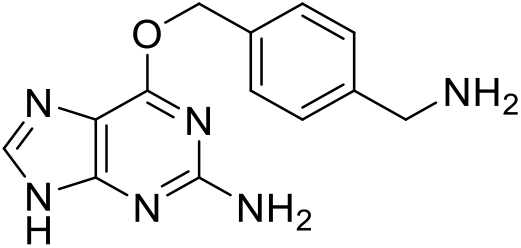

BG-NH_2_ was synthesized according to the procedure from Keppler *et al*. 2003 (6).

##### 1.1.3 4-((4-(aminomethyl)benzyl)oxy)-6-chloropyrimidin-2-amine (CP-NH_2_)

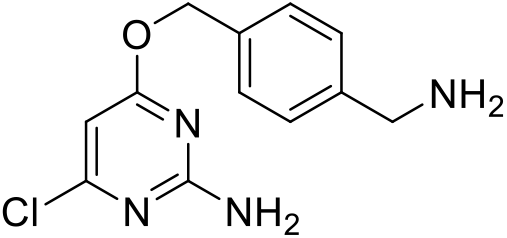

CP-NH_2_ was synthesized according to the procedure from Srikun *et al*. 2010 (7).

##### 1.1.4 2-((4-(aminomethyl)benzyl)oxy)pyrimidin-4-amine (BC-NH_2_)

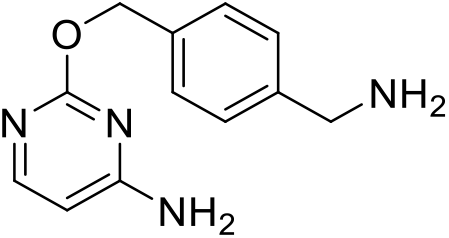

BC-NH_2_ was synthesized according to the procedure from Gautier *et al*. 2008 (12).

#### 1.2 General procedure A for peptide coupling reactions

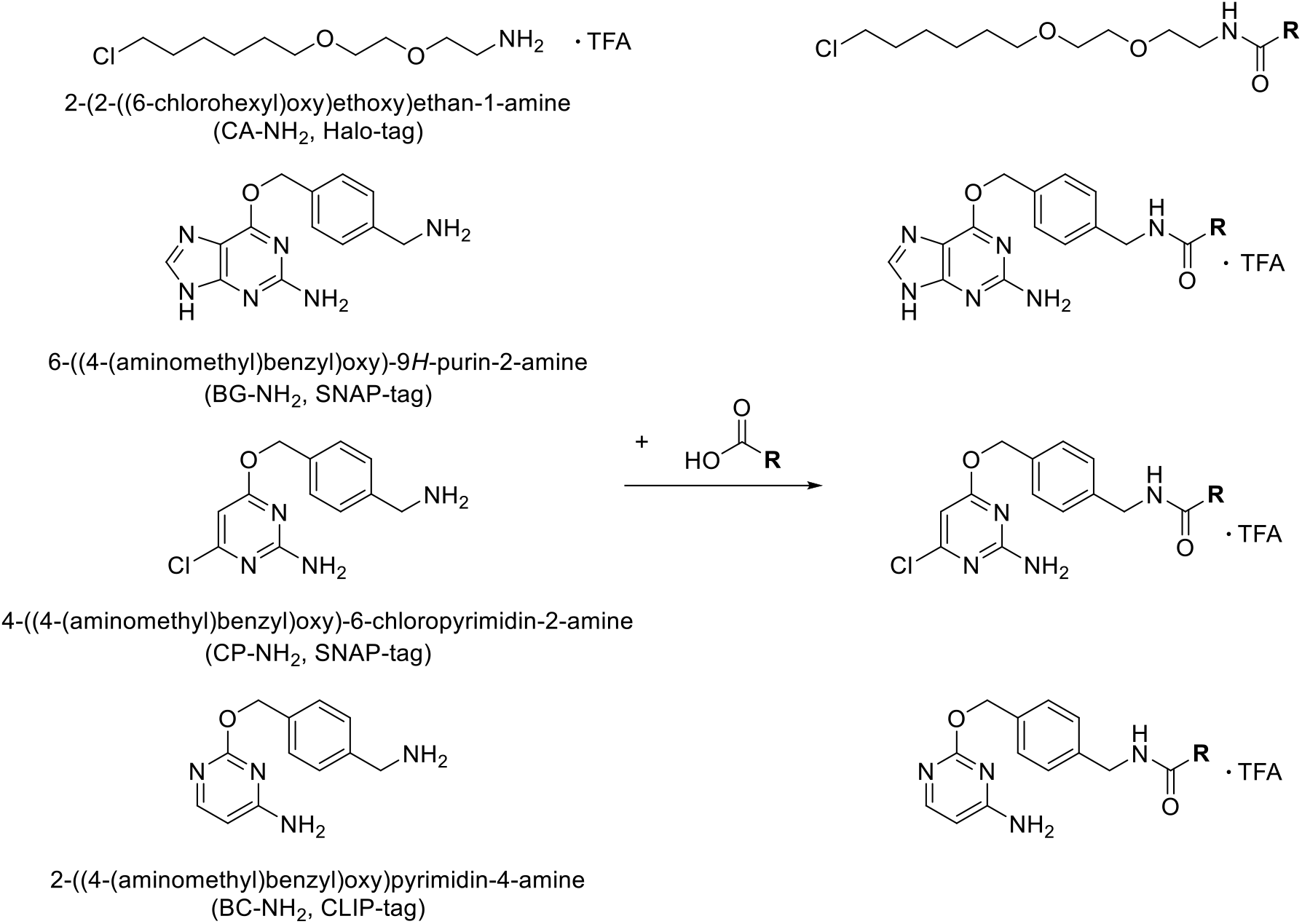

To a solution of TSTU (1.2 equiv.) in dry DMSO (0.3 mL), DIPEA (10.2 equiv. for Halo-tag-, 5.0 equiv. for SNAP-substrates) and different carboxylic acids (1.1 equiv.) were added. After 5 min., a solution of 10 mg of corresponding amine (1.0 equiv.) in dry DMSO (0.1 mL) was added and the reaction mixture was stirred at r.t. for 2 hours. The reaction mixture was quenched by addition of water (100 μL) and acidified with acetic acid (50 μL), then purified by semi-preparative HPLC, eluted with a gradient of MeCN/H_2_O + 0.1% TFA (equilibration at 15% MeCN for 5 min, then gradient of 15 - 100% MeCN over 25 min, followed by 100% MeCN for 10 min.). Fractions containing the desired product were combined and lyophilized. Final compounds were stored as DMSO stocks for biochemical testing.

#### 1.3 HT7 substrates

##### 1.3.1 2-azido-N-(2-(2-((6-chlorohexyl)oxy)ethoxy)ethyl)acetamide (CA-N_3_)

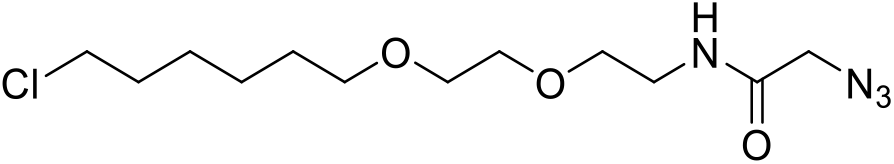

Reaction was conducted according to general procedure A using CA-NH_2_ and 2-azidoacetic acid (4.6 μL, 32.6 μmol). The desired product (4.6 mg, 15.0 μmol) was obtained as a yellowish oil in 51% yield.

**^1^H NMR** (400 MHz, DMSO-*d*_6_) *δ* [ppm] = 8.15 (t, *J* = 5.8 Hz, 1H), 3.81 (s, 2H), 3.62 (t, *J* = 6.6 Hz, 2H), 3.53 – 3.40 (m, 6H), 3.37 (t, *J* = 6.6 Hz, 2H), 3.24 (dd, *J* = 5.7 Hz, *J* = 5.8 Hz, 2H), 1.75 – 1.65 (m, 2H), 1.54 – 1.43 (m, 2H), 1.43 – 1.25 (m, 4H).

**^13^C NMR** (101 MHz, DMSO-*d*_6_) *δ* [ppm] = 167.31, 70.17, 69.56, 69.40, 68.83, 50.69, 45.36, 38.67, 32.00, 29.04, 26.10, 24.91.

**HRMS** (ESI): calc. for C_12_H_23_ClN_4_NaO_3_^+^ [M+Na]^+^: 329.1351; found 329.1354.

##### 1.3.2 (1R,4R)-N-(2-(2-((6-chlorohexyl)oxy)ethoxy)ethyl)bicyclo[2.2.1]hept-5-ene-2-carboxamide (CA-Nor1)

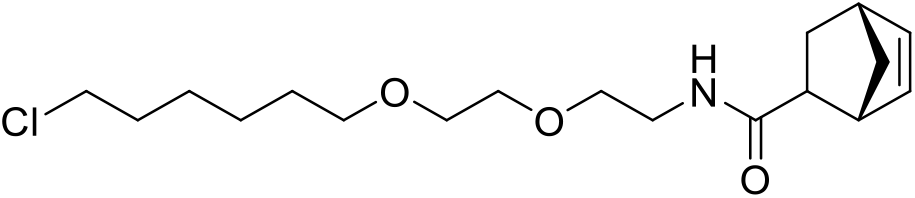

Reaction was conducted according to general procedure A using CA-NH_2_ and (1R,4R)-bicyclo[2.2.1]hept-5-ene-2-carboxylic acid (6.2 μL, 32.6 μmol). The desired endo-isomer (5.6 mg, 16.3 μmol) of was obtained in 55% yield.

**^1^H NMR** (400 MHz, DMSO-*d*_6_) *δ* [ppm] = 7.59 (t, *J* = 5.7 Hz, 1H), 6.08 (dd, *J* = 5.8, 3.1 Hz, 1H), 5.80 (dd, *J* = 5.8, 3.0 Hz, 1H), 3.62 (t, *J* = 6.6 Hz, 2H), 3.54 – 3.30 (m, 8H), 3.23 – 3.02 (m, 3H), 2.84 – 2.71 (m, 2H), 1.77 – 1.63 (m, 3H), 1.55 – 1.43 (m, 2H), 1.42 – 1.18 (m, 7H).

**^13^C NMR** (101 MHz, DMSO-*d*_6_) *δ*[ppm] = 172.86, 136.76, 132.18, 70.18, 69.58, 69.45, 69.09, 49.35, 45.59, 45.37, 43.25, 42.08, 38.55, 32.02, 29.09, 28.35, 26.13, 24.94.

**HRMS (ESI)** calc. for C_18_H^31^ClNO_3_^+^ [M+H]^+^: 344.1987; found 344.1989.

##### 1.3.3 2-((1S,4S)-bicyclo[2.2.1]hept-5-en-2-yl)-N-(2-(2-((6-chlorohexyl)oxy)ethoxy)ethyl)acetamide (CA-Nor2)

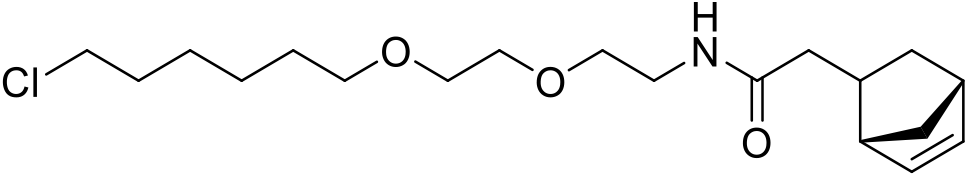

Reaction was conducted according to general procedure A using CA-NH_2_ and 2-((1S,4S)-bicyclo[2.2.1]hept-5-en-2-yl)acetic acid (5.6 μL, 32.6 μmol) yielding 6.4 mg (17.9 μmol) of the desired product as a colorless oil in 60% yield.

**^1^H NMR** (400 MHz, DMSO-*d*_6_) *δ* [ppm] = 7.73 (t, *J* = 5.7 Hz, 1H), 6.15 (dd, *J* = 5.8, 3.0 Hz, 1H), 5.95 (dd, *J* = 5.8, 2.9 Hz, 1H), 3.62 (t, *J* = 6.6 Hz, 2H), 3.47 – 3.36 (m, 8H), 3.23 – 3.09 (m, 2H), 2.76 – 2.68 (m, 2H), 2.40 – 2.29 (m, 1H), 1.89 – 1.74 (m, 3H), 1.74 – 1.65 (m, 2H), 1.53 – 1.43 (m, 2H), 1.39 – 1.17 (m, 6H), 0.47 (m, *J* = 11.5, 4.4, 2.6 Hz, 1H).

**^13^C NMR** (101 MHz, DMSO-*d*_6_) *δ*[ppm] = 171.61, 136.88, 132.47, 70.15, 69.52, 69.42, 69.08, 49.03, 45.32, 45.13, 42.02, 40.58, 40.14, 39.93, 39.73, 39.51, 39.31, 39.10, 38.89, 38.35, 35.06, 31.99, 31.37, 29.04, 26.08, 24.89.

**HRMS** (ESI) calc. for C_19_H_32_ClNNaO_3_^+^ [M+Na]^+^: 380.1963; found 380.1963.

##### 1.3.4 N-(2-(2-((6-chlorohexyl)oxy)ethoxy)ethyl)-2-(4-(6-methyl-1,2,4,5-tetrazin-3-yl)phenyl)acetamide (CA-Tz)

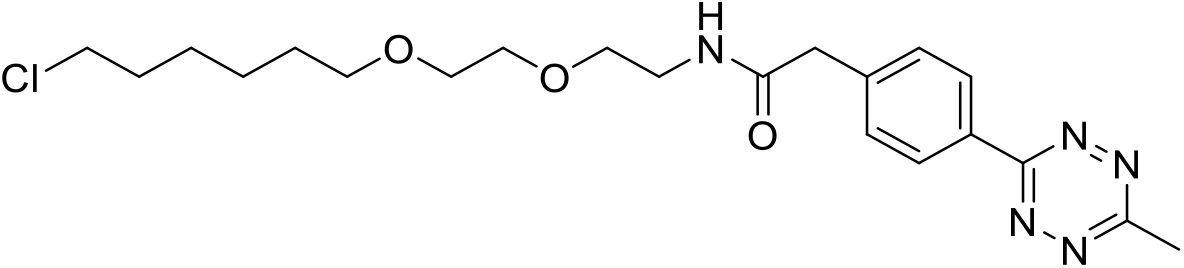

Reaction was conducted according to general procedure A using CA-NH_2_ and 2-(4-(6-methyl-1,2,4,5-tetrazin-3-yl)phenyl)acetic acid (7.5 mg, 32.6 μmol) yielding 7.4 mg (17.0 μmol) of the desired product as a rose solid in 57% yield.

**^1^H NMR** (400 MHz, DMSO-*d*_6_) *δ* [ppm] = 8.44 – 8.36 (m, 2H), 8.23 (t, *J* = 5.6 Hz, 1H), 7.58 – 7.50 (m, 2H), 3.61 (t, *J* = 6.6 Hz, 2H), 3.56 (s, 2H), 3.53 – 3.40 (m, 6H), 3.36 (t, *J* = 6.6 Hz, 2H), 3.23 (q, *J* = 5.7 Hz, 2H), 2.99 (s, 3H), 1.75 – 1.62 (m, 2H), 1.53 – 1.42 (m, 2H), 1.42 – 1.25 (m, 4H).

**^13^C NMR** (101 MHz, DMSO-*d*_6_) *δ* [ppm] = 169.58, 167.05, 163.22, 141.29, 130.05, 130.00, 127.28, 70.20, 69.60, 69.45, 69.05, 45.37, 42.19, 38.79, 32.02, 29.07, 26.12, 24.93, 20.83.

**HRMS** (ESI) calc. for C_21_H_31_ClN_5_O_3_^+^ [M+H]^+^: 436.2110; found 436.2113.

##### 1.3.5 4-azido-N-(2-(2-((6-chlorohexyl)oxy)ethoxy)ethyl)benzamide (CA-PhN3)

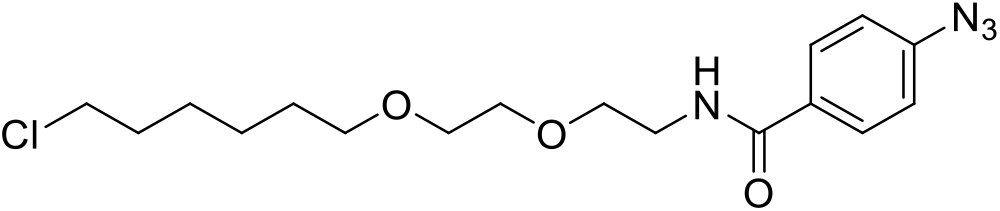

Reaction was conducted according to general procedure A using CA-NH_2_ and 4-azidobenzoic acid (5.3 mg, 32.6 μmol) to obtain 6.1 mg (15.5 μmol) of the desired product as a colorless oil in 56% yield.

**^1^H NMR** (400 MHz, DMSO-*d*_6_) *δ* [ppm] = 8.52 (t, *J* = 5.6 Hz, 1H), 7.90 (d, *J* = 8.6 Hz, 2H), 7.20 (d, *J* = 8.6 Hz, 2H), 3.60 (t, *J* = 6.6 Hz, 2H), 3.56 – 3.49 (m, 4H), 3.50 – 3.44 (m, 2H), 3.44 – 3.30 (m, 4H), 1.74 – 1.61 (m, 2H), 1.51 – 1.39 (m, 2H), 1.40 – 1.20 (m, 4H).

**^13^C NMR** (101 MHz, DMSO-*d*_6_) *δ* [ppm] = 165.23, 142.19, 130.95, 129.06, 118.85, 70.17, 69.62, 69.40, 68.84, 45.35, 39.21, 32.00, 29.07, 26.12, 24.91.

**HRMS** (ESI) calc. for C_17_H_25_ClN_4_NaO_3_^+^ [M+Na]^+^: 391.1507; found 391.1511.

###### 1.3.5.1 N-(2-(2-((6-chlorohexyl)oxy)ethoxy)ethyl)-4-vinylbenzamide (CA-Vbn)

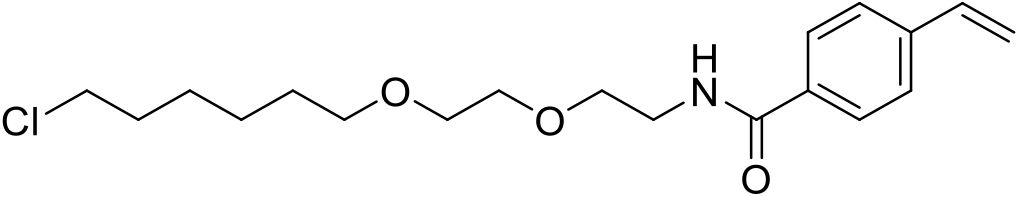

Reaction was conducted according to general procedure A using CA-NH_2_ and 4-vinylbenzoic acid (4.8 mg, 32.6 μmol) to obtain 7.5 mg (21.2 μmol) of the desired product as a colorless oil in 72% yield.

**^1^H NMR** (400 MHz, DMSO-*d*_6_) *δ* [ppm] = 8.49 (t, *J* = 5.6 Hz, 1H), 7.83 (d, *J* = 8.2 Hz, 2H), 7.55 (d, *J* = 8.2 Hz, 2H), 6.78 (dd, *J* = 17.7, 10.9 Hz, 1H), 5.94 (d, *J* = 17.7 Hz, 1H), 5.36 (d, *J* = 10.9 Hz, 1H), 3.62 – 3.57 (m, 2H), 3.55 – 3.51 (m, 4H), 3.49 – 3.45 (m, 2H), 3.44 – 3.37 (m, 4H), 1.73 – 1.59 (m, 2H), 1.50 – 1.40 (m, 2H), 1.40 – 1.24 (m, 4H).

**^13^C NMR** (101 MHz, DMSO-*d*_6_) *δ* [ppm] = 166.28, 140.15, 136.39, 134.03, 127.99, 126.41, 116.59, 70.66, 70.11, 69.88, 69.33, 45.84, 39.67, 32.48, 29.55, 26.59, 25.39.

**HRMS** (ESI) calc. for C_19_H_28_ClNNaO_3_^+^ [M+Na]^+^: 376.1650; found 376.1640.

##### 1.3.6 ((1*R*,8*S*,9*s*)-bicyclo[6.1.0]non-4-yn-9-yl)methyl (2-(2-((6-chlorohexyl)oxy)ethoxy)ethyl)carbamate (CA-BCN)

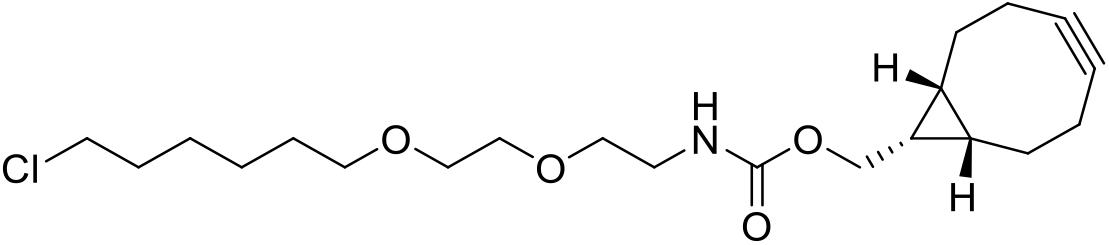

BCN-NHS (14.0 mg, 47.6 μmol, 1.1 eq) was dissolved in 500 μL DMSO. DIPEA (71.4 μL, 432 μmol, 10 equiv.) was added followed by CA-NH_2_ (14.0 mg, 43.2 μmol, 1.0 equiv.) solubilized in DMSO. The solution was stirred for 30 min. The crude product was purified by preparative HPLC eluted with MeCN / H_2_O (0.1% TFA) (50% - 90% MeCN over 60 min) to obtain 11.9 mg (29.8 μmol) of the product as a clear oil in 69% yield after lyophilization.

**^1^H NMR** (400 MHz, DMSO-d6): δ = 7.07 (t, J=5.7, 1H), 4.03 (d, J=8.0, 2H), 3.62 (t, J=6.6, 2H), 3.52 – 3.44 (m, 4H), 3.38 (dt, J=11.3, 6.3, 4H), 3.11 (q, J=6.0, 2H), 2.30 – 2.06 (m, 6H), 1.78 – 1.64 (m, 2H), 1.59 – 1.42 (m, 4H), 1.41 – 1.19 (m, 4H), 0.95 – 0.78 (m, 2H).

**^13^C NMR** (100 MHz, DMSO-*d*_6_): δ = 156.4, 99.0, 70.2, 69.5, 69.4, 69.1, 61.3, 45.4, 40.1, 32.0, 29.1, 28.6, 26.1, 24.9, 20.8, 19.5, 17.6.

**HRMS** (ESI) calc. for [M+H]^+^: 400.2249, found 400.2250.

###### 1.3.6.1 Cyclooct-2-yn-1-yl (2-(2-((6-chlorohexyl)oxy)ethoxy)ethyl)carbamate (CA-SCO)

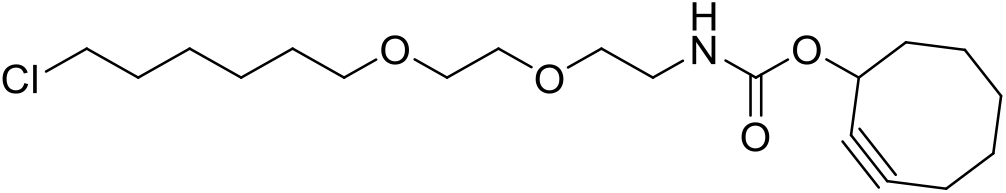

CA-NH_2_ (15 mg, 44.4 μmol, 1.3 equiv.) was dissolved in dry DMSO (0.15 mL) and a solution of cyclooct-2-yn-1-yl (4-nitrophenyl) carbonate (10 mg, 34.2 μmol, 1.0 equiv.) in dry DMF (0.4 mL) was added followed by DIPEA (58 μL, 348 μmol: 10.2 equiv.). The reaction mixture was stirred at r.t. for 1h. The resulted mixture was acidified with 50 μL of acetic acid and afterwards purified by semi-preparative HPLC eluted with MeCN / H_2_O (0.1% TFA) (15% MeCN for 2 min., then 15 - 100% MeCN over 25 min., followed by 100% MeCN for 15 min.) to give 8.7 mg (23.3 μmol) of the desired product as a colorless oil in 68% yield after lyophilization.

**^1^H NMR** (400 MHz, DMSO-*d*_6_) *δ* [ppm] = 7.18 (t, *J* = 5.9 Hz, 1H), 5.18 – 5.09 (m, 1H), 3.62 (t, *J* = 6.6 Hz, 2H), 3.50 – 3.43 (m, 4H), 3.40 – 3.34 (m, 4H), 3.09 (q, *J* = 5.9 Hz, 2H), 2.30 – 2.00 (m, 3H), 1.93 – 1.78 (m, 3H), 1.76 – 1.65 (m, 3H), 1.64 – 1.54 (m, 2H), 1.53 – 1.43 (m, 3H), 1.42 – 1.25 (m, 4H).

**^13^C NMR** (101 MHz, DMSO-*d*_6_) *δ* [ppm] = 155.29, 100.82, 91.79, 70.19, 69.53, 69.42, 68.99, 65.70, 45.38, 41.59, 40.07, 33.85, 32.03, 29.21,29.06, 26.13, 25.78, 24.94, 19.95.

**HRMS** (ESI) calc. for C_19_H_32_ClNNaO_4_^+^ [M+Na]^+^; 396.1912; found 396.1923.

##### 1.3.7 *N*-(2-(2-((6-chlorohexyl)oxy)ethoxy)ethyl)acetamide (CA-Ac)

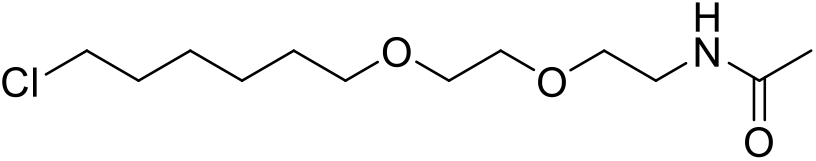

Tert-butyl (2-(2-((6-chlorohexyl)oxy)ethoxy)ethyl)carbamate (301 mg, 0.93 μmol, 1.0 equiv.) was deprotected by addition of TFA (2 mL) and afterwards dried under a stream of pressured air for 15 min. DIPEA (307 μL, 1.86 mmol, 2.0 equiv.) and DMSO (333 μL) were added followed by dropwise addition of acetic anhydride (131 μL, 1.39 mmol, 1.5 equiv.) while stirring. The reaction was stirred at r.t for 1 h. The mixture was quenched with saturated solution of NaHCO_3_ (20 mL) and extracted with DCM (3 × 20 mL). The combined organic layers were washed with brine and dried over MgSO4. All volatiles were evaporated and the crude product was purified over normal phase flash chromatography (MeOH: DCM = 2%: 98% to 3%: 97%). The fractions containing the product were combined to give 238 mg (896 μmol) of the desired product as a colorless oil in 97% yield after evaporation.

**^1^H NMR** (400 MHz, CDCl3) *δ* [ppm] = 6.05 (s, 1H), 3.67 – 3.38 (m, 12H), 1.98 (s, 3H), 1.83 – 1.71 (m, 2H), 1.61 (p, *J* = 6.8 Hz, 2H), 1.52 – 1.31 (m, 4H).

**^13^C NMR** (101 MHz, CDCl3) *δ* [ppm] = 169.92, 71.09, 70.07, 69.83, 69.60, 44.84, 39.10, 32.32, 29.28, 26.49, 25.24, 23.10.

**HRMS** (ESI) calc. for C1_2_H_25_ClNO_3_^+^ [M+H]^+^: 266.1517; found 266.1518.

##### 1.3.1 5-((2-(2-((6-chlorohexyl)oxy)ethoxy)ethyl)carbamoyl)-2-(6-(dimethylamino)-3-(dimethyliminio)-3H-xanthen-9-yl)benzoate (CA-5-TMR)

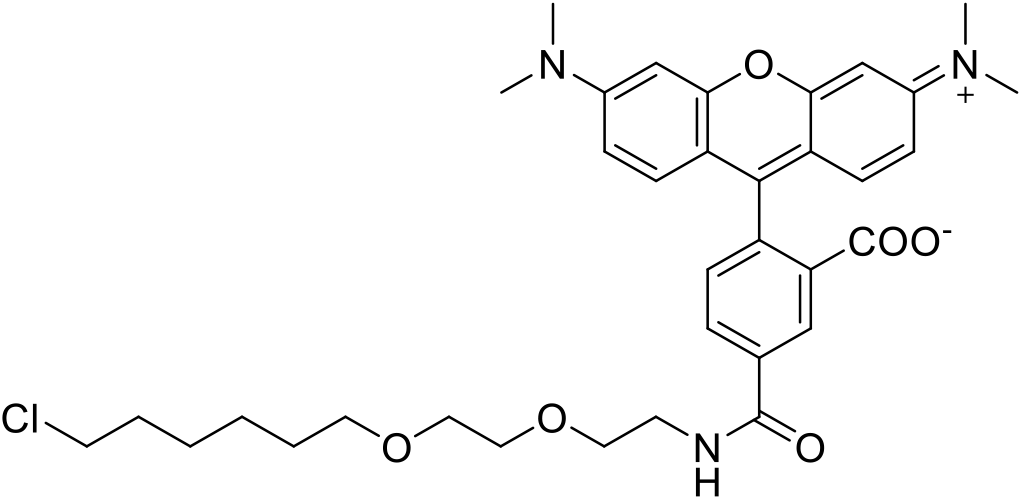

To a solution of TMR-5-COOH (2.5 mg, 5.81 μmol, 1.0 equiv.) in dry DMSO (500 μL), benzotriazolyloxytris(dimethylamino)-phosphonium hexafluorophosphat (BOP) (0.5 M in DMSO, 16.4 μL, 8.21 μmol, 1.5 equiv.) was added and the reaction was shaken at 500 rpm and r.t. for 5 min. DIPEA (3.84 μL, 23.2 μmol, 4.0 equiv.) and CA-NH_2_ (1 M in DMSO, 8.71 μL, 8.71 μmol, 1.5 equiv.) were added and the reaction was shaken at 500 rpm and r.t. for 4 h. The crude product was acidified with acetic acid and purified over preparative HPLC eluted with MeCN / H_2_O (0.1% FA) (10% - 90% MeCN over 50 min) to give 1.2 mg (1.89 μmol) of the desired product in 33% yield after lyophilization.

**HRMS** (ESI): calc. for C_36_H_44_N_2_O_6_Cl^+^ [M+H]^+^: 635.2887; found 635.2882.

##### 1.3.2 5-((2-(2-((6-chlorohexyl)oxy)ethoxy)ethy1)carbamoy1)-2-(6-(dimethylamino)-3-(dimethyliminio)-10,10-dimethyl-3,10-dihydroanthracen-9-yl)benzoate (CA-5-CPY)

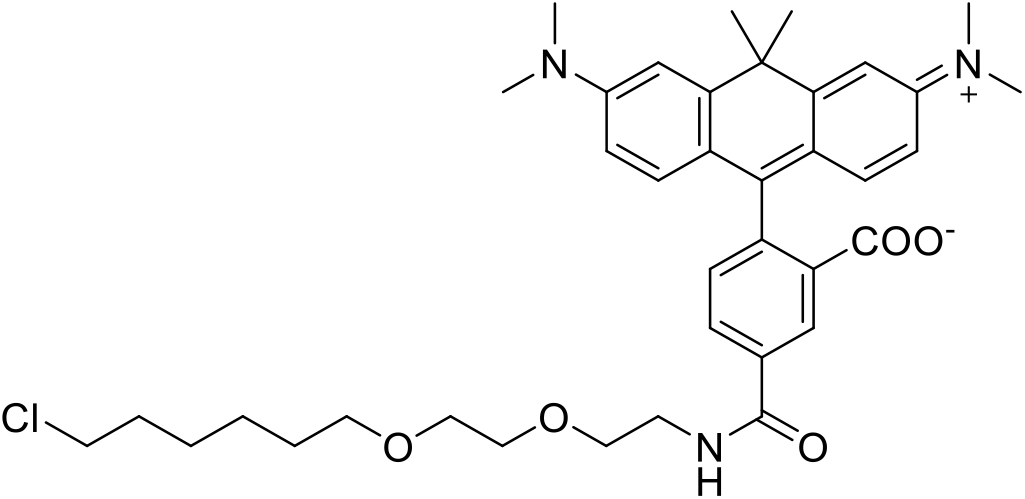

To a solution of CPY-5-COOH (2.5 mg, 5.48 μmol, 1.0 equiv.) in dry DMSO (1 mL), BOP (0.5 M in DMSO, 16.4 μL, 8.21 μmol, 1.5 equiv.) was added and the reaction was shaken at 500 rpm and r.t. for 5 min. DIPEA (3.62 μL, 21.9 μmol, 4.0 equiv.) and CA-NH_2_ (1 M in DMSO, 8.21 μL, 8.21 μmol, 1.5 equiv.) were added and the reaction was shaken at 500 rpm and r.t. for 4 h. The crude product was acidified with acetic acid and purified over preparative HPLC eluted with MeCN / H_2_O (0.1% FA) (10% - 90% MeCN over 50 min) to give 0.38 mg (0.57 μmol) of the desired product in 10% yield after lyophilization.

**HRMS** (ESI): calc. for C_38_H_49_N_3_O_5_Cl^+^ [M+H]^+^: 662.3360; found 662.3349.

##### 1.3.3 1-(6-((2-(2-((6-chlorohexyl)oxy)ethoxy)ethyl)amino)-6-oxohexyl)-3,3-dimethyl-2-((E)-3-((Z)-1,3,3-trimethylindolin-2-ylidene)prop-1-en-1-yl)-3H-indol-1-ium (CA-Cy3)

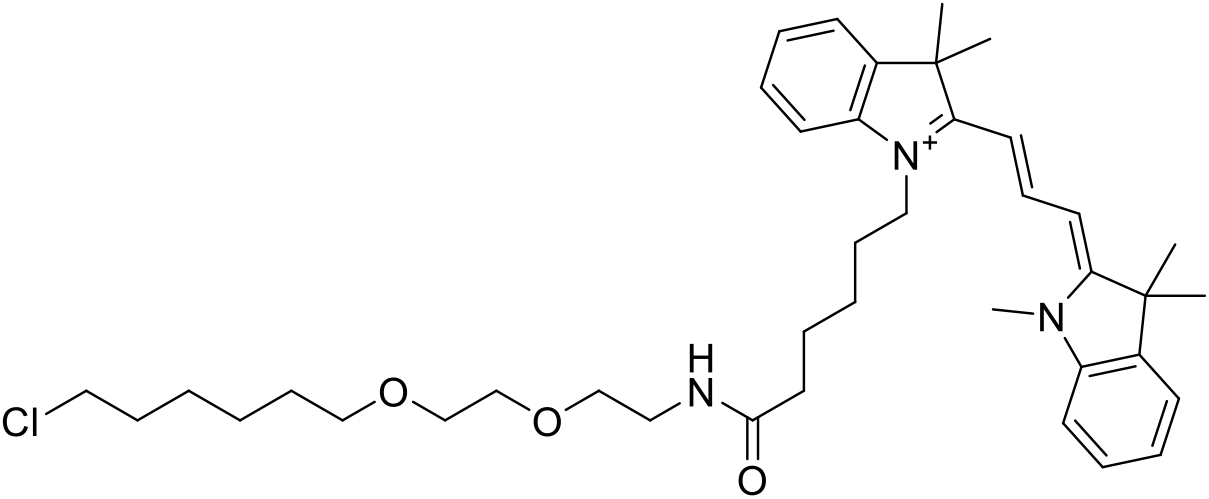

Cy3-COOH was synthesized according to Ueno et al. 2010 (3). To a solution of Cy3-COOH (100 mg, 219 μmol, 1.0 equiv.) in dry DMSO (2 mL), DIPEA (217 μL, 1.3 mmol, 6.0 equiv.) and TSTU (92.1 mg, 306 μmol, 1.4 equiv.) were added and the reaction mixture was stirred for 10 min. at r.t. CA-NH_2_ (58 mg, 262 μmol, 1.2 equiv.) in 0.5 mL DMSO was added and the reaction was stirred for 30 min, at r.t. The reaction was quenched by addition of acetic acid (230 μL) and 10% H_2_O, followed by purification over preparative HPLC eluted with MeCN / H_2_O (0.1% FA) (10% - 90% MeCN over 60 min) to give 102 mg (154 μmol) of the desired product in 70% yield after lyophilization.

**HRMS** (ESI): calc. for C_40_H_57_N_3_O_3_Cl^+^ [M]^+^: 662.4083; found 662.4084.

##### 1.3.4 1-(6-((2-(2-((6-chlorohexyl)oxy)ethoxy)ethyl)amino)-6-oxohexyl)-3,3-dimethyl-2-((1 E,3E)-5-((Z)-1,3,3-trimethylindolin-2-ylidene)penta-1,3-dien-1-yl)-3H-indol-1-ium (CA-Cy5)

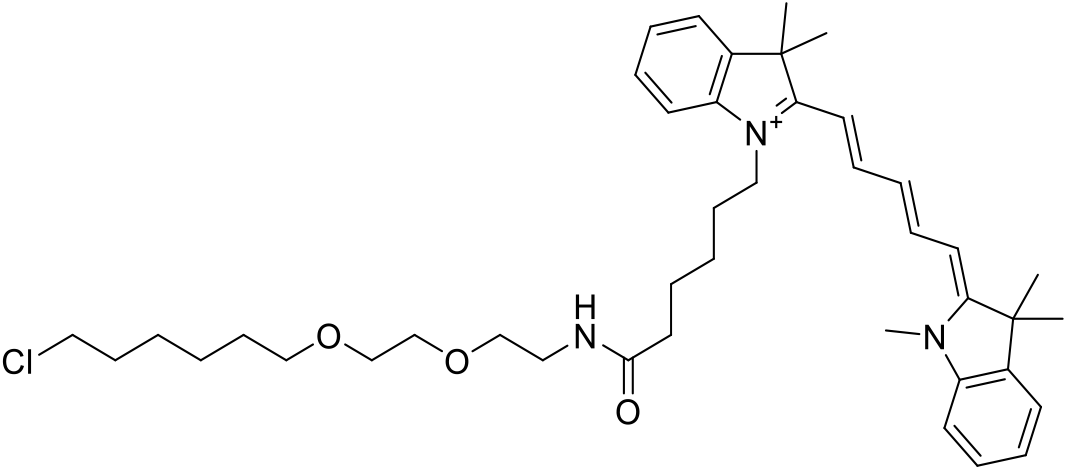

Cy5-COOH was synthesized according to Ueno et al. 2010 (3). To a solution of Cy5-COOH (100 mg, 207 μmol, 1.0 equiv.) in dry DMSO (2 mL), DIPEA (205 μL, 1.24 mmol, 6.0 equiv.) and TSTU (87.1 mg, 289 μmol, 1.4 equiv.) were added and the reaction mixture was stirred for 10 min. at r.t. CA-NH_2_ (55.5 mg, 248 μmol, 1.2 equiv.) in 0.5 mL DMSO was added and the reaction was stirred for 30 min, at r.t. The reaction was quenched by addition of acetic acid (291 μL) and 10% H_2_O, followed by purification over preparative HPLC eluted with MeCN / H_2_O (0.1% FA) (10% - 90% MeCN over 60 min) to give 98 mg (142 μmol) of the desired product in 69% yield after lyophilization.

**HRMS** (ESI): calc. for C_42_H_59_N_3_O_3_Cl^+^ [M]^+^: 688.4239; found 688.4239.

##### 1.3.5 4-carboxy-2-(3-(dimethyliminio)-6-((4-methoxy-4-oxobutyl)(methyl)amino)-3H-xanthen-9-yl)benzoate (CA-TMR-biotin-1)

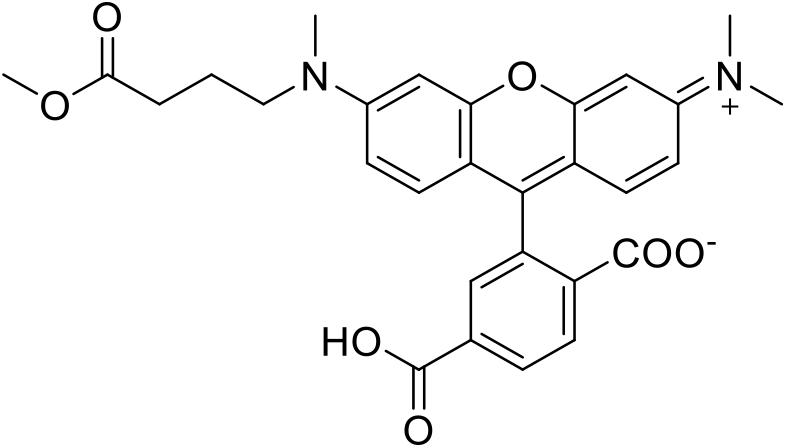

The compound was synthesized according to the procedure from Masharina et al. 2012 (15).

##### 1.3.6 4-((2-(2-((6-chlorohexyl)oxy)ethoxy)ethyl)carbamoyl)-2-(3-(dimethyliminio)-6-((4-methoxy-4-oxobutyl)(methyl)amino)-3H-xanthen-9-yl)benzoate (CA-TMR-biotin-2)

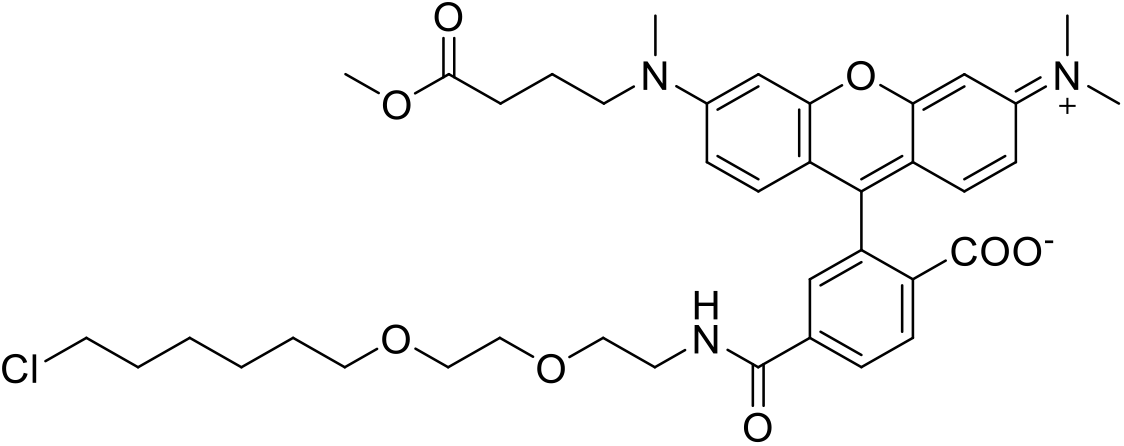

To a solution of CA-TMR-biotin-1 (17.0 mg, 32.9 μmol, 1.0 equiv.) in dry DMF, TSTU (11.9 mg, 39.5 μmol, 1.2 equiv.) and DIPEA (32.6 μL, 197 μmol, 6.0 equiv.) were added and the reaction was stirred at r.t. for 5 min. CA-NH_2_ (14.7 mg, 65.8 μmol, 2.0 equiv.) was added and the reaction was stirred at r.t. for 2 h. The crude product was acidified with acetic acid and purified via preparative eluted with MeCN / H_2_O (0.1% TFA) (10% - 90% MeCN over 50 min) to give 10 mg (13.8 μmol) of the desired product in 42% yield after lyophilization.

**HRMS** (ESI): calc. for C_39_H_49_N_3_O_8_Cl^+^ [M+H]^+^: 722.3208; found 722.3202.

##### 1.3.7 2-(6-((3-carboxypropyl)(methyl)amino)-3-(dimethyliminio)-3H-xanthen-9-yl)-4-((2-(2-((6-chlorohexyl)oxy)ethoxy)ethyl)carbamoyl)benzoate (CA-TMR-biotin-3)

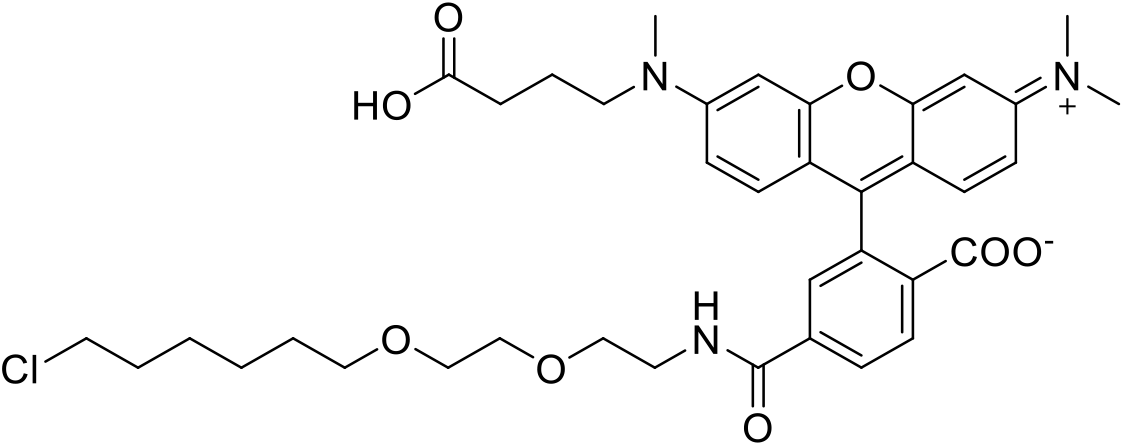

To a solution of CA-TMR-biotin-2 (8.0 mg, 11.1 μmol, 1.0 equiv.) in THF:H_2_O (4:1), lithium hydroxide (1M in H_2_O, 22.2 μL, 22.2 μmol, 2.0 equiv.) was added and the reaction was stirred at r.t. for 6 h. The crude product was acidified with acetic acid and purified via preparative HPLC eluted with MeCN / H_2_O (0.1% TFA) (10% - 90% MeCN over 50 min) to give 6.3 mg (8.9 μmol) of the desired product in 80% yield after lyophilization.

**HRMS** (ESI): calc. for C_39_H_49_N_3_O_8_Cl^+^ [M+H]^+^: 708.3051; found 708.3049.

##### 1.3.8 4-((2-(2-((6-chlorohexyl)oxy)ethoxy)ethyl)carbamoyl)-2-(3-(dimethyliminio)-6-((4,18-dioxo-22-((3aR,4R,6aS)-2-oxohexahydro-1H-thieno[3,4-d]imidazol-4-yl)-8,11,14-trioxa-5,17-diazadocosyl)(methyl)amino)-3H-xanthen-9-yl)benzoate (CA-TMR-biotin)

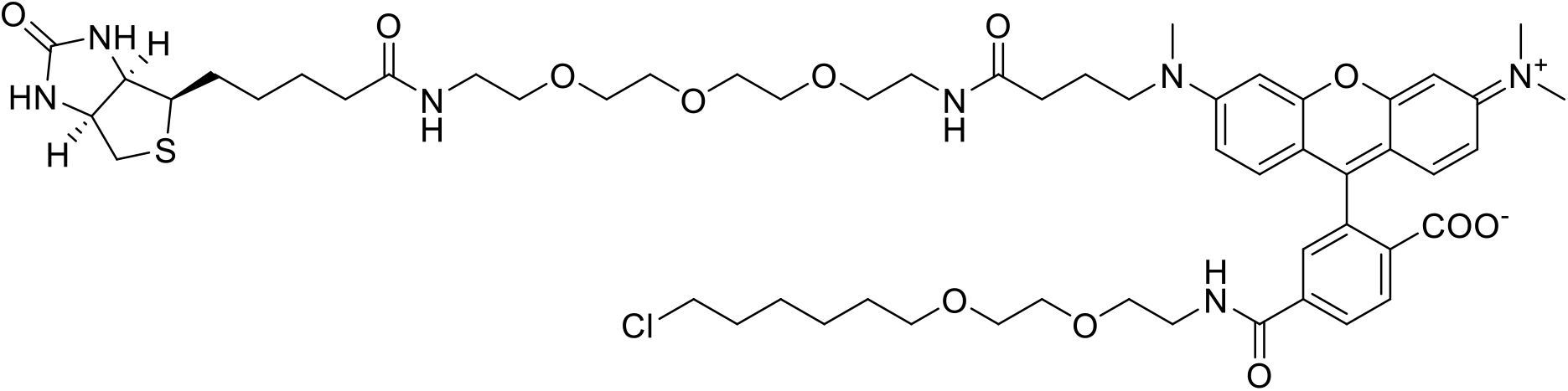

To a solution of CA-TMR-biotin-3 (6.0 mg, 8.47 μmol, 1.0 equiv.) in dry DMF, TSTU (3.06 mg, 10.2 μmol, 1.2 equiv.) and DIPEA (8.4 μL, 50.8 μmol, 6.0 equiv.) were added and the reaction was stirred at r.t. for 5 min. Biotin-PEG3-NH_2_ (7.09 mg, 16.9 μmol, 2.0 equiv.) was added and the reaction was stirred at r.t. for another 2 h. The crude product was acidified with acetic acid and purified via preparative HPLC eluted with MeCN / H_2_O (0.1% TFA) (10% - 90% MeCN over 50 min) to give 6.2 mg (5.6 μmol) of the desired product in 66% yield after lyophilization.

**HRMS** (ESI): calc. for C_56_H_80_N_7_O_12_ClS^2+^ [M+2H]^2+^: 554.7628; found 554.7632.

#### 1.4 SNAP substrates based on benzylguanine (BG)

##### 1.4.1 *N*-(4-(((2-amino-9*H*-purin-6-yl)oxy)methyl)benzyl)-2-azidoacetamide (BG-N_3_)

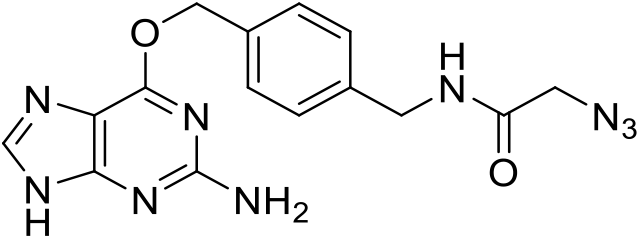

Reaction was conducted according to general procedure A using BG-NH_2_ and 2-azidoacetic acid (40.7 μmol; 5.7 μL) and 11.1 mg (23.8 μmol) of the desired product were obtained as a colorless TFA-salt in 64% yield.

**^1^H NMR** (400 MHz, DMSO-*d*_6_) *δ* [ppm] = 8.65 (t, *J* = 5.9 Hz, 1H), 8.34 (s, 1H), 7.55 – 7.44 (m, 2H), 7.36 – 7.28 (m, 2H), 5.52 (s, 2H), 4.31 (d, *J* = 5.9 Hz, 2H), 3.89 (s, 2H).

**HRMS** (ESI) calc. for C_15_H_16_N_9_O_2_^+^ [M+H]^+^: 354.1421; found 354.1423.

##### 1.4.2 *N*-(4-(((2-amino-9*H*-purin-6-yl)oxy)methyl)benzyl)-2-((1 *S*, 4*S*)-bicyclo[2.2.1]hept-5-en-2-yl)acetamide (BG-Nor2)

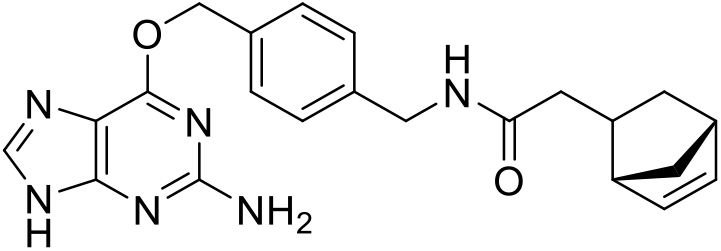

Reaction was conducted according to general procedure A with a reduced reaction time of 15 min. using BG-NH_2_ and 2-((1*S*,4*S*)- bicyclo[2.2.1]hept-5-en-2-yl)acetic acid (40.7 μmol; 7.0 μL) resulting in 15.9 mg (30.7 μmol) of the desired product as a colorless TFA-salt in 83% yield.

**^1^H NMR** (400 MHz, DMSO-*d*_6_) *δ* [ppm] = 8.47 (s, 1H), 8.29 (t, *J* = 6.0 Hz, 1H), 7.49 (d, *J* = 8.1 Hz, 2H), 7.27 (d, *J* = 8.1 Hz, 2H), 6.16 (dd, *J* = 5.7, 3.0 Hz, 1H), 5.96 (dd, *J* = 5.7, 2.9 Hz, 1H), 5.53 (s, 2H), 4.25 (d, *J* = 6.0 Hz, 2H), 2.77 – 2.69 (m, 2H), 2.45 – 2.34 (m, 1H), 1.95 (dd, *J* = 13.8, 7.6 Hz, 1H), 1.90 – 1.77 (m, 2H), 1.33 – 1.26 (m, 1H), 1.25 – 1.19 (m, 1H), 0.50 (m, *J* = 11.4, 4.5, 2.5 Hz, 1H).

**^13^C NMR** (101 MHz, DMSO-*d*_6_) *δ* [ppm] = 171.71, 158.83, 158.03, 153.44, 140.89, 140.30, 137.07, 133.90, 132.45, 128.84, 127.13, 68.12, 49.10, 45.26, 42.09, 41.71, 40.67, 35.15, 31.47.

**HRMS** (ESI) calc. for C_22_H_25_N_6_O_2_^+^ [M+H]^+^: 405.2034; found 405.2034.

##### 1.4.3 *N*-(4-(((2-amino-9*H*-purin-6-yl)oxy)methyl)benzyl)-2-(4-(6-methyl-1,2,4,5-tetrazin-3-yl)phenyl)acetamide (BG-Tz)

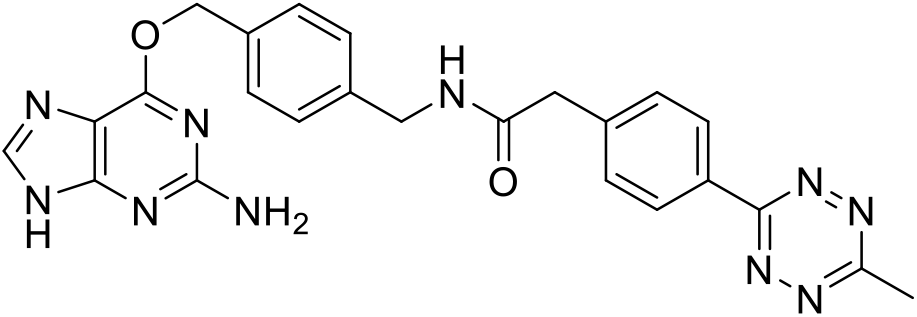

Reaction was conducted according to general procedure A using BG-NH_2_ and 2-(4-(6-methyl-1,2,4,5-tetrazin-3-yl)phenyl)acetic acid (40.7 μmol, 9.4 mg) to give 12.4 mg (17.0 μmol) of the desired product as a rose TFA-salt in 56% yield.

**^1^H NMR** (400 MHz, DMSO-*d*_6_) *δ* [ppm] = 8.70 (t, *J* = 5.9 Hz, 1H), 8.45 – 8.39 (m, 2H), 8.37 (s, 1H), 7.60 – 7.53 (m, 2H), 7.52 – 7.44 (m, 2H), 7.33 – 7.26 (m, 2H), 5.51 (s, 2H), 4.31 (d, *J* = 5.9 Hz, 2H), 3.64 (s, 2H), 2.99 (s, 3H), 2.54 (s, 3H).

**^13^C NMR** (101 MHz, DMSO-*d*_6_) *δ* [ppm] = 170.04, 167.53, 163.69, 159.37, 158.85, 158.53, 154.36, 141.59, 140.96, 140.84, 140.19, 134.78, 130.62, 130.61, 129.34, 127.81, 68.28, 42.69, 40.90, 21.31.

**HRMS** (ESI) calc. for C_24_H_23_N_10_O_2_^+^ [M+H]^+^: 483.2000; found 483.2006.

##### 1.4.4 *N*-(4-(((2-amino-9*H*-purin-6-yl)oxy)methyl)benzyl)-4-azidobenzamide (BG-PhN_3_)

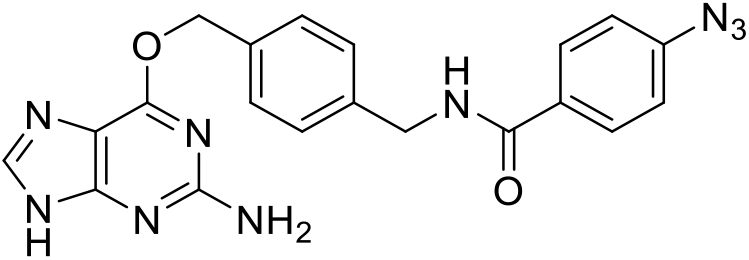

Reaction was conducted according to general procedure A with a reduced reaction time of 15 min. using BG-NH_2_ and 4-azidobenzoic acid (6.6 mg, 40.7 μmol) to obtain 15.5 mg (29.3 μmol) of the desired product as a colorless TFA-salt in 79% yield.

**^1^H NMR** (400 MHz, DMSO-*d*_6_) *δ* [ppm] = 9.10 (t, *J* = 5.9 Hz, 1H), 8.38 (s, 1H), 7.99 – 7.89 (m, 2H), 7.55 – 7.46 (m, 2H), 7.38 – 7.33 (m, 2H), 7.25 – 7.16 (m, 2H), 5.52 (s, 2H), 4.48 (d, *J* = 5.9 Hz, 2H).

**^13^C NMR** (101 MHz, DMSO-*d*_6_) *δ* [ppm] = 165.27, 158.91, 158.61, 158.26, 153.77, 142.36, 140.57, 140.02, 134.19, 130.81, 129.15, 128.89, 127.36, 118.96, 67.94, 42.48.

**HRMS** (ESI) calc. for C_20_H_18_N_9_O_2_^+^ [M+H]^+^: 416.1578; found 416.1577.

##### 1.4.5 *N*-(4-(((2-amino-9*H*-purin-6-yl)oxy)methyl)benzyl)-4-vinylbenzamide (BG-VBn)

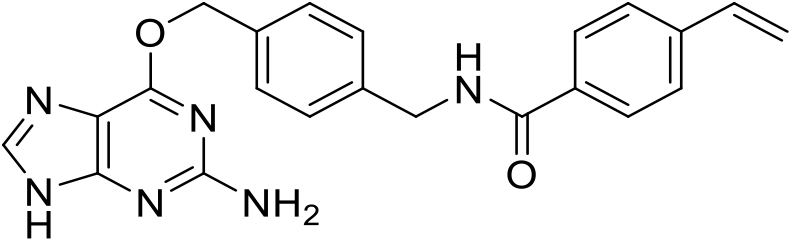

Reaction was conducted according to general procedure A with a reduced reaction time of 15 min. using BG-NH_2_ and 4-vinylbenzoic acid (40.7 μmol; 6.5 mg) to obtain 14.7 mg (28.6 μmol) of the desired product as a colorless TFA-salt in 77% yield.

**^1^H NMR** (400 MHz, DMSO-*d*_6_) *δ* [ppm] = 9.09 (t, *J* = 6.0 Hz, 1H), 8.44 (s, 1H), 7.87 (d, *J* = 8.3 Hz, 2H), 7.57 (d, *J* = 8.3 Hz, 2H), 7.51 (d, *J* = 7.9 Hz, 2H), 7.36 (d, *J* = 7.9 Hz, 2H), 6.79 (dd, *J* = 17.7, 11.0 Hz, 1H), 5.95 (d, *J* = 17.7 Hz, 1H), 5.53 (s, 2H), 5.37 (d, *J* = 11.0 Hz, 1H), 4.49 (d, *J* = 6.0 Hz, 2H).

**^13^C NMR** (101 MHz, DMSO-*d*_6_) *δ* [ppm] = 165.81, 158.85, 158.62, 158.28, 153.55, 140.79, 140.13, 139.84, 135.91, 134.06, 133.43, 128.92, 127.61, 127.35, 126.03, 116.24, 68.09, 42.46.

**HRMS** (ESI) calc. for C_22_H_21_N_6_O_2_^+^ [M+H]^+^: 401.1721; found 401.1707.

##### 1.4.6 ((1R,8S,9s)-bicyclo[6.1.0]non-4-yn-9-yl)methyl (4-(((2-amino-9*H*-purin-6-yl)oxy)methyl)benzyl)carbamate (BG-BCN)

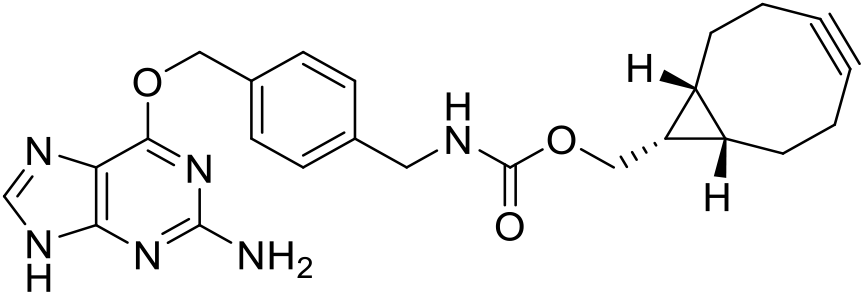

A solution of ((1*R*,8*S*,9*s*)-bicyclo[6.1.0]non-4-yn-9-yl)methyl (2,5-dioxopyrrolidin-1-yl) carbonate (10 mg, 34.3 μmol, 1.0 equiv.) in dry DMSO (0.4 mL) was added to a solution of 10.2 mg BG-NH_2_ (37.8 μmol, 1.1 equiv.) in dry DMSO (0.1 mL) followed by 28.4 μL of DIPEA (172 μmol, 5 equiv.). The reaction was stirred at r.t. for 30 min. The resulted mixture was acidified with acetic acid (3 μL) and H_2_O (53 μL), then purified by semi-preparative HPLC eluted with MeCN / H_2_O (0.1% TFA) (10% MeCN for 10 min., then 10 - 90% MeCN over 55 min. followed by 99% MeCN for 5 min.) to give 14.0 mg (31.4 μmol) of the desired product as a colorless solid in 91% yield after lyophilization.

**^1^H NMR** (400 MHz, DMSO-*d*_6_) *δ* [ppm] = 8.36 (s, 1H), 7.70 (t, *J* = 6.2 Hz, 1H), 7.52 – 7.46 (m, 2H), 7.28 (d, *J* = 8.0 Hz, 2H), 5.51 (s, 2H), 4.18 (d, *J* = 6.0 Hz, 2H), 4.06 (d, *J* = 8.0 Hz, 2H), 2.28 – 2.07 (m, 6H), 1.52 (d, *J* = 12.4 Hz, 2H), 1.28 (dt, *J* = 18.3, 9.1 Hz, 1H), 0.86 (t, *J* = 9.8 Hz, 2H).

**HRMS** (ESI) calc. for C_24_H_27_N_6_O_3_^+^ [M+H]^+^: 447.2139; found 447.2135.

###### 1.4.6.1 *N*-(4-(((2-amino-9*H*-purin-6-yl)oxy)methyl)benzyl)acetamide (BG-Ac)

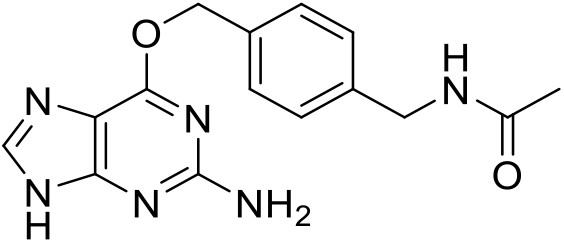

BG-NH_2_ (300 mg, 1.11 mmol, 1.0 equiv.) was dissolved in dry DMSO (2.5 mL) and 367 μL of DIPEA (2.22 mmol, 2.0 equiv.) was added followed by dropwise addition of acetic anhydride (156 μL, 1.66 mmol, 1.5 equiv.) while stirring. The reaction mixture was stirred at r.t. for 1 h. Afterwards, the reaction was quenched with acetic acid (387 μL) and H_2_O (341 μL) followed by centrifugation at 3’000 rpm for 3 min. The pellet was washed twice with H_2_O and afterwards lyophilized to obtain 190 mg (608 μmol) of the desired product as a colorless solid in 55% yield.

**^1^H NMR** (400 MHz, DMSO-*d*_6_) *δ* [ppm] = 8.30 (s, 1H), 7.45 (d, *J* = 8.1 Hz, 2H), 7.27 (d, *J* = 7.9 Hz, 2H), 6.82 (s, 2H), 5.48 (s, 2H), 4.24 (d, *J* = 5.9 Hz, 2H), 1.86 (s, 3H).

**HRMS** (ESI) calc. for C_15_H_17_N_6_O_2_^+^ [M+H]^+^: 313.1408; found 313.1406.

##### 1.4.7 Cyclooct-2-yn-1-yl (4-(((2-amino-9*H*-purin-6-yl)oxy)methyl)benzyl)carbamate (BG-SCO)

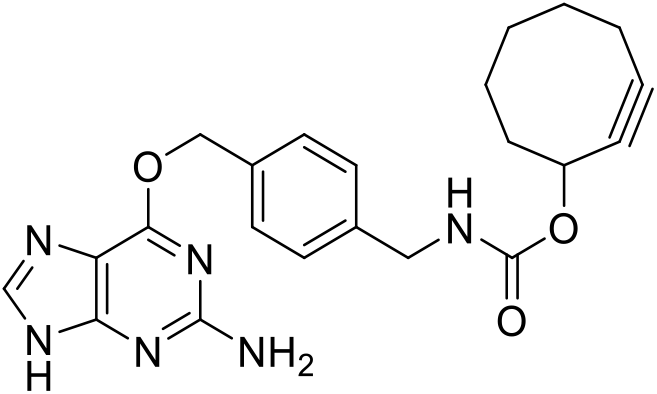

BG-NH_2_ (10 mg, 37.0 μmol, 1.0 equiv.) was dissolved in dry DMSO (0.5 mL) and a solution of cyclooct-2-yn-1-yl (4-nitrophenyl) carbonate (10.7 mg, 37 μmol, 1.0 equiv.) in dry DMF (0.4 mL) was added followed by DIPEA (30.6 μL, 142 μmol: 5.0 equiv.). The reaction mixture was stirred at r.t. for 1 h. The resulted mixture was acidified with acetic acid (25 μL) and afterwards purified by semi-preparative HPLC eluted with MeCN / H_2_O (0.1% TFA) (15% MeCN for 5 min., then 15 - 100% MeCN over 25 min., followed by 100% MeCN for 15 min.) to give 16 mg (23.3 μmol) of the desired product as a colorless TFA-salt in 81% yield after lyophilization.

**^1^H NMR** (400 MHz, DMSO-*d*_6_) *δ* [ppm] = 8.44 (s, 1H), 7.81 (t, *J* = 6.1 Hz, 1H), 7.50 (d, *J* = 8.0 Hz, 2H), 7.28 (d, *J* = 8.0 Hz, 2H), 5.52 (s, 2H), 5.20 – 5.11 (m, 1H), 4.17 (d, *J* = 6.1 Hz, 2H), 2.30 – 2.02 (m, 3H), 1.96 – 1.85 (m, 1H), 1.89 – 1.76 (m, 2H), 1.76 – 1.64 (m, 1H), 1.64 – 1.53 (m, 2H), 1.55 – 1.41 (m, 1H).

**^13^C NMR** (101 MHz, DMSO-*d*_6_) *δ* [ppm] = 158.86, 158.10, 155.53, 153.58, 140.80, 140.06, 134.15, 128.90, 127.16, 107.66, 100.97, 91.76, 68.06, 65.95, 43.50, 41.58, 33.85, 29.21, 25.79, 19.95.

**HRMS** (ESI) calc. for C_22_H_24_N_6_NaO_3_^+^ [M+Na]^+^: 443.1802; found 443.1797.

##### 1.4.8 5-((4-(((2-amino-9H-purin-6-yl)oxy)methyl)benzyl)carbamoyl)-2-(6-(dimethylamino)-3-(dimethyliminio)-2,3,4,4a-tetrahydro-1H-xanthen-9-yl)benzoate (BG-5-TMR)

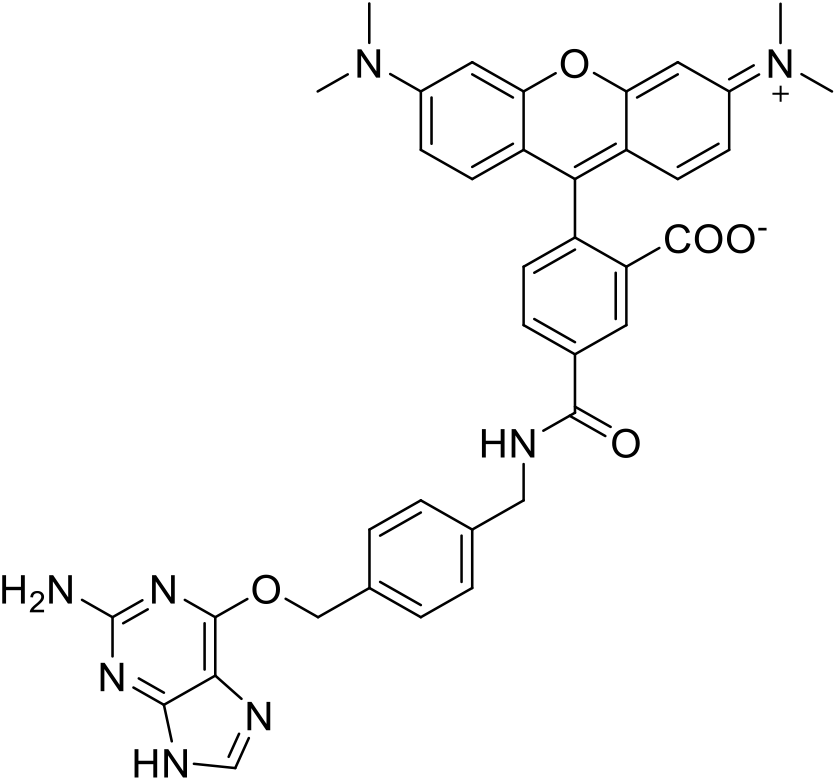

TSTU (1.45 mg, 4.82 μmol, 1.2 equiv.) was dissolved in dry DMSO-*d*_6_ (500 μL). TMR-5-COOH (1.15 mg, 2.68 μmol, 1.0 equiv.) was dissolved in the TSTU solution and DIPEA (1.77 μL, 10.7 μmol, 4.0 equiv.) was added. The mixture was stirred at r.t. for 10 min. BG-NH_2_ (1.08 mg, 4.01 μmol, 1.5 equiv.) was dissolved in dry DMSO-*d*_6_ (200 μL) and added to the reaction. The reaction mixture was stirred at r.t. for 1 h. The compound was purified over preparative HPLC eluted with MeCN / H_2_O (0.1% TFA) (10% MeCN for 10 min., then 10 - 90% MeCN over 40 min., followed by 90% MeCN for 5 min.) to give after lyophilization 378 μg (554 nmol) of the desired product in 21% yield.

**HRMS** (ESI): calc. for C_38_H_37_N_8_O_5_ [M+2H]^2+^: 342.1399; found 342.1394.

**^1^H NMR** (TMR-5-COOH) (400 MHz, DMSO-*d*_6_) *δ* [ppm] = 8.39 (s, *J* = 1.5 Hz, 1H), 8.28 (dd, *J* = 8.1, 1.5 Hz, 1H), 7.33 (d, *J* = 8.0 Hz, 1H), 6.58 – 6.45 (m, 6H), 2.95 (s, 12H).

**^13^C NMR** (TMR-5-COOH) (101 MHz, DMSO-*d*_6_) *δ* [ppm] = 168.31, 166.09, 152.03, 135.96, 132.76, 128.50, 109.05, 97.95, 40.15, 39.99, 39.79.

##### 1.4.9 5-((4-(((2-amino-9H-purin-6-yl)oxy)methyl)benzyl)carbamoyl)-2-(6-(dimethylamino)-3-(dimethyliminio)-10,10-dimethyl-1,2,3,4,4a,10-hexahydroanthracen-9-yl)benzoate (BG-5-CPY)

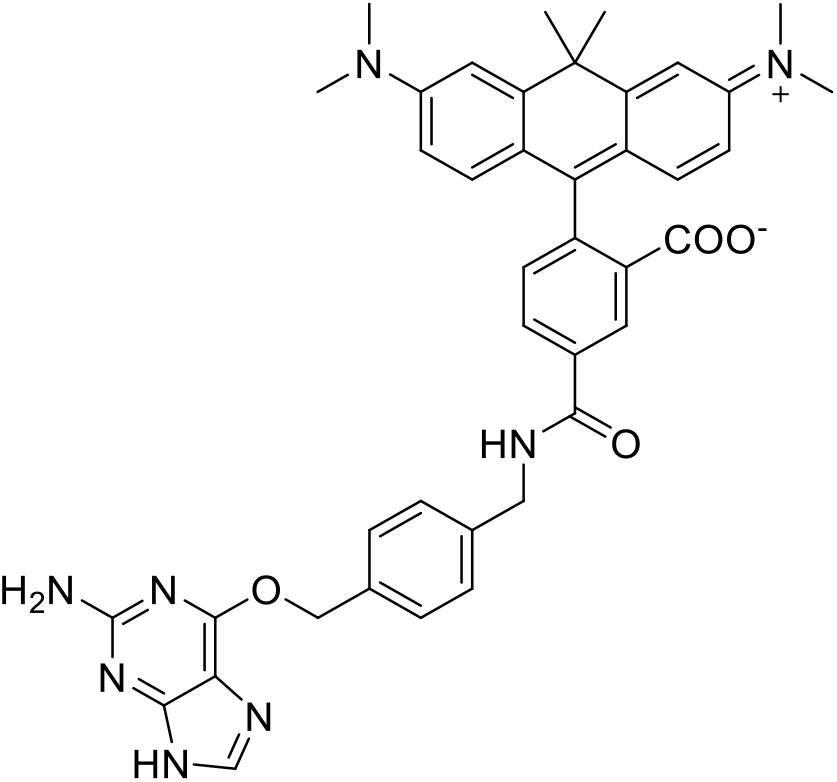

TSTU (1.44 mg, 4.78 μmol, 1.2 equiv.) was dissolved in dry DMSO-*d_6_* (500 μL). CPY-5-COOH (2.0 mg, 4.38 μmol, 1.1 equiv.) was dissolved in the TSTU solution and DIPEA (2.63 μL, 15.9 μmol, 4 equiv.) was added. The mixture was stirred at r.t. for 10 min. BG-NH2 (1.08 mg, 3.98 μmol, 1.5 equiv.) was dissolved in dry DMSO-*d*_6_ (200 μL) and added to the reaction. The reaction mixture was stirred at r.t. for 1 h. The compound was purified over preparative HPLC eluted with MeCN / H_2_O (0.1% TFA) (10% MeCN for 10 min., then 10 - 90% MeCN over 40 min., followed by 90% MeCN for 5 min.) to give 346 μg (488 nmol) of the desired product in 18% yield after lyophilization.

**HRMS** (ESI): calc. for C_41_H_42_N_8_O_4_ [M+2H]^2+^: 355.1659; found 355.1659.

###### 1.4.9.1 1-(6-((4-(((2-amino-9H-purin-6-yl)oxy)methyl)benzyl)amino)-6-oxohexyl)-3,3-dimethyl-2-((E)-3-((Z)-1,3,3-trimethylindolin-2-ylidene)prop-1-en-1-yl)-3H-indol-1-ium (BG-Cy3)

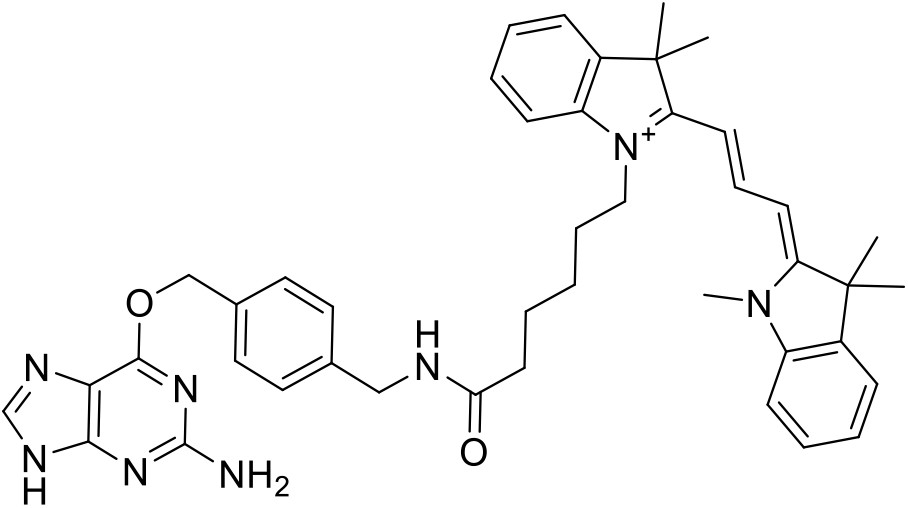

Cy3-COOH was synthesized according to Ueno et al. 2010 (3). To a solution of Cy3-COOH (100 mg, 219 μmol, 1.0 equiv.) in dry DMSO (1.5 mL), DIPEA (217 μL, 1.3 mmol, 6.0 equiv.) and TSTU (92.1 mg, 306 μmol, 1.4 equiv.) were added and the reaction mixture was stirred for 10 min. at r.t. BG-NH_2_ (70.9 mg, 262 μmol, 1.2 equiv.) was added and the reaction was stirred for 30 min. at r.t. The reaction was quenched by addition of acetic acid (230 μL) and 10% H_2_O, followed by purification over preparative HPLC eluted with MeCN / H_2_O (0.1% FA) (10% - 90% MeCN over 60 min.) to give. 28.5 mg (40.1 μmol) of the desired product in 18% yield after lyophilization.

**HRMS** (ESI): calc. for C_43_H_50_N_8_O_2_^2+^ [M+H]^2+^: 355.2023; found 355.2022.

##### 1.4.10 1-(6-((4-(((2-amino-9H-purin-6-yl)oxy)methyl)benzyl)amino)-6-oxohexyl)-3,3-dimethyl-2-((1E, 3E)-5-((Z)-1,3,3-trimethylindolin-2-ylidene)penta-1,3-dien-1-yl)-3H-indol-1-ium (BG-Cy5)

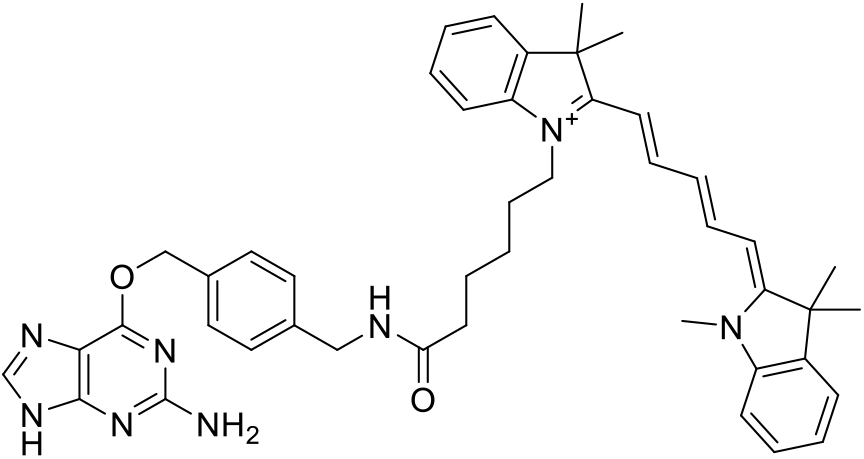

Cy5-COOH was synthesized according to Ueno et al. 2010(3). To a solution of Cy5-COOH (50.0 mg, 103 μmol, 1.0 equiv.) in dry DMSO (1.5 mL), DIPEA (103 μL, 620 μmol, 6.0 equiv.) and TSTU (43.6 mg, 145 μmol, 1.4 equiv.) were added and the reaction mixture was stirred for 10 min. at r.t. BG-NH_2_ (33.5 mg, 124 μmol, 1.2 equiv.) was added and the reaction was stirred for 30 min. at r.t. The reaction was quenched by addition of acetic acid (109 μL) and 10% H_2_O, followed by purification over preparative HPLC eluted with MeCN / H_2_O (0.1% FA) (10% - 90% MeCN over 60 min.) to give 45 mg (61.1 μmol) of the desired product in 59% yield after lyophilization.

**HRMS** (ESI): calc. for C_45_H_52_N_8_O_2_^2+^ [M+H]^2+^: 368.2101; found 368.2102.

#### 1.5 SNAP substrates based on chloropyrimidine (CP)

##### 1.5.1 *N*-(4-(((2-amino-6-chloropyrimidin-4-yl)oxy)methyl)benzyl)-2-azidoacetamide (CP-N_3_)

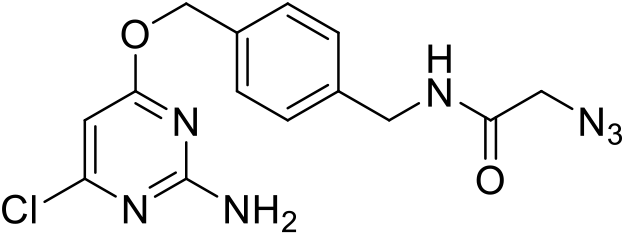

Reaction was conducted according to general procedure A using CP-NH_2_ and 2-azidoacetic acid (5.8 μL, 41.6 μmol) to obtain 10.1 mg (21.9 μmol) of the desired product as a colorless TFA-salt in 58% yield.

**^1^H NMR** (400 MHz, DMSO-*d*_6_) *δ* [ppm] = 8.62 (t, *J* = 5.8 Hz, 1H), 7.40 (d, *J* = 7.7 Hz, 2H), 7.28 (d, *J* = 7.7 Hz, 2H), 6.13 (s, 1H), 5.29 (s, 2H), 4.30 (d, *J* = 5.8 Hz, 2H), 3.88 (s, 2H).

**^13^C NMR** (101 MHz, DMSO-*d*_6_) *δ* [ppm] = 170.28, 167.32, 162.77, 160.01, 138.90, 134.90, 128.44, 127.46, 94.42, 67.21, 50.78, 42.01.

**HRMS** (ESI) calc. for C_14_H_15_ClN_7_O_2_^+^ [M+H]^+^: 348.0970; found 348.0971.

##### 1.5.2 *N*-(4-(((2-amino-6-chloropyrimidin-4-yl)oxy)methyl)benzyl)-2-((1*S*,4*S*)-bicyclo[2.2.1]hept-5-en-2-yl)acetamide (CP-Nor2)

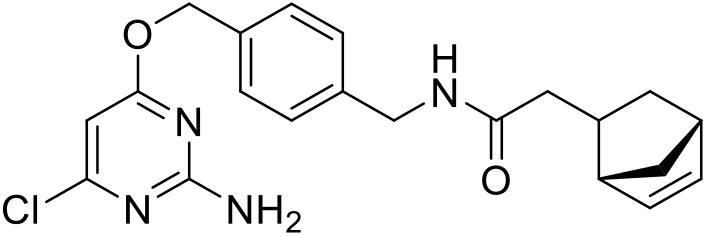

Reaction was conducted according to general procedure A with a reduced reaction time of 15 min. using CP-NH_2_ and 2-((1*S*,4*S*)-bicyclo[2.2.1]hept-5-en-2-yl)acetic acid (7.1 μL, 41.6 μmol) resulting in 14.5 mg (28.3 μmol) of the desired product as a colorless TFA-salt in 75% yield.

**^1^H NMR** (400 MHz, DMSO-*d*_6_) *δ* [ppm] = 8.25 (t, *J* = 5.9 Hz, 1H), 7.38 (d, *J* = 8.1 Hz, 2H), 7.23 (d, *J* = 8.1 Hz, 2H), 7.10 (brs, 2H), 6.16 (dd, *J* = 5.7, 3.0 Hz, 1H), 6.13 (s, 1H), 5.96 (dd, *J* = 5.7, 2.9 Hz, 1H), 5.28 (s, 2H), 4.24 (d, *J* = 5.9 Hz, 2H), 2.77 – 2.69 (m, 2H), 2.46 – 2.35 (m, 1H), 1.94 (dd, *J* = 13.8, 7.6 Hz, 1H), 1.90 – 1.76 (m, 2H), 1.35 – 1.26 (m, 1H), 1.26 – 1.18 (m, 1H), 0.50 (m, *J* = 11.4, 4.3, 2.5 Hz, 1H).

**^13^C NMR** (101 MHz, DMSO-*d*_6_) *δ* [ppm] = 171.61, 170.28, 162.75, 159.97, 139.81, 137.01, 134.53, 132.42, 128.31, 127.11, 94.39, 67.23, 49.07, 45.24, 42.06, 41.69, 40.64, 35.11, 31.45.

**HRMS** (ESI) calc. for C_21_H_23_ClN_4_NaO_2_^+^ [M+Na]^+^; 421.1402; found 421.1403.

##### 1.5.3 *N*-(4-(((2-amino-6-chloropyrimidin-4-yl)oxy)methyl)benzyl)-2-(4-(6-methyl-1,2,4,5-tetrazin-3-yl)phenyl)acetamide (CP-Tz)

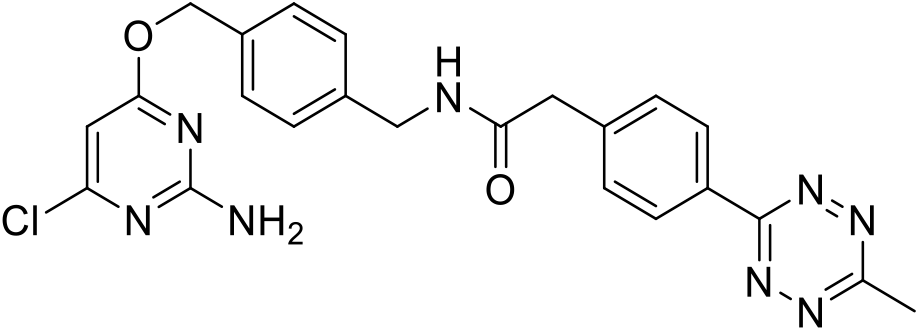

Reaction was conducted according to general procedure A using CP-NH_2_ and 2-(4-(6-methyl-1,2,4,5-tetrazin-3-yl)phenyl)acetic acid (9.6 mg, 41.6 μmol). The product was purified by preparative HPLC eluted with MeCN / H_2_O (0.1% TFA) (10% MeCN for 10 min., then 10 - 90% MeCN over 40 min., followed by 90% MeCN for 10 min.) to give 2.6 mg (4.4 μmol) of the desired product as a rose TFA-salt in 12% yield after lyophilization.

**^1^H NMR** (400 MHz, DMSO-*d*_6_) *δ* [ppm] = 8.66 (t, *J* = 5.9 Hz, 1H), 8.41 (d, *J* = 8.3 Hz, 2H), 7.56 (d, *J* = 8.3 Hz, 2H), 7.38 (d, *J* = 7.9 Hz, 2H), 7.26 (d, *J* = 7.9 Hz, 2H), 7.10 (s, 2H), 6.13 (s, 1H), 5.28 (s, 2H), 4.29 (d, *J* = 5.9 Hz, 2H), 3.63 (s, 2H), 2.99 (s, 3H).

**^13^C NMR** (101 MHz, DMSO-*d*_6_) *δ* [ppm] = 170.28, 169.51, 167.04, 163.21, 162.76, 159.99, 141.13, 139.32, 134.75, 130.14, 130.07, 128.42, 127.34, 94.40, 67.22, 42.21, 42.09, 20.83.

**HRMS** (ESI) calc. for C_23_H_22_ClN_8_O_2_^+^ [M+H]^+^: 477.1549; found 477.1553.

##### 1.5.4 *N*-(4-(((2-amino-6-chloropyrimidin-4-yl)oxy)methyl)benzyl)-4-azidobenzamide (CP-PhN_3_)

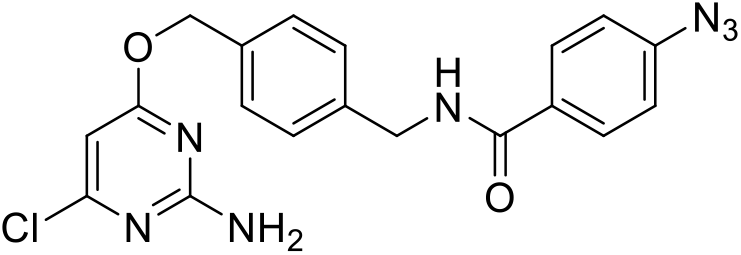

Reaction was conducted according to general procedure A with a reduced reaction time of 15 min. using BG-NH_2_ and 4-azidobenzoic acid (6.8 mg, 41.6 μmol) to obtain 12.0 mg (22.9 μmol) of the desired product as a colorless TFA-salt in 61% yield.

**^1^H NMR** (400 MHz, DMSO-*d*_6_) *δ* [ppm] = 9.06 (t, *J* = 5.9 Hz, 1H), 7.94 (d, J = 8.4 Hz, 2H), 7.39 (d, *J* = 7.9 Hz, 2H), 7.32 (d, *J* = 7.9 Hz, 2H), 7.21 (d, *J* = 8.4 Hz, 2H), 7.10 (s, 2H), 5.29 (s, 2H), 4.47 (d, *J* = 5.9 Hz, 2H).

**^13^C NMR** (101 MHz, DMSO-*d*_6_) *δ* [ppm] = 170.27, 165.20, 162.75, 159.97, 142.30, 139.59, 134.66, 130.83, 129.12, 128.36, 127.29, 118.90, 94.38, 67.23, 42.44.

**HRMS** (ESI) calc. for C_19_H_16_ClN_7_NaO_2_^+^ [M+Na]^+^: 432.0946; found 432.0942.

##### 1.5.5 *N*-(4-(((2-amino-6-chloropyrimidin-4-yl)oxy)methyl)benzyl)-4-vinylbenzamide (CP-Vbn)

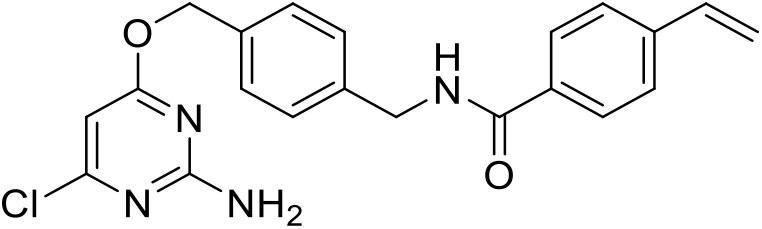

Reaction was conducted according to general procedure A with a reduced reaction time of 15 min. using CP-NH2 and 4-vinylbenzoic acid (41.6 μmol; 6.2 mg) to obtain 11.6 mg (22.8 μmol) of the desired product as a colorless TFA-salt in 60% yield.

**^1^H NMR** (400 MHz, DMSO-*d*_6_) *δ* [ppm] = 9.04 (t, *J* = 6.0 Hz, 1H), 7.87 (d, *J* = 8.3 Hz, 2H), 7.57 (d, *J* = 8.3 Hz, 2H), 7.40 (d, *J* = 8.1 Hz, 2H), 7.34 (d, *J* = 8.1 Hz, 2H), 6.79 (dd, *J* = 17.7, 10.9 Hz, 1H), 7.10 (brs, 2H), 6.12 (s, 1H), 5.95 (d, *J* = 17.7 Hz, 1H), 5.37 (d, *J* = 10.9 Hz, 1H), 5.29 (s, 2H), 4.47 (d, *J* = 6.0 Hz, 2H).

**^13^C NMR** (101 MHz, DMSO-*d*_6_) *δ* [ppm] = 170.29, 165.77, 162.77, 159.99, 139.80, 139.68, 135.91, 134.66, 133.45, 128.41, 127.61, 127.31, 126.00, 116.20, 94.40, 67.27, 42.43.

**HRMS** (ESI) calc. for C_21_H_20_ClN_4_O_2_^+^ [M+H]^+^: 395.1269; found 395.1258.

##### 1.5.6 4-(Benzyloxy)-6-chloropyrimidin-2-amine (CP)

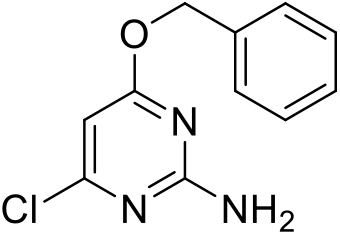

2-Amino-4,6-dichloropyrimidine (200 mg, 1.22 mmol, 1.0 equiv.) was dissolved in dry DMF (2 mL). Benzyl alcohol (63 μL, 1.22 mmol, 1.0 equiv.), KO^*t*^Bu (342.2 mg, 3.04 mmol, 2.5 equiv.) and KI (20.2 mg, 0.122 mmol, 0.1 equiv.) were added and the reaction mixture was stirred at room temperature for 4 h. Afterwards, the reaction was quenched with water and extracted with EtOAc (3 ×). The combined organic layers were washed with brine and dried over MgSO_4_. The volatiles were evaporated and the crude product was purified over normal phase flash chromatography (hexane:DCM = 50%: 50% to 100% DCM). The fractions containing the product were combined, volatiles were evaporated and 134 mg (0.569 mmol) of the desired product was obtained as a yellowish solid in 47% yield.

**^1^H NMR** (400 MHz, CDCl3): *δ* = 7.43–7.30 (m, 5H), 6.01 (d, J = 0.7 Hz, 1H), 5.31 (s, 2H), 2.26 (s, 3H) ppm.

**^13^C NMR** (101 MHz, CDCl3): *δ* = 170.6, 168.4, 162.6, 136.7, 128.7, 128.5, 128.0, 127.4, 97.2, 93.0, 77.4, 77.1, 76.7, 67.5, 123.7 ppm.

**HRMS** (ESI) calc. for C_11_H_11_ClN_3_O^+^ [M+H]^+^: 236.0585; found 236.0583.

##### 1.5.7 ((1*R*,8*S*,9*s*)-bicyclo[6.1.0]non-4-yn-9-yl)methyl(4-(((2-amino-6-chloropyrimidin-4-yl)oxy)methyl)benzyl)carbamate (CP-BCN)

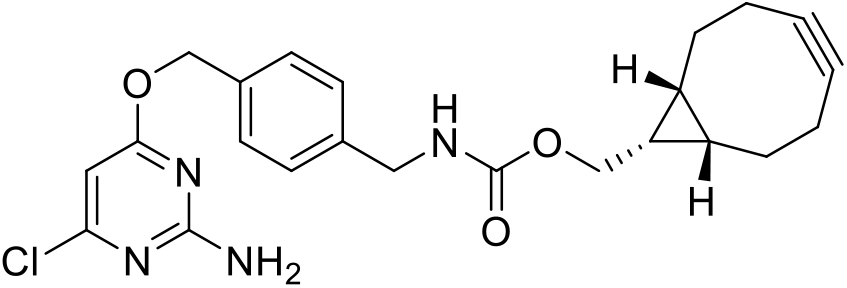

((1*R*,8*S*,9*s*)-bicyclo[6.1.0]non-4-yn-9-yl)methyl (2,5-dioxopyrrolidin-1-yl) carbonate (10.0 mg, 34.3 μmol; 1.0 equiv.) was dissolved in dry DMSO (0.5 mL) and DIPEA (28.4 μL, 172 μmol, 5 equiv.) followed by CP-NH_2_ (10.0 mg, 37.8 μmol, 1.1 equiv.) were added. The reaction was stirred at r.t. for 30 min. The resulted mixture was acidified with acetic acid (3 μL) and H2O (53 μL), then purified by preparative HPLC eluted with MeCN / H_2_O (0.1% TFA) (10% MeCN for 10 min., then 10 - 90% MeCN over 55 min., followed by 99% MeCN for 5 min.) to give 1.4 mg (3.11 μmol) of the desired product as a colorless solid in 9% yield after lyophilization.

**^1^H NMR** (400 MHz, DMSO-*d*_6_) *δ* [ppm] = 7.68 (q, *J* = 6.4 Hz, 1H), 7.38 (d, *J* = 8.0 Hz, 2H), 7.25 (d, *J* = 7.8 Hz, 2H), 6.12 (s, 1H), 5.28 (s, 2H), 4.17 (d, *J* = 6.1 Hz, 2H), 4.06 (d, *J* = 8.0 Hz, 2H), 2.29 – 1.72 (m, 6H), 1.71 – 1.38 (m, 2H), 1.35 – 0.60 (m, 3H).

**HRMS** (ESI) calc. for C_23_H_26_ClN_4_O_3_^+^ [M+H]^+^: 441.1688; found 441.1688.

##### 1.5.8 *N*-(4-(((2-amino-6-chloropyrimidin-4-yl)oxy)methyl)benzyl)acetamide (CP-Ac)

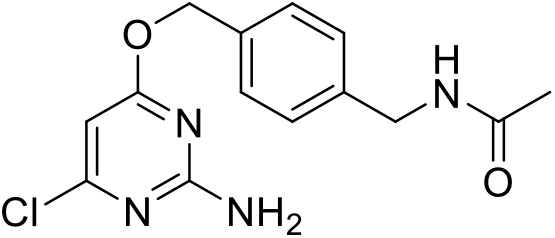

CP-NH_2_ (300 mg, 1.13 mmol, 1.0 equiv.) was dissolved in dry DMSO (1.5 mL) and DIPEA (375 μL, 2.27 mmol, 2.0 equiv.) was added followed by dropwise addition of acetic anhydride (160 μL, 1.70 mmol, 1.5 equiv.) while stirring. The reaction mixture was stirred at r.t. for 1 h. Afterwards, the reaction was quenched with acetic acid (387 μL) and H_2_O (341 μL) followed by purification over preparative HPLC eluted with MeCN / H_2_O (0.1% TFA) (30% MeCN for 10 min., then 30 - 90% MeCN over 55 min., followed by 99% MeCN for 5 min.) to give 201 mg (655 μmol) of the desired product as a colorless solid in 58% yield after lyophilization.

**^1^H NMR** (400 MHz, DMSO-*d*_6_) *δ* [ppm] = 8.33 (t, *J* = 6.0 Hz, 1H), 7.41 – 7.35 (m, 2H), 7.28 – 7.22 (m, 2H), 7.09 (s, 2H), 6.13 (s, 1H), 5.29 (s, 2H), 4.24 (d, *J* = 5.9 Hz, 2H), 1.86 (s, 3H).

**HRMS** (ESI) calc. for C_14_H_16_ClN_4_O_2_^+^ [M+H]^+^: 307.0956; found 307.0957.

##### 1.5.9 Cyclooct-2-yn-1-yl (4-(((2-amino-6-chloropyrimidin-4-yl)oxy)methyl)benzyl)carbamate (CP-SCO)

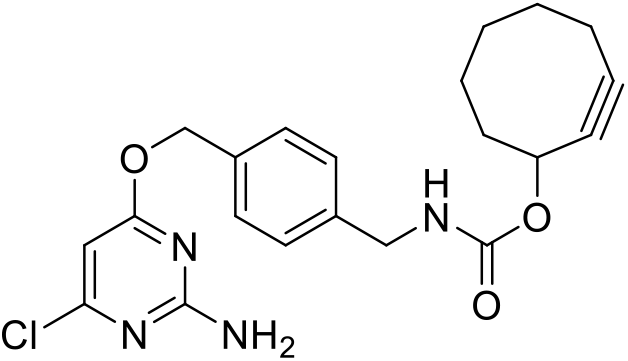

CP-NH_2_ (10 mg; 37.8 μmol, 1.3 equiv.) was dissolved in dry DMF (0.3 mL) and a solution of 8.4 mg cyclooct-2-yn-1-yl (4-nitrophenyl) carbonate (29.1 μmol, 1.0 equiv.) in dry DMF (0.2 mL) was added followed by DIPEA (24.0 μL, 145 μmol: 5.0 equiv.). The reaction mixture was stirred at r.t. for 2 h. The resulted mixture was acidified with acetic acid (25 μL) and afterwards purified by semi-preparative HPLC eluted with MeCN / H_2_O (0.1% TFA) (15% MeCN for 2 min., then 15 - 100% MeCN over 25 min., followed by 100% MeCN for 15 min.) to give 12.0 mg (22.7 μmol) of the desired product as a colorless TFA-salt in 78% yield after lyophilization.

**^1^H NMR** (400 MHz, DMSO-*d*_6_) *δ* [ppm] = 7.78 (t, *J* = 6.2 Hz, 1H), 7.38 (d, *J* = 8.1 Hz, 2H), 7.24 (d, *J* = 8.1 Hz, 2H), 6.13 (s, 1H), 5.28 (s, 2H), 5.21 – 5.11 (m, 1H), 4.15 (d, *J* = 6.2 Hz, 2H), 2.29 – 2.02 (m, 3H), 1.95 – 1.85 (m, 1H), 1.85 – 1.78 (m, 2H), 1.76 – 1.65 (m, 1H), 1.65 – 1.54 (m, 2H), 1.54 – 1.42 (m, 1H).

**^13^C NMR** (101 MHz, DMSO-*d*_6_) *δ* [ppm] = 170.30, 162.78, 159.99, 155.51, 139.64, 134.73, 128.40, 127.11, 100.94, 94.41, 91.76, 67.25, 65.93, 43.50, 41.58, 33.84, 29.21, 25.79, 19.96.

**HRMS** (ESI) calc. for C_21_H_23_ClN_4_NaO_3_^+^ [M+Na]^+^: 437.1351; found 437.1358.

##### 1.5.10 2-(6-amino-3-iminio-4,5-disulfonato-3*H*-xanthen-9-yl)-4-((4-(((2-amino-6-chloropyrimidin-4-yl)oxy)methyl)benzyl)carbamoyl)benzoate (CP-Alexa488)

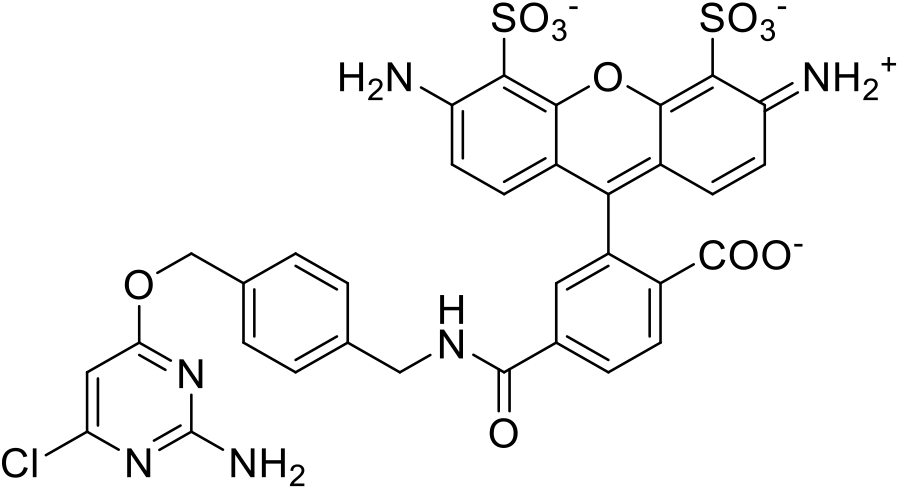

In an Eppendorf tube, CP-NH_2_ (0.34 μg, 1.27 μmol, 2.0 equiv.) was dissolved in dry DMSO (100 μL) followed by addition of DIPEA (885 μL, 5.1 μmol, 8.0 equiv.) and a solution of 2-(6-amino-3-iminio-4,5-disulfonato-3H-xanthen-9-yl)-4-(((2,5-dioxopyrrolidin-1-yl)oxy)carbonyl)benzoate (0.4 mg, 0.64 μmol, 1.0 equiv.) in dry DMSO (100 μL). The reaction was kept at r.t. for 1 h. The compound was purified over preparative HPLC eluted with MeCN / H_2_O (0.1% TFA) (10% MeCN for 10 min., then 10 - 90% MeCN over 40 min., followed by 90% MeCN for 5 min.) to give 195 μg (252 nmol) of the desired product as a yellow solid in 79% yield after lyophilization.

**HRMS** (ESI) calc. for C_33_H_25_ClN_6_O_11_S_2_ [M+3H]^+^: 781.0784; found 781.0772.

##### 1.5.11 4-((4-(((2-amino-6-chloropyrimidin-4-yl)oxy)methyl)benzyl)carbamoyl)-2-(6-hydroxy-3-oxo-3H-xanthen-9-yl)benzoate (CP-Fluorescein)

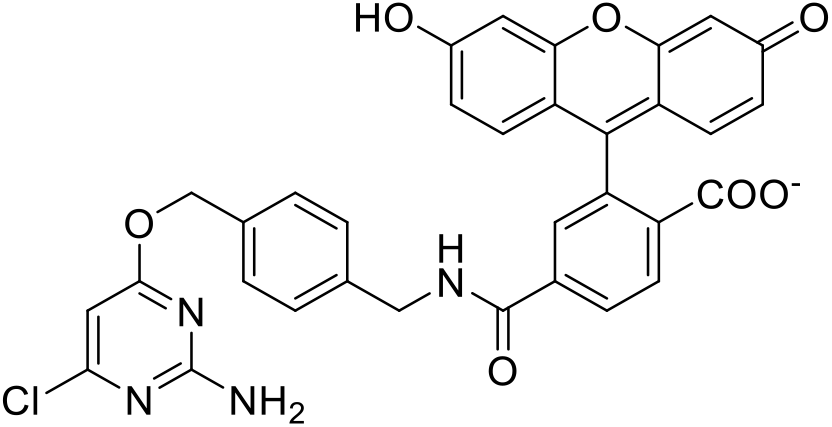

Fluorescein-6-COOH (25.0 mg, 66.4 μmol, 1.0 equiv.) was dissolved in dry DMSO (1.25 mL) and DIPEA (22.0 μL, 133 μmol, 2.0 equiv.) as well as TSTU (24.0 mg, 79.7 μmol, 1.2 equiv.) were added and the mixture was stirred at r.t. for 30 min. Afterwards, CP-NH2 (26.4 mg, 99.7 μmol, 1.5 equiv.) was added and the reaction mixture was stirred at r.t. for 1 h. The resulted mixture was quenched with acetic acid (22.0 μL) and 10% H_2_O, then the compound was purified over preparative HPLC eluted with MeCN / H_2_O (0.1% TFA) (10% MeCN for 10 min., then 10 - 90% MeCN over 40 min., followed by 90% MeCN for 5 min.) to give 31 mg (49.8 μmol) of the desired product in 75% yield after lyophilization.

**HRMS** (ESI) calc. for C_33_H_24_ClN_4_O_7_^+^ [M+H]^+^: 623.1328; found 623.1327.

##### 1.5.12 4-((4-(((2-amino-6-chloropyrimidin-4-yl)oxy)methyl)benzyl)carbamoyl)-2-(6-(dimethylamino)-3-(dimethyliminio)-10,10-dimethyl-3,10-dihydroanthracen-9-yl)benzoate (CP-CPY)

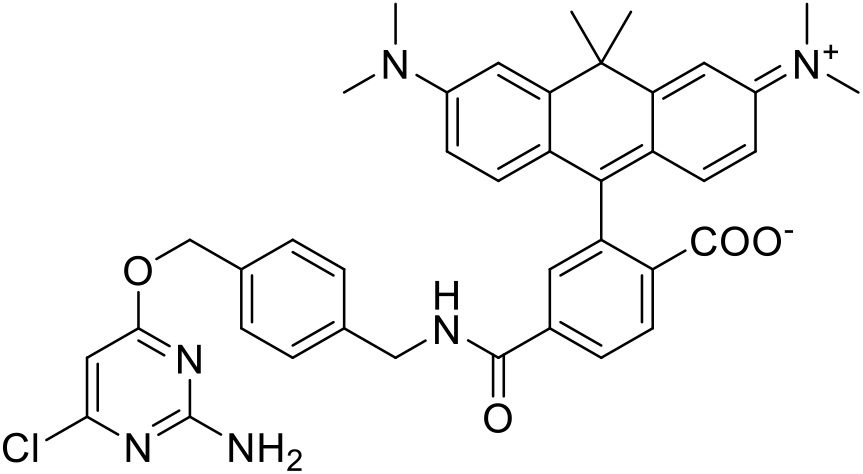

CPY-6-COOH(1) (250 mg, 530 μmol, 1.0 equiv.) was dissolved in dry DMSO (2 mL) and DIPEA (362 μL, 2.19 mmol, 4.0 equiv.) as well as TSTU (231 mg, 767 μmol, 1.4 equiv.) were added and the mixture was stirred at r.t. for 5 min. Afterwards, CP-NH_2_ (217 mg, 821 μmol, 1.5 equiv.) was added and the reaction mixture was stirred at r.t. for 35 min. The resulted mixture was acidified with acetic acid (362 μL) and H_2_O (500 μL), then the compound was purified over preparative HPLC eluted with MeCN / H_2_O (0.1% TFA) (10% MeCN for 10 min., then 10 - 90% MeCN over 40 min., followed by 90% MeCN for 5 min.) to give 130 mg (184.9 μmol) of the desired product in 34% yield after lyophilization.

**^1^H NMR** (400 MHz, acetone-*d*_6_) *δ* [ppm] = 8.51 (t, J = 6.4 Hz, 1H), 8.23 (d, J = 8.1 Hz, 1H), 8.12 (d, J = 8.7 Hz, 1H), 7.67 (s, 1H), 7.39 – 7.30 (m, 4H), 7.11 (s, 2H), 6.67 (s, 4H), 6.36 (s, 1H), 6.07 (m, J = 10.7, 2.5 Hz, 1H), 5.30 (m, J = 11.2, 2.5 Hz, 2H), 4.55 (d, J = 5.9 Hz, 2H), 3.11 (s, 12H), 1.89 (d, J = 2.5 Hz, 3H), 1.76 (d, J = 2.4 Hz, 3H).

**^13^C NMR** (101 MHz, acetone-*d*_6_) *δ* [ppm] = 171.72, 165.87, 161.60, 140.12, 136.37, 134.01, 129.34, 129.25, 128.85, 120.23, 113.03, 110.69, 96.16, 68.31, 44.02, 40.62, 35.59, 33.04, 30.42, 30.22, 30.03, 29.84, 29.65, 29.45, 29.26, 26.13.

**HRMS** (ESI) calc. for C_40_H_39_ClN_6_O_4_^+^ [M+H]^+^: 703.2794; found 703.2792.

##### 1.5.13 4-((4-(((2-amino-6-chloropyrimidin-4-yl)oxy)methyl)benzyl)carbamoyl)-2-(7-(dimethylamino)-3-(dimethyliminio)-5,5-dimethyl-3,5-dihydrodibenzo[b,e]silin-10-yl)benzoate (CP-SiR)

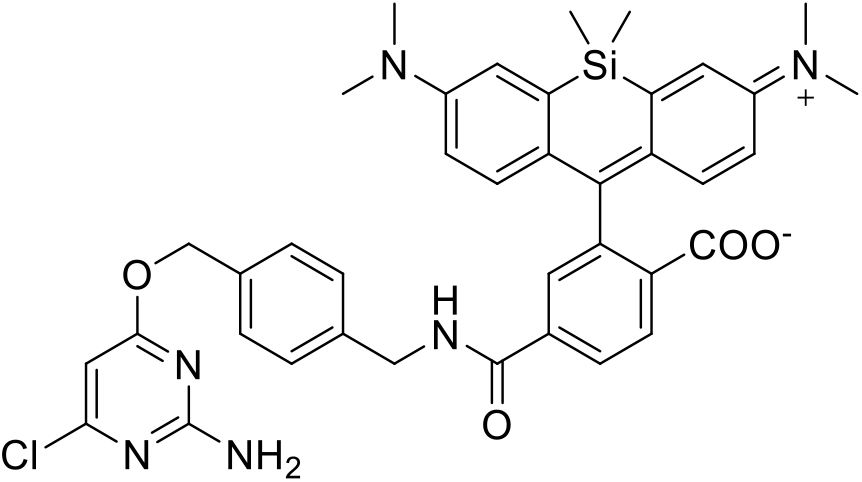

SiR-6-COOH(4) (481 mg, 1.02 mmol, 1.1 equiv.) was dissolved in dry DMSO (4 mL) and DIPEA (919 μL, 5.56 mmol, 6.0 equiv.) was added. The mixture was sonicated until complete solution and TSTU (391 mg, 1.30 mmol, 1.4 equiv.) were added and the mixture was stirred at r.t. for 5 min. Afterwards, CP-NH_2_ (294 mg, 1.11 mmol, 1.2 equiv.) was added and the reaction mixture was stirred at r.t. for 2h. The resulted mixture was quenched by addition of acetic acid (973 μL) and 10% H_2_O, followed by purification over preparative HPLC eluted with MeCN / H_2_O (0.1% FA) (10% - 90% MeCN over 60 min) to give. 355 mg (494 μmol) of the desired product in 53% yield after lyophilization.

**HRMS** (ESI): calc. for C_39_H_39_N_6_O_4_Si^+^ [M+H]^+^: 719.2563; found 719.2561.

#### 1.6 CLIP substrates

##### 1.6.1 4-((4-(((4-aminopyrimidin-2-yl)oxy)methyl)benzyl)carbamoyl)-2-(6-(dimethylamino)-3-(dimethyliminio)-10,10-dimethyl-3,10-dihydroanthracen-9-yl)benzoate (BC-CPY)

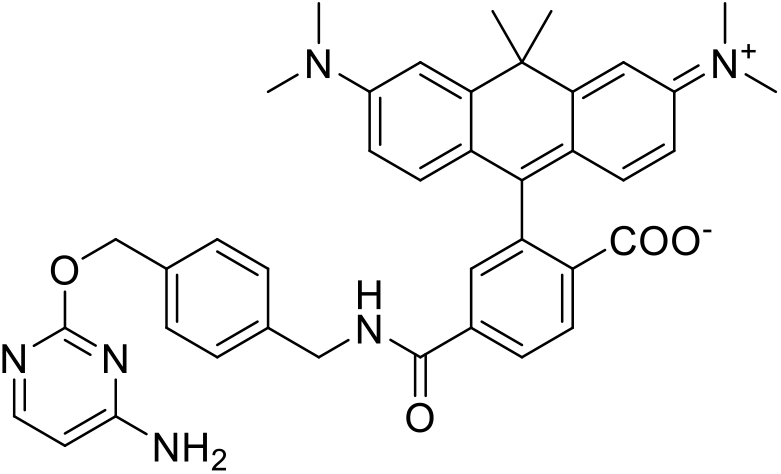

CPY-6-COOH(1) (250 mg, 530 μmol, 1.0 equiv.) was dissolved in dry DMSO (2 mL). DIPEA (362 μL, 2.19 mmol, 4.0 equiv.) and TSTU (231 mg, 767 μmol, 1.4 equiv.) were added and the mixture was stirred at r.t. for 5 min. Afterwards, BC-NH_2_ (189 mg, 821 μmol, 1.5 equiv.) was added and the reaction mixture was stirred at r.t. for 35 min. The resulted mixture was acidified with acetic acid (362 μL) and H_2_O (500 μL), then compound was purified over preparative HPLC eluted with MeCN / H_2_O (0.1% TFA) (10% MeCN for 10 min., then 10 - 90% MeCN over 40 min., followed by 90% MeCN for 5 min.) to give 180 mg (269.1 μmol) of the desired product in 49% yield after lyophilization.

**^1^H NMR** (400 MHz, acetone-*d*_6_) *δ* [ppm] = 8.51 (t, *J* = 6.9 Hz, 1H), 8.21 (d, *J* = 8.1 Hz, 1H), 8.10 – 8.01 (m, 2H), 7.61 (s, 1H), 7.38 (d, *J* = 7.4 Hz, 2H), 7.32 (d, *J* = 7.8 Hz, 2H), 7.27 (s, 1H), 7.05 (s, 2H), 6.61 (s, 4H), 6.40 (d, *J* = 6.6, 2.4 Hz, 1H), 5.36 (s, 2H), 4.56 – 4.50 (m, 2H), 3.04 (s, 12H), 1.88 (d, *J* = 2.5 Hz, 3H), 1.76 (d, *J* = 2.6 Hz, 3H).

**^13^C NMR** (101 MHz, acetone-*d*_6_) *δ* [ppm] = 169.57, 165.97, 162.80, 152.54, 152.01, 148.78, 141.22, 140.20, 136.01, 130.47, 129.26, 128.83, 126.23, 124.12, 120.13, 112.83, 110.38, 100.42, 69.65, 44.02, 43.89, 40.54, 39.65, 35.58, 33.19, 30.42, 30.23, 30.03, 29.84, 29.65, 29.46, 29.26.

**HRMS** (ESI): calc. for C_40_H_42_N_6_O_4_^2+^ [M+2H] ^2+^: 335.1628; found 335.1629.

#### 1.7 Additional substrates

##### 1.7.1 2-(6-(dimethylamino)-3-(dimethyliminio)-3H-xanthen-9-yl)-4-(methylcarbamoyl)benzoate (meAm-6-TMR)

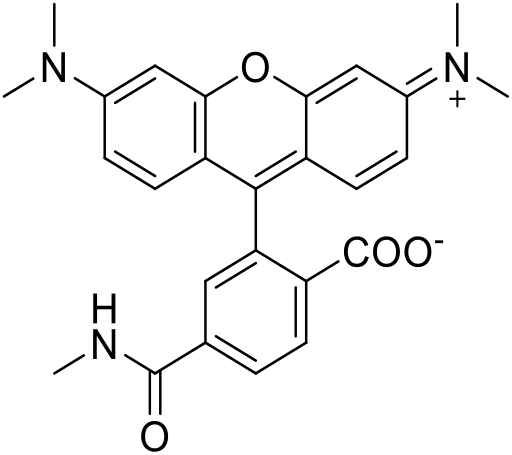

To a solution of TMR-6-COOH (1.0 mg, 2.32 μmol, 1.1 equiv.) in dry DMSO (500 μL), TSTU (763 μg, 2.53 μmol, 1.2 equiv.) was added and the mixture was stirred at r.t. for 5 min. Afterwards, DIPEA (1.4 μL, 8.45 μmol, 4 equiv.) and methylamine (2 M, 1.06 μL, 2.11 μmol, 1 equiv.) were added and the reaction mixture was stirred at r.t. overnight. The compound was purified over preparative HPLC eluted with MeCN / H_2_O (0.1% TFA) (10% MeCN for 10 min., then 10 - 90% MeCN over 40 min., followed by 90% MeCN for 5 min.) to give 91.1 μg (205.4 nmol) of the desired product in 10% yield after lyophilization.

**^1^H NMR** (TMR-6-COOH) (400 MHz, DMSO-*d*_6_) *δ* [ppm] = 8.21 (dd, *J* = 8.0, 1.4 Hz, 1H), 8.17 – 7.99 (m, 1H), 7.61 – 7.56 (m, 1H), 6.58 – 6.45 (m, 6H), 2.95 (s, 12H).

**^13^C NMR** (TMR-6-COOH) (101 MHz, DMSO-*d*_6_) *δ* [ppm] = 168.56, 166.53, 152.67, 152.47, 131.16, 128.91, 109.56, 105.91, 98.43, 40.46, 40.26.

**HRMS** (ESI): calc. for C_26_H_26_N_3_O_4_^+^ [M+H]^+^: 444.1918; found 444.1914.

##### 1.7.2 2-(6-(dimethylamino)-3-(dimethyliminio)-10,10-dimethyl-3,10-dihydroanthracen-9-yl)-4-(methylcarbamoyl)benzoate (meAm-6-CPY)

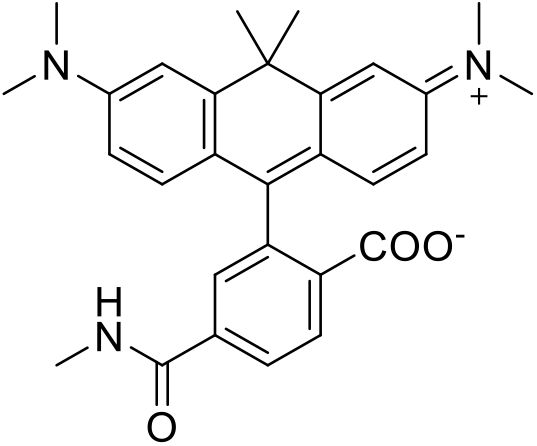

To a solution of CPY-6-COOH (1.0 mg, 2.19 μmol, 1.1 equiv.) in dry DMSO (500 μL), TSTU (719 μg, 2.39 μmol, 1.2 equiv.) was added and the mixture was stirred at r.t. for 5 min. Afterwards, DIPEA (1.32 μL, 7.97 μmol, 4.0 equiv.) and methylamine (2 M, 0.996 μL, 1.99 μmol, 1.0 equiv.) were added and the reaction mixture was stirred at r.t. overnight. The compound was purified over preparative HPLC eluted with MeCN / H_2_O (0.1% TFA) (10% MeCN for 10 min., then 10 - 90% MeCN over 40 min., followed by 90% MeCN for 5 min.) to give 97.7 μg (208.1 nmol) of the desired product in 10% yield after lyophilization.

**HRMS** (ESI): calc. for C_29_H_32_N_3_O_3_ [M+H]^+^: 470.2438; found 470.2434.

##### 1.7.3 2-(6-(dimethylamino)-3-(dimethyliminio)-3H-xanthen-9-yl)-5-(methylcarbamoyl)benzoate (meAm-5-TMR)

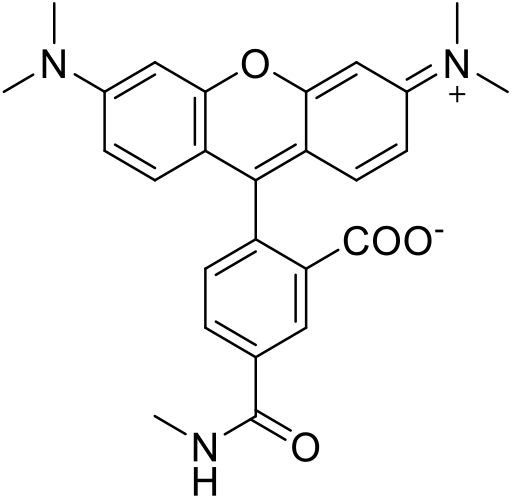

To a solution of TMR-5-COOH (2.5 mg, 5.81 μmol, 1.0 equiv.) in dry DMSO (500 μL), BOP (2.59 mg, 8.71 μmol, 1.5 equiv.) was added and the reaction was shaken at r.t. and 500 rpm for 5 min. DIPEA (3.84 μL, 23.2 μmol, 4.0 equiv.) and methylamine (2M in THF, 4.36 μL, 8.71 μmol, 1.5 equiv.) were added and the reaction was shaken at r.t. and 500 rpm for 4 h. The crude product was acidified with acetic acid and purified over preparative HPLC eluted with MeCN / H_2_O (0.1% FA) (10% - 90% MeCN over 50 min) to give 0.97 mg (2.19 μmol) of the desired product in 38% yield after lyophilization.

**^1^H NMR** (TMR-5-COOH) (400 MHz, DMSO-*d*_6_) *δ* [ppm] = 8.39 (s, *J* = 1.5 Hz, 1H), 8.28 (dd, *J* = 8.1, 1.5 Hz, 1H), 7.33 (d, *J* = 8.0 Hz, 1H), 6.58 – 6.45 (m, 6H), 2.95 (s, 12H).

**^13^C NMR** (TMR-5-COOH) (101 MHz, DMSO-*d*_6_) *δ* [ppm] = 168.31, 166.09, 152.03, 135.96, 132.76, 128.50, 109.05, 97.95, 40.15, 39.99, 39.79.

**HRMS** (ESI): calc. for C_26_H_26_N_3_O_4_^+^ [M+H]^+^: 444.1923; found 444.1914.

##### 1.7.4 2-(6-(dimethylamino)-3-(dimethyliminio)-10,10-dimethyl-3,10-dihydroanthracen-9-yl)-5-(methylcarbamoyl)benzoate (meAm-5-CPY)

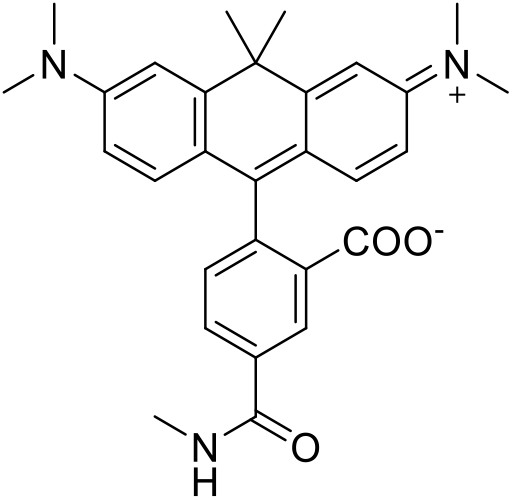

To a solution of CPY-5-COOH (2.5 mg, 5.48 μmol, 1.0 equiv.) in dry DMSO (1 mL), BOP (0.5 M in DMSO, 17.4 μL, 8.71 μmol, 1.5 equiv.) was added and the reaction was shaken at r.t and 500 rpm for 5 min. DIPEA (3.62 μL, 21.9 μmol, 4.0 equiv.) and methylamine (2 M in THF, 4.11 μL, 8.21 μmol, 1.5 equiv.) were added and the reaction was shaken at 500 rpm, r.t. for 4 h. The crude product was acidified with acetic acid and purified over preparative HPLC eluted with MeCN / H_2_O (0.1% FA) (10% - 90% MeCN over 50 min) to give 0.77 mg (1.64 μmol) of the desired product in 30% yield after lyophilization.

**HRMS** (ESI): calc. for C_29_H_32_N_3_O_3_^+^ [M+H]^+^: 470.2443; found 470.2437.

**Figure S1:**
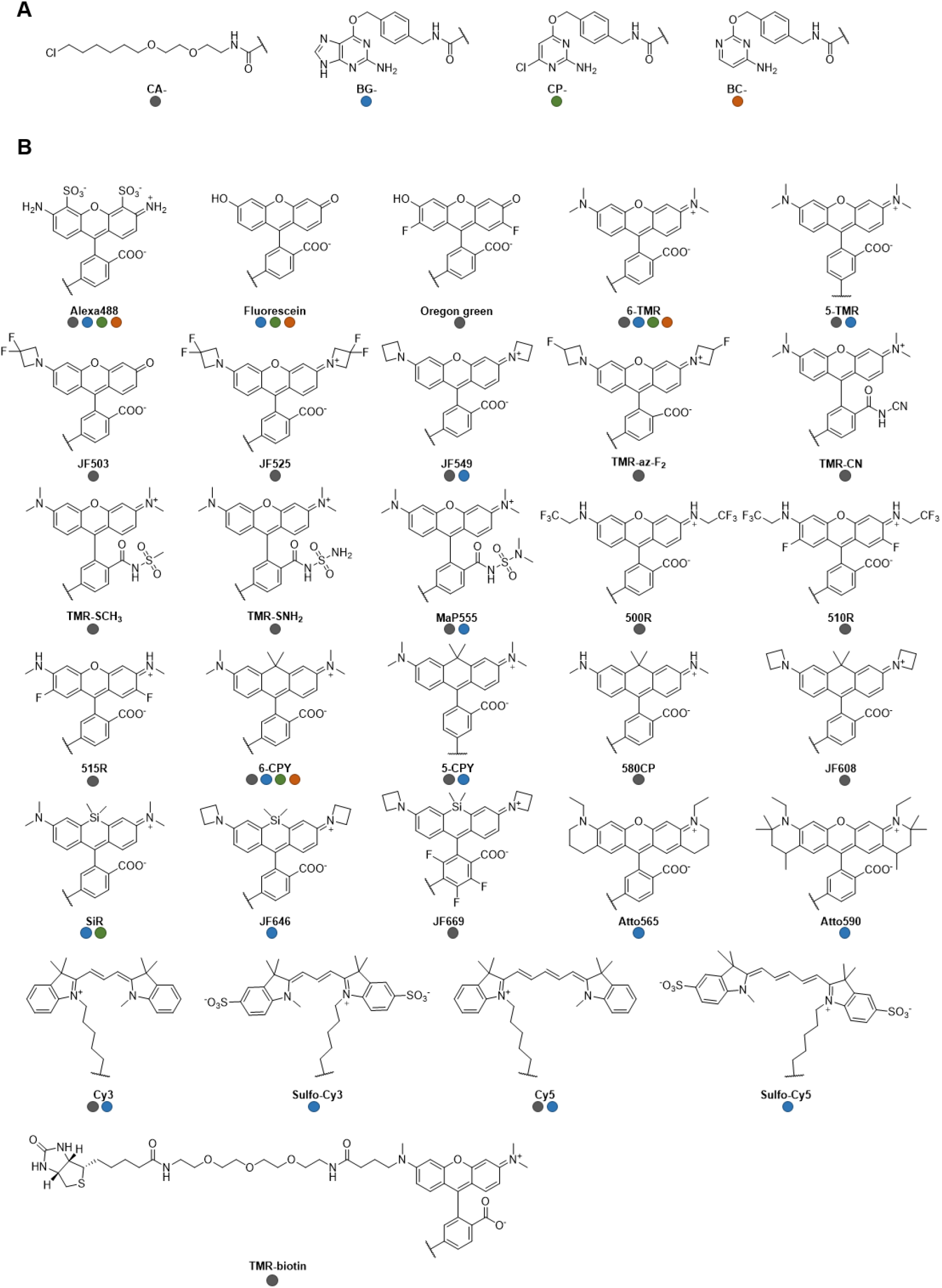
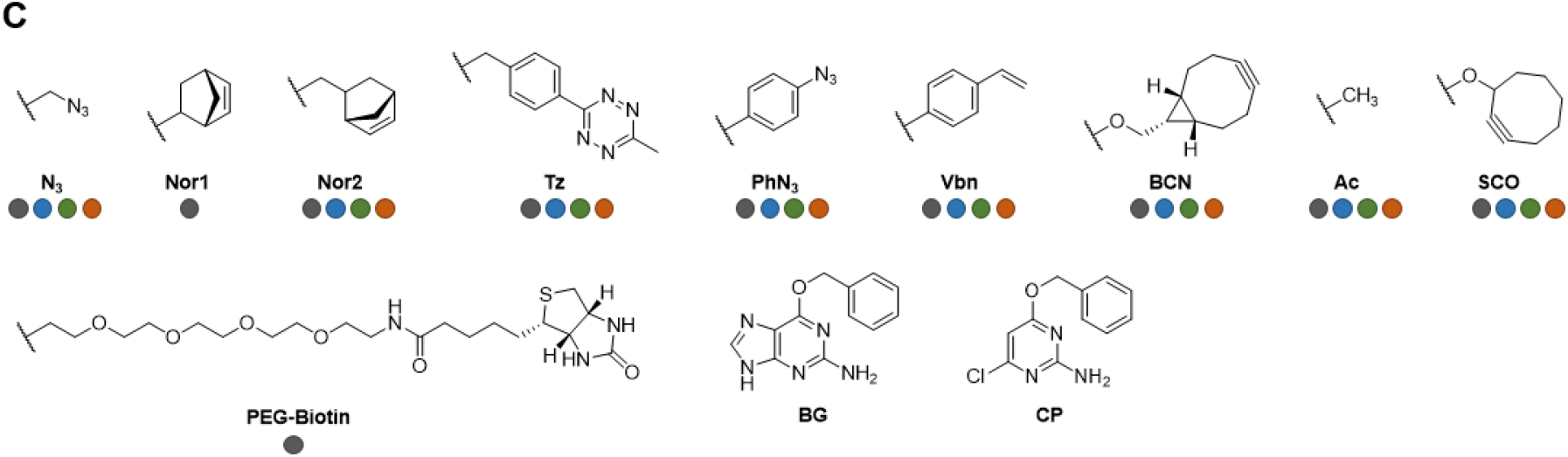
Chemical structures of SLP substrates. **A**. Chemical structures of HT7 (CA), SNAP (BG and CP) and CLIP (BC) core substrates. **B**. Chemical structures of fluorescent substituents. **C**. Chemical structures of non-fluorescent substituents. Colored dots indicate the tested substrates for the corresponding SLPs (grey = CA, blue = BG, green = CP and orange = BC).

**Figure S2:**
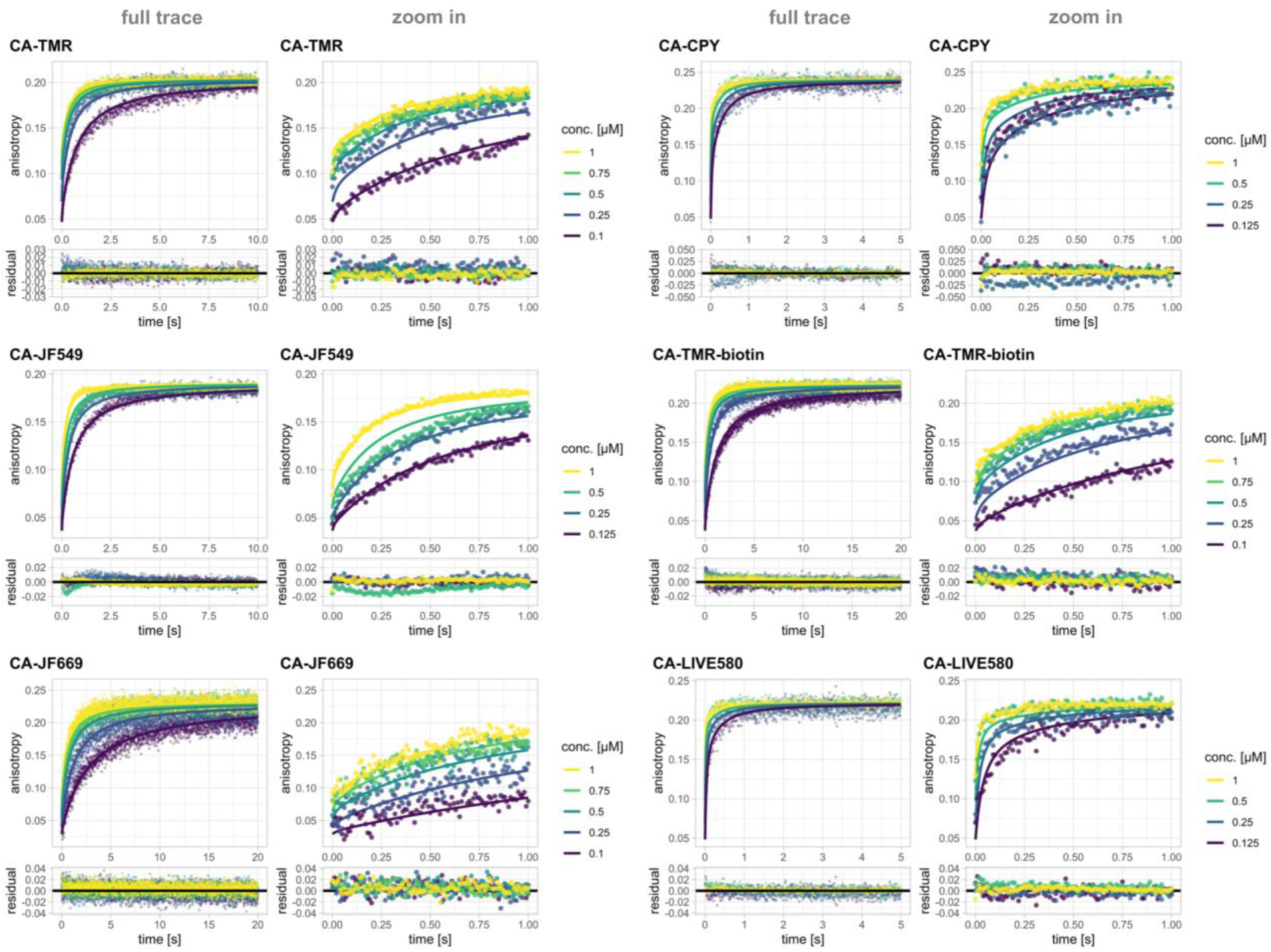
Labeling kinetics of HT7 with fluorescent CA substrates. Full anisotropy traces (points) and predications of fits based on model 2 (lines) along with zoom on the first second are represented on the top panels. Residuals from the fits are depicted in the bottom panels. Kinetics were recorded by following fluorescence anisotropy changes over time using a stopped flow device. All conditions are 1:1 mixtures of protein and substrate at the given concentrations (conc.). For structures of CA substrates see **Fig. S1.**

**Figure S3:**
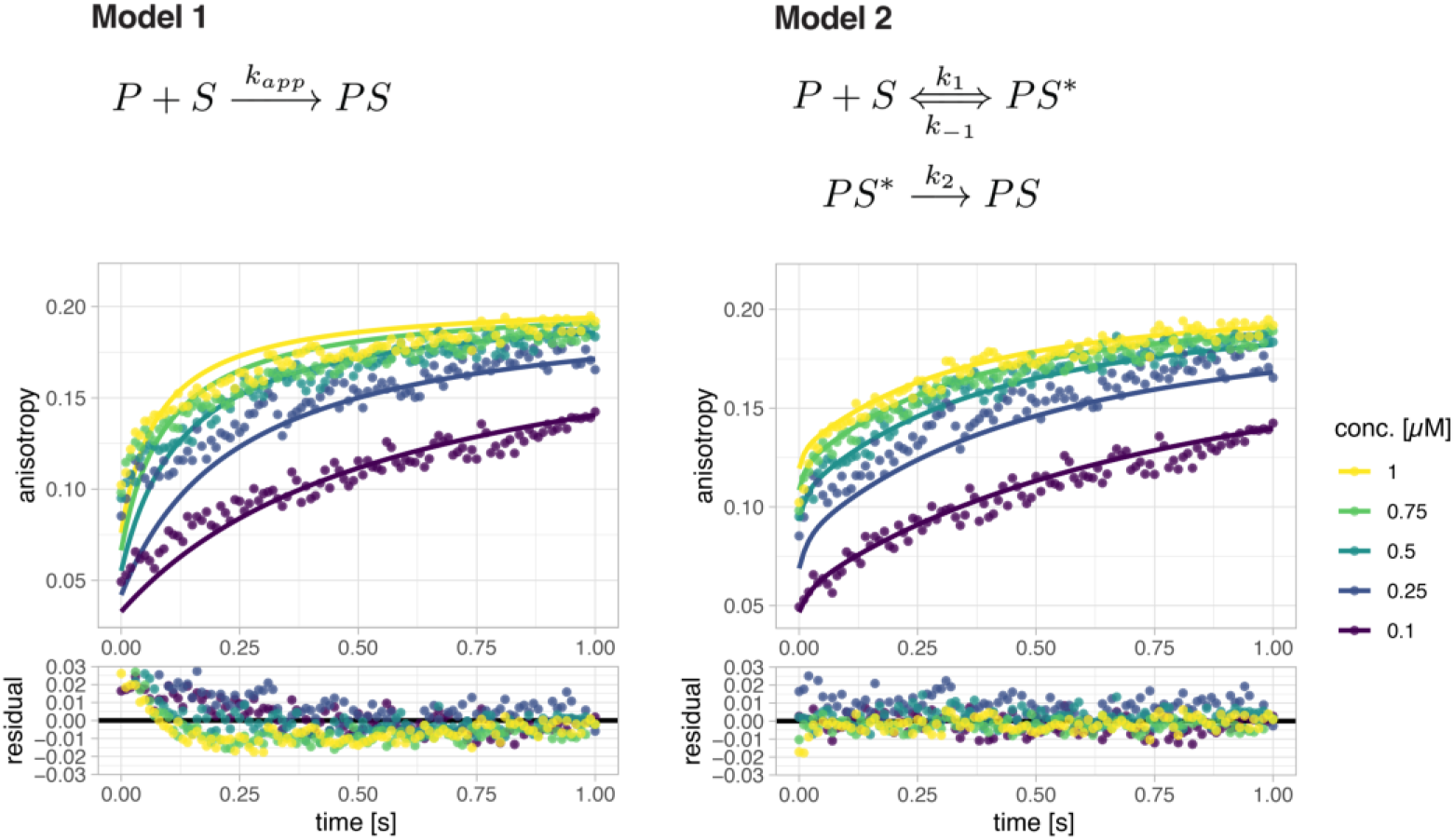
Comparison of model 1 and model 2 fitted to HT7 labeling kinetics. Anisotropy traces (points) and predications of fits based on either model 1 or model 2 (lines) of the labeling reaction between HT7 and CA-TMR are represented in the top panels. Residuals from the fits are depicted in the bottom panels. Kinetics were recorded by following fluorescence anisotropy changes over time using a stopped flow device. All conditions are 1:1 mixtures of protein and substrate at the given concentrations (conc.). Model 2 describes the data better than the simplified model 1. For structures of CA substrates see **Fig. S1.**

**Figure S4:**
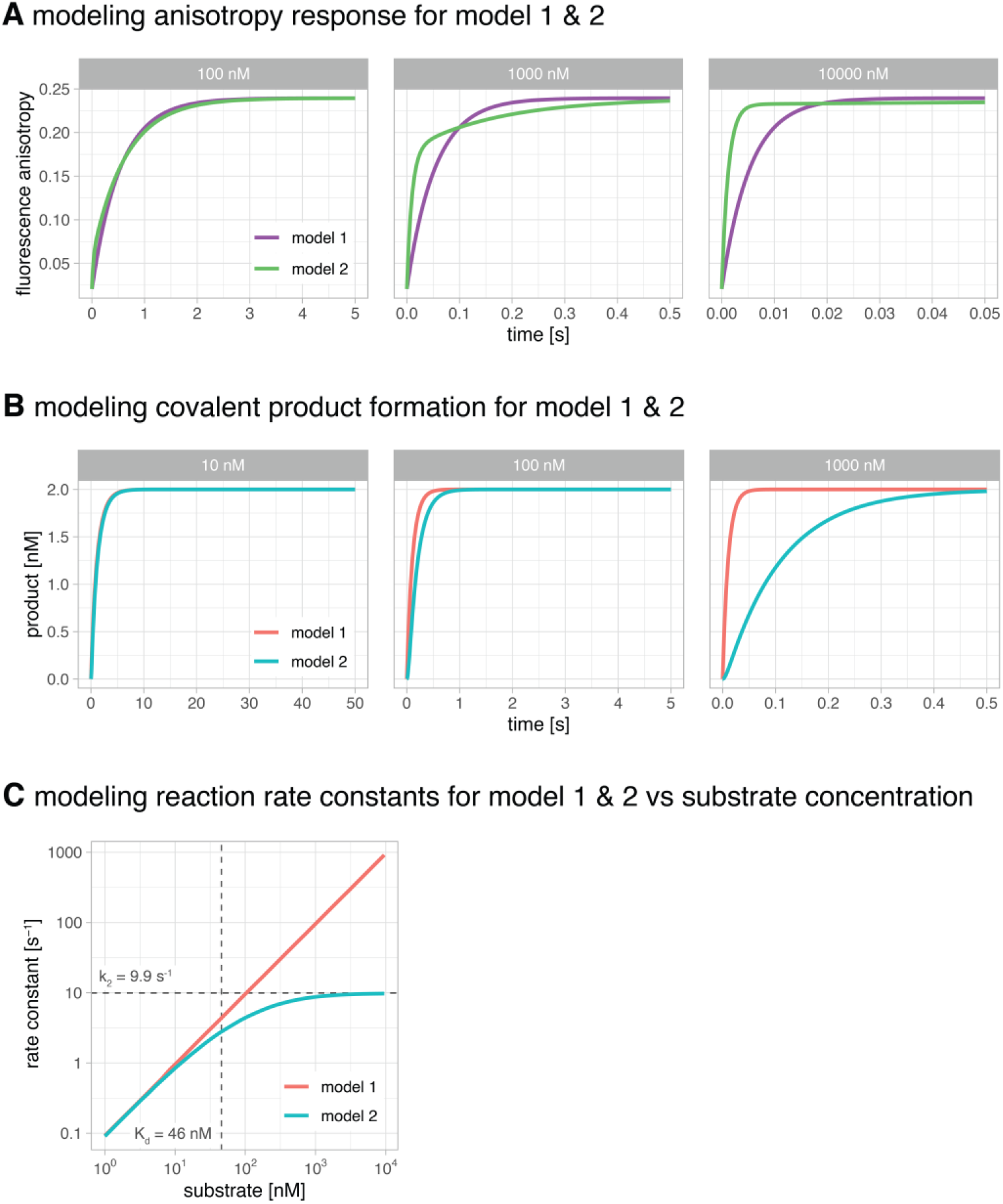
Modeling of HT7 labeling kinetics using measured parameters to compare the kinetic models 1 and 2. **A.** Modeling of the fluorescence anisotropy response at different reactant concentrations using model 1 and 2 with parameters determined for HT7 labeling with CA-TMR. At concentrations below K_d_ (327 nM for CA-TMR) both models yield a rather similar response. At concentrations higher than K_d_ (1000 nM) the response for model 2 shows a strong biphasic character as observed in the measured data, which is not matching the monoexponential behavior of model 1. At very high concentrations (10000 nM) the response for model 2 is again close to a monoexponential curve but the kinetic is much faster than the model 1 curve. This happens since the rise in fluorescence anisotropy for model 2 in the first milliseconds is not due to covalent reaction but mostly binding (k_1_). The binding rate constant k_1_ is faster than k_app_ if k_−1_ is not zero (k_app_ = k_1_ * k_2_ / (k_2_ + k_−1_)). Hence directly estimating k_app_ from fluorescence anisotropy traces by fitting model 1 to the data is only valid for concentrations below K_d_ or if k_−1_ ≪ k_2_. **B.** Modeling the formation of covalently labeled product at different reactant concentrations using model 1 and 2 with parameters determined for HT7 labeling with CA-CPY. At concentrations below K_d_ (46 nM for CA-CPY) both models yield a rather similar behavior. At higher concentrations model 1 predicts a much faster product formation than model 2 since it does not account for enzyme saturation. **C.** Plot of the apparent first order reaction rate constant for product formation against substrate concentration for model 1 and 2 with parameters for CA-CPY. In contrast to model 1, model 2 accounts for enzyme saturation leading to a maximum reaction rate of k_max_ = k_2_ = 9.9 s^−1^. The models start do diverge significantly once the substrate concentration exceeds K_d_ (46 nM). As a consequence, model 2 should be used for predicting formation of labeled HT7 if labeling is performed at high concentrations.

**Figure S5:**
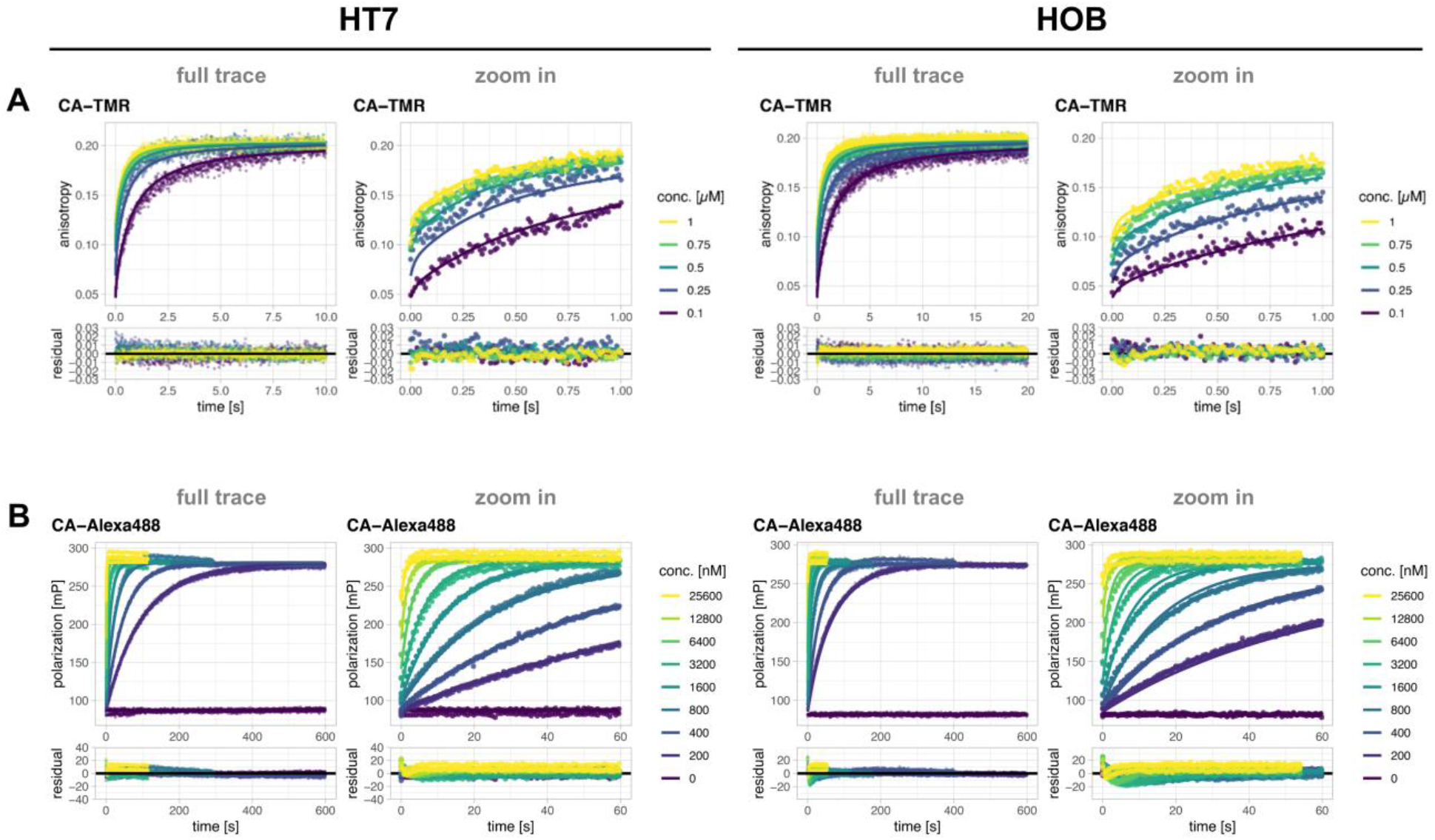
Labeling kinetics of HT7 and HOB with CA-TMR (A) and CA-Alexa488 (B). **A**: Labeling kinetics of HT7 and HOB with CA-TMR. Full anisotropy traces (points) and predications of fits based on model 2 (lines) along with zoom on the initial part are represented on the top panels. Residuals from the fits are depicted in the bottom panels. Kinetics were recorded by following fluorescence anisotropy changes over time using a stopped flow device. All conditions are 1:1 mixtures of protein and substrate at the given concentrations (conc.). **B:** Labeling kinetics of HT7 and HOB with CA-Alexa488. Full fluorescence polarization traces (points) and predications of fits based on model 1 (lines) along with zoom on the initial part are represented on the top panels. Residuals from the fits are depicted in the bottom panels. Kinetics were recorded by following fluorescence polarization changes over time using a plate reader. All experiments were performed at a fixed substrate concentration of 50 nM with varying protein concentrations. For structures of CA substrates see **Fig. S1.**

**Figure S6:**
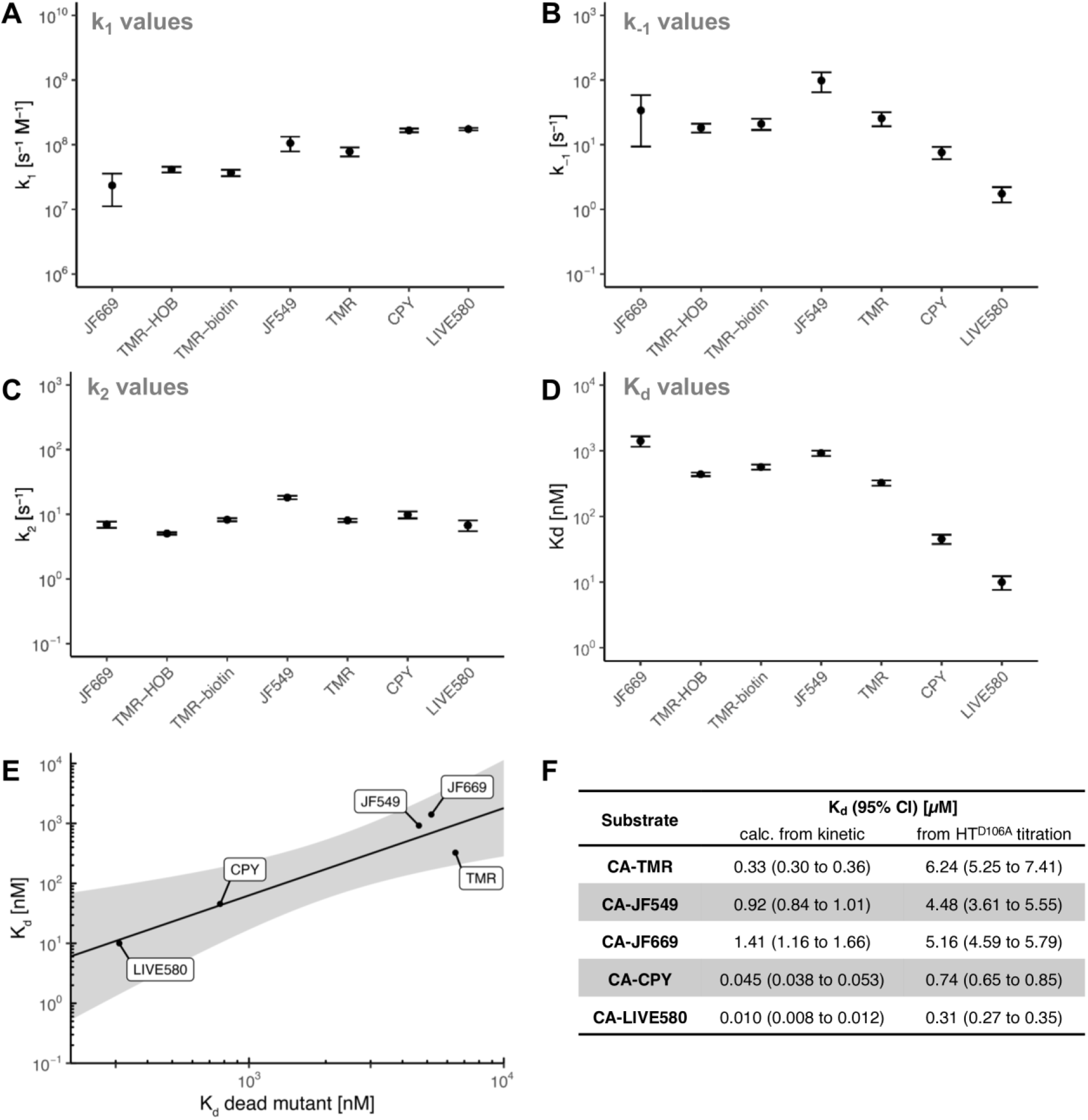
Rate and equilibrium constants of HT7 labeling with various fluorescent CA substrates. Rate constants k_1_ (**A**), k_−1_ (**B**), k_2_ (**C**) and the calculated dissociation constants (K_d_ = k_−1_/k_1_, **D.**) obtained from fitting model 2 to stopped flow labeling experiments of HT7 and HOB. The catalytic rate constant (k_2_) is rather constant among these substrates, while there are significant differences in the dissociation constant (K_d_). The K_d_ variations are due to large differences in k_−1_ and minor differences in k_1_. As a result, differences in k_app_ can be mostly explained by affinity differences of HT7 towards its substrates. **E.** Correlation between the calculated K_d_ from the stopped flow kinetic experiments and the K_d_ obtained from titration experiments performed with the dead mutant HT7^D106A^. Log transformed values were fitted to a linear model (log(y) = 1.455 * log(x) - 2.567; black line, 95% confidence bands in grey, depicting the area in which the true regression line lies with 95% confidence). The linear correlation in logarithmic space suggests that the K_d_ of CA rhodamine substrates with HT7^D106A^ could represent a valid proxy to estimate their K_d_ with the native HT7. **F** K_d_ values of the tested substrates calculated from the kinetics (k_1_/k_−1_) and measured by fluorescence polarization titration against the dead mutant _HT_D106A.

**Figure S7:**
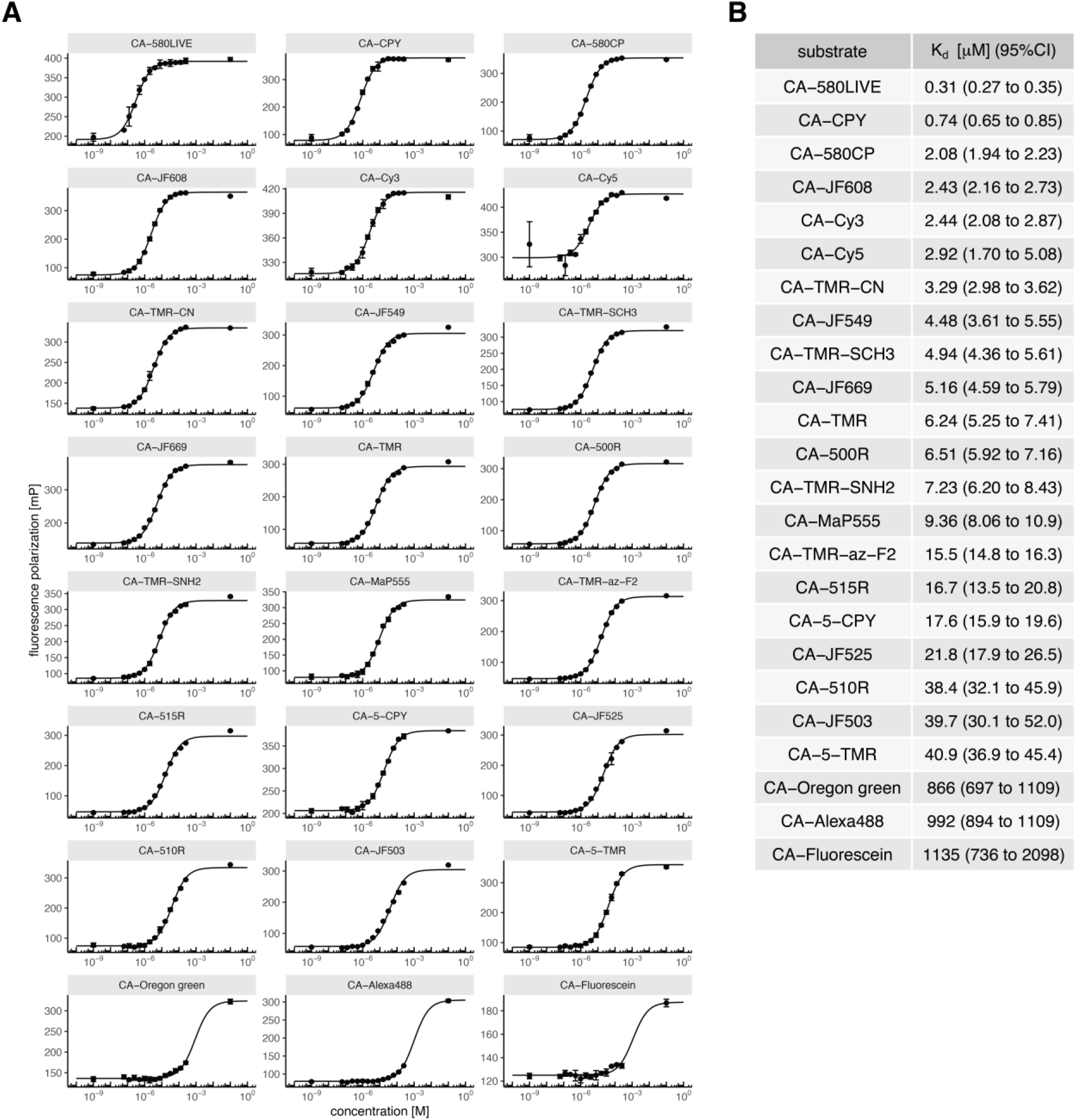
Affinity of the dead mutant HT7^D106A^ to fluorescent CA substrates. **A**. Titration curves of fluorescent CA substrates against HT7^D106A^ measured via fluorescence polarization. The FP value of each dye fully bound to native HT7 was added at c = 0.1 M to improve fitting of the upper plateau. (See corresponding methods section for more details). **B.** Table summarizing fitted K_d_ values with 95% confidence intervals. For structures of CA substrates see **Fig. S1.**

**Figure S8:**
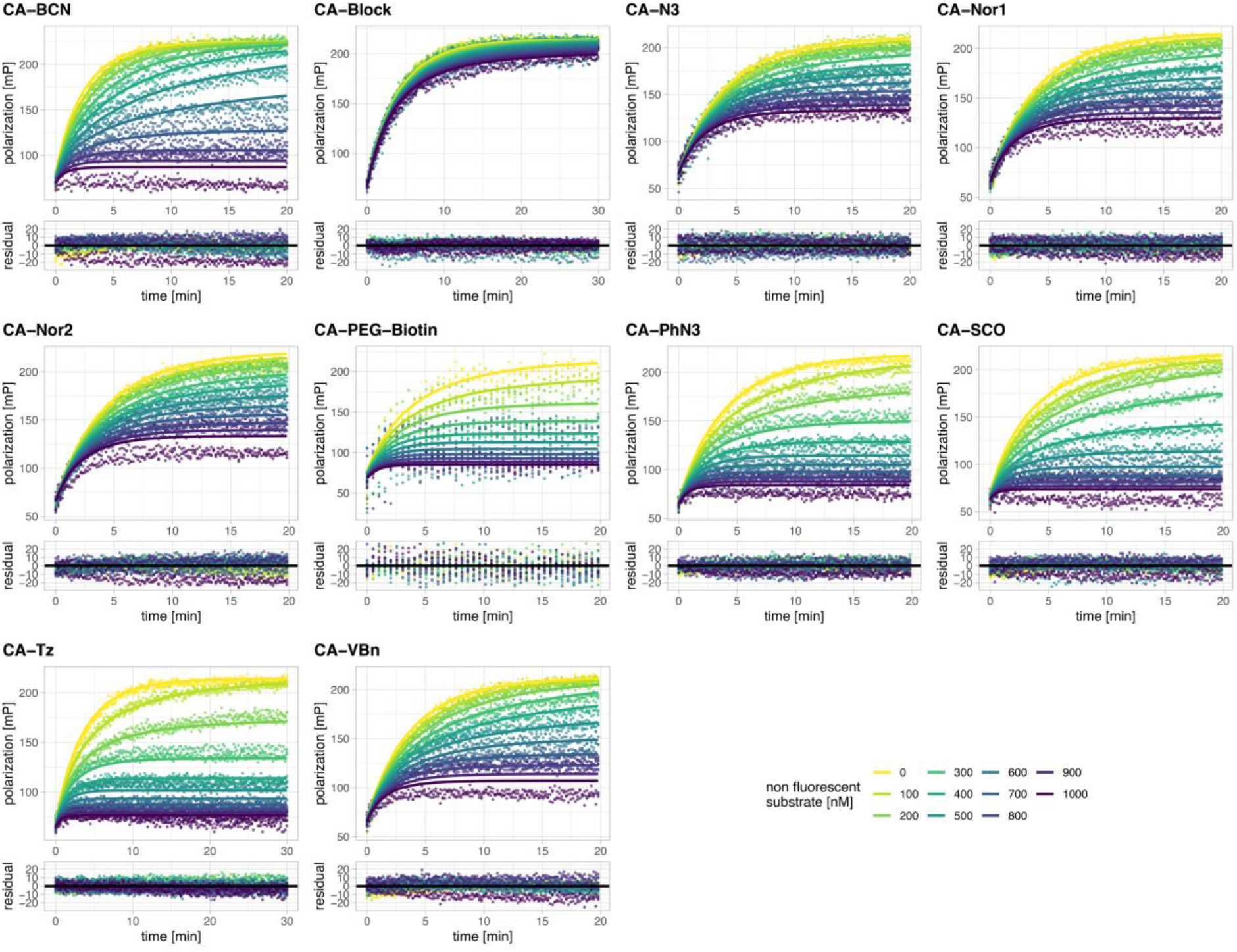
Labeling kinetics of HT7 with non-fluorescent CA substrates. Fluorescence polarization traces (points) of kinetic competition assays and predications of fits (lines) based on a simple competitive model (see methods section for details) of HT7 labeling with CA-Alexa488 in the presence of different concentrations of non-fluorescent CA substrates are represented on the top panels. Residuals from the fits are depicted in the bottom panels. Kinetics were recorded by following fluorescence polarization changes over time using a plate reader. For structures of CA substrates see **Fig. S1.**

**Figure S9:**
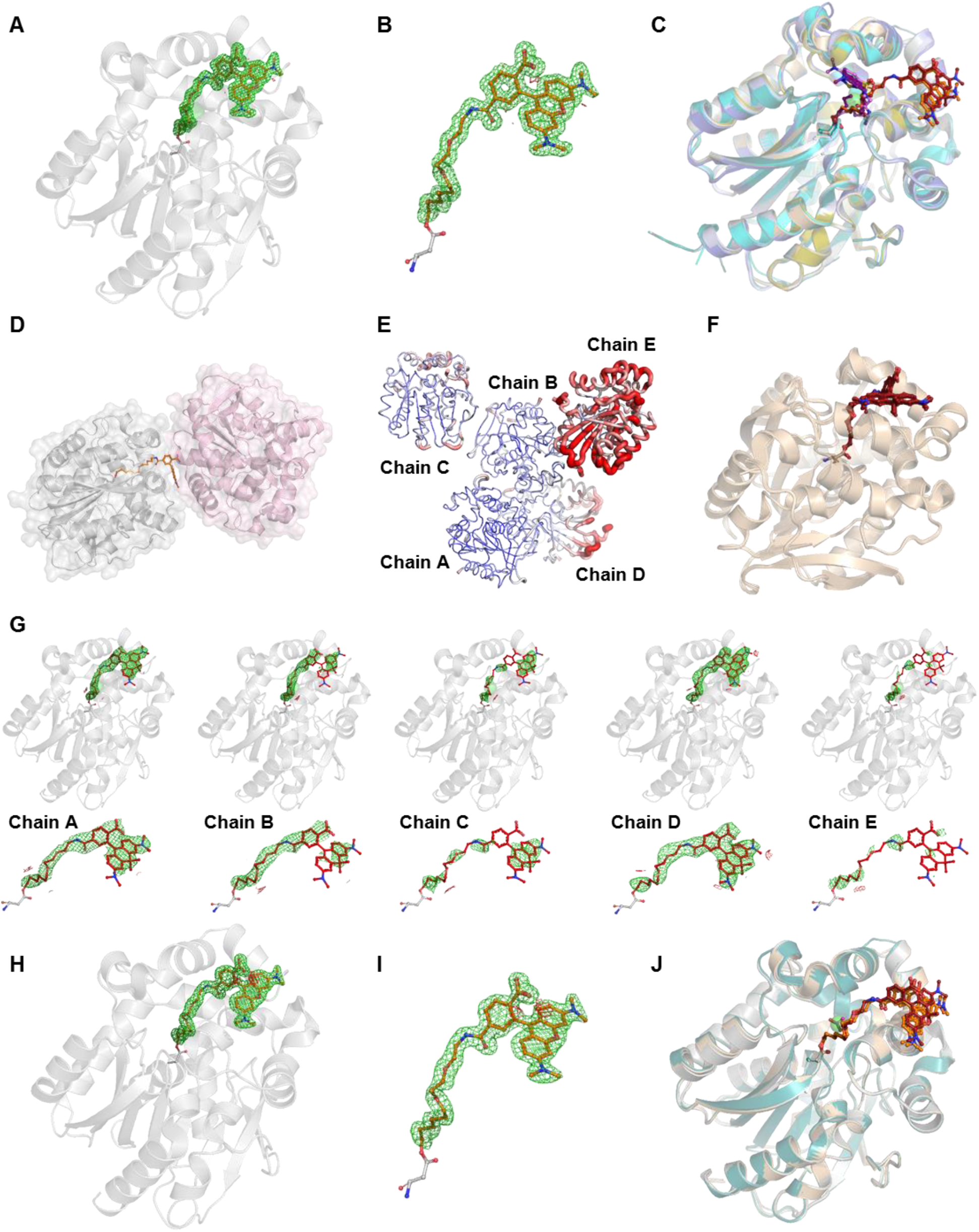
Validation of HT7-TMR and HT7-CPY X-ray structures. **A**. Omit-map of the TMR ligand of the HT7-TMR X-ray structure. The protein is represented as grey cartoon with the catalytic aspartate (grey) and the TMR ligand (orange) represented as sticks. **B.** Zoom on the isolated labeled catalytic aspartate of HT7-TMR. The omit map of the alkane-TMR is contoured at 3 σ and represented as green and red mesh for missing and extra density, respectively. **C**. Structure alignment of HT7-TMR and HT7-CPY (chain A) with different X-ray structures of HaloTag. All structures are represented as cartoons with their respective catalytic aspartate and ligands represented as sticks. When present the chloride is represented as green sphere. **D**. Alkane-TMR constraints by the crystal packing. Two monomers of HT7-TMR are represented as grey and light-pink cartoons and surfaces. The conformation of alkane-TMR (orange sticks) of the grey monomer is constrained by the light-pink monomer that was generated as symmetry mate. **E**. B-factor putty representation of the different chains of the asymmetric unit of the HT7-CPY crystal structure. Blue = 15; Red = 120. Chain E and to a lesser extent Chain D present an overall higher B-factor compared the other monomers. **F**. Structure alignment of the different monomers in the asymmetric unit of the HT7-CPY structure. The monomers are represented as wheat cartoon with the catalytic aspartate and alkane-CPY represented as sticks; all featuring similar conformations. **G**. Omit-maps of the alkane-CPY ligands of the different monomers in the HT7-CPY asymmetric unit. Proteins are represented as grey cartoons with the catalytic aspartates (grey) and alkane-CPYs (firebrick) represented as sticks. **H**. Omit-map of the TMR ligand of the HOB-TMR X-ray structure. The protein is represented as grey cartoon with the catalytic aspartate (grey) and the TMR ligand (orange) represented as sticks. **I.** Zoom on the isolated labeled catalytic aspartate of HOB-TMR. The omit map of the alkane-TMR is contoured at 3 σ and represented as green and red mesh for missing and extra density, respectively. **J**. Structure alignment of HT7-TMR, HT7-CPY (chain A) and HOB-TMR. Proteins are represented as grey, wheat and dark-green cartoons, respectively. The alkane-TMR and −CPY ligands are represented as orange and firebrick sticks, respectively. The catalytic aspartate is represented as sticks of the same color as the cartoons.

**Figure S10:**
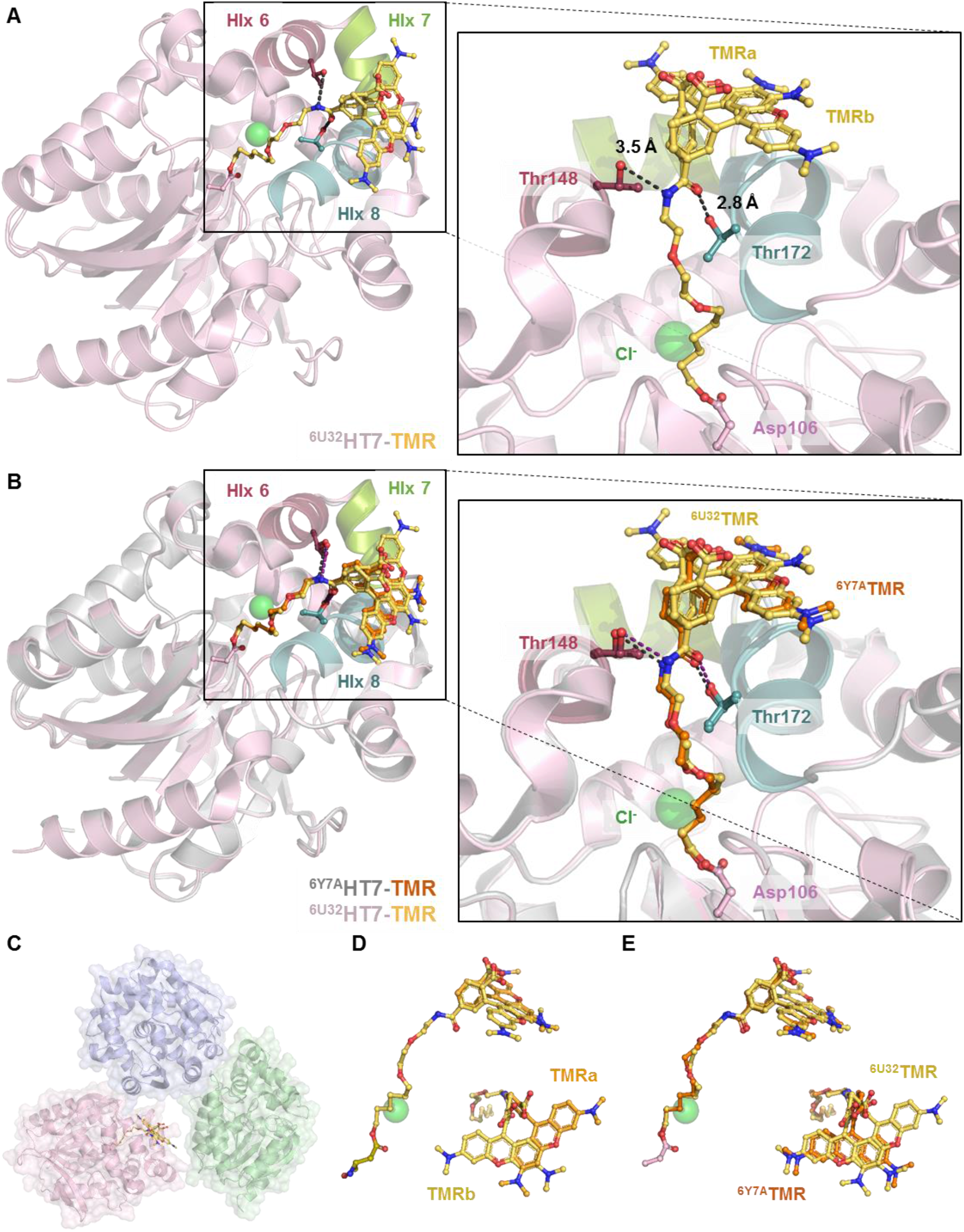
Structural comparison between HT7-TMR structures from PDB IDs 6U32 and 6Y7A. **A**. Structure of HT7-TMR (PDB ID 6U32, previously published (16)) featuring two conformations of the alkane-TMR ligand **B**. Structural comparison between ^6U32^HT7-TMR and ^6Y7A^HT7-TMR (PDB ID 6Y7A, this study). Hydrogen bonds between ^6Y7A^HT7-TMR and ^6U32^HT7-TMR and their respective reacted substrates are represented as black and dark-purple lines, respectively. **C**. ^6U32^HT7-TMR crystal packing. Three monomers of HT7-TMR are represented as blue, green and pink cartoons. The conformation of the alkane-TMR (yellow/orange sticks) of the pink monomer is not constrained by the other symmetry mates. **D**. Zoom on the catalytic aspartate and alkane-TMR substrate highlighting the alternative conformations observed in the ^6U32^HT7-TMR crystal structure. The two alternative TMR conformations (a and b) are represented as different tone of yellow/orange sticks. **E**. Structural comparison of the substrate positioning between ^6U32^HT7-TMR and ^6Y7A^HT7-TMR. Alkane-^6U32^TMR and alkane-^6Y7A^TMR are represented as yellow and orange sticks, respectively. The ^6Y7A^TMR present a very similar conformation than one of the ^6U32^TMR conformation which can’t be observed due to the crystal packing in the ^6Y7A^HT7-TMR crystal structure.

**Figure S11:**
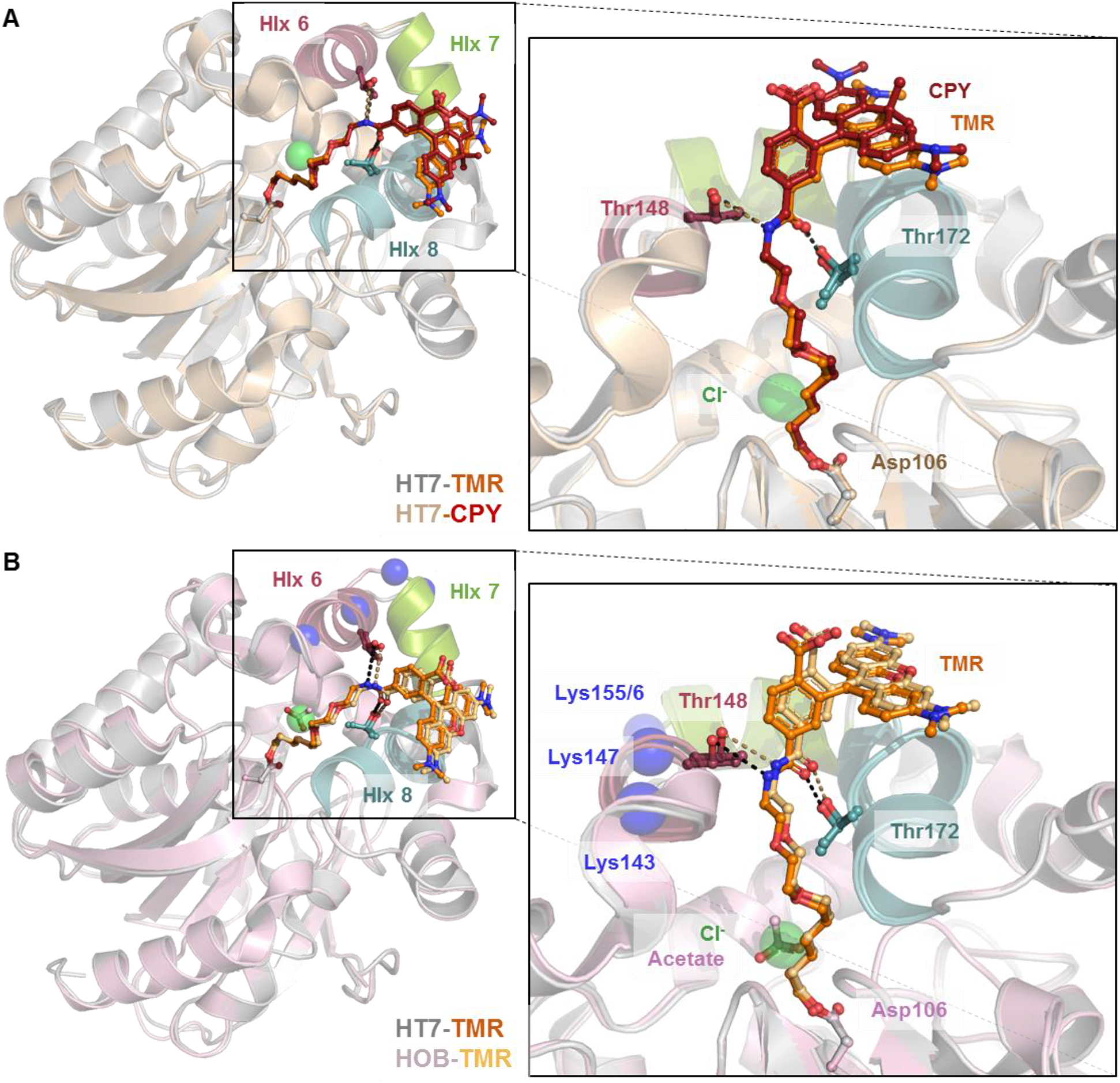
Structural comparison between HT7-TMR, HT7-CPY (A) and HOB-TMR (B). **A.** Structural comparison between HT7-TMR and HT7-CPY. Hydrogen bonds between HT7-TMR and HT7-CPY and their respective reacted substrates are represented as black and sand dashed lines, respectively. **B**. Structural comparison between HT7-TMR and HOB-TMR. Hydrogen bonds between HT7 and HOB and their respective reacted substrates are represented as black and sand dashed lines, respectively.

**Figure S12:**
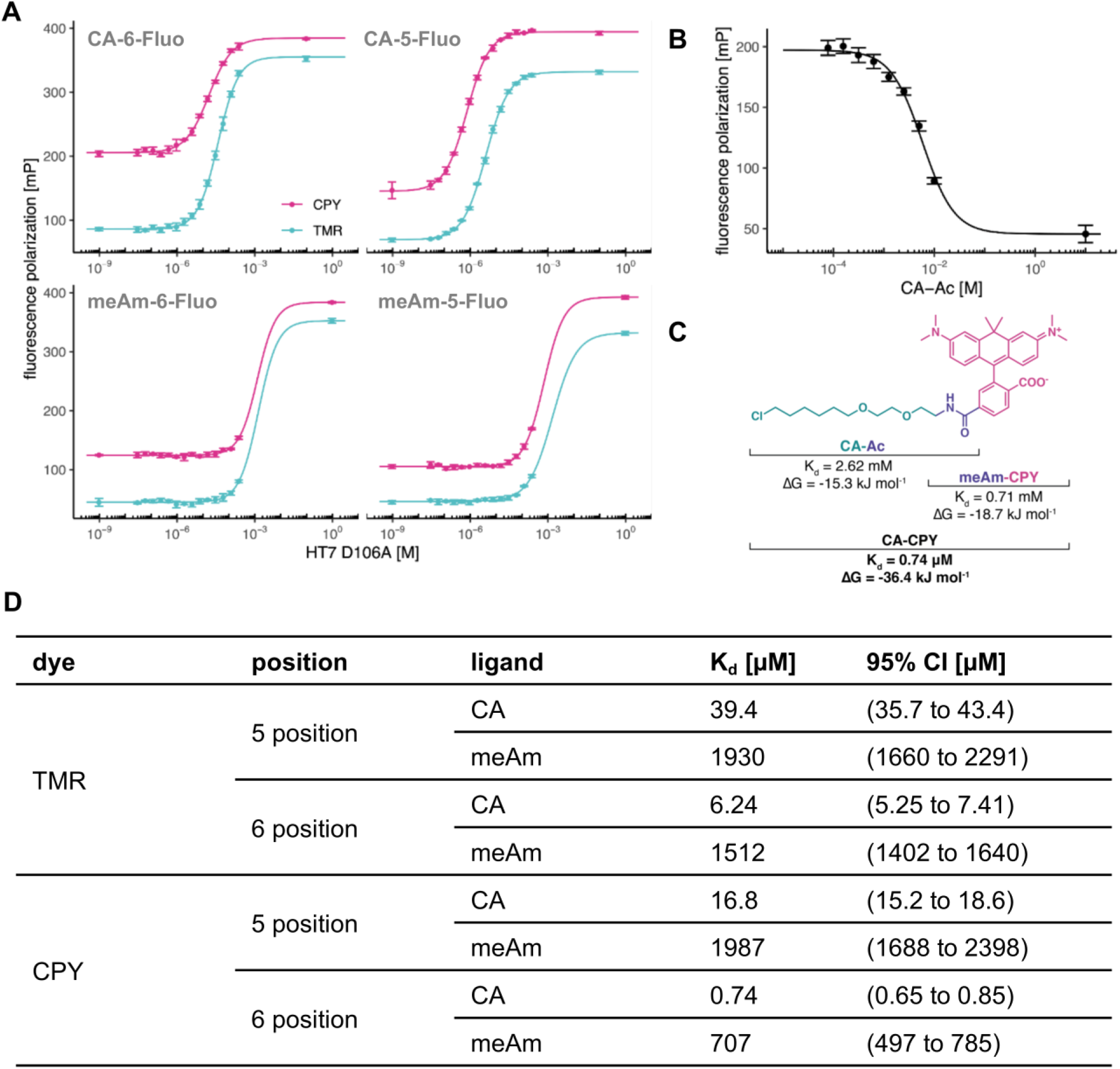
Biochemical study of the interaction of HT7 with CA-fluorophores. **A.** Affinity of the dead mutant HT7^D106A^ towards different fluorophore derivatives measured via fluorescence polarization assay. The FP value of each dye as CA substrate fully bound to native HT7 was added at c = 0.1 M or c = 1 M in order to improve fitting of the upper plateau. **B.** Affinity of HT7^D106A^ to CA-Ac measured via fluorescence polarization competition assay against CA-TMR. **C.** Summary of dissociation constants (K_d_) and calculated free binding energies (ΔG) of HT7^D106A^ with CA-Ac, mAm-5-CPY and CA-5-CPY. The representation highlights the additive nature of the binding energies from the chloroalkane and the CPY moieties for the binding energy of the full substrate. **D.** Table summarizing values and confidence intervals (95%) of the fits.

**Figure S13:**
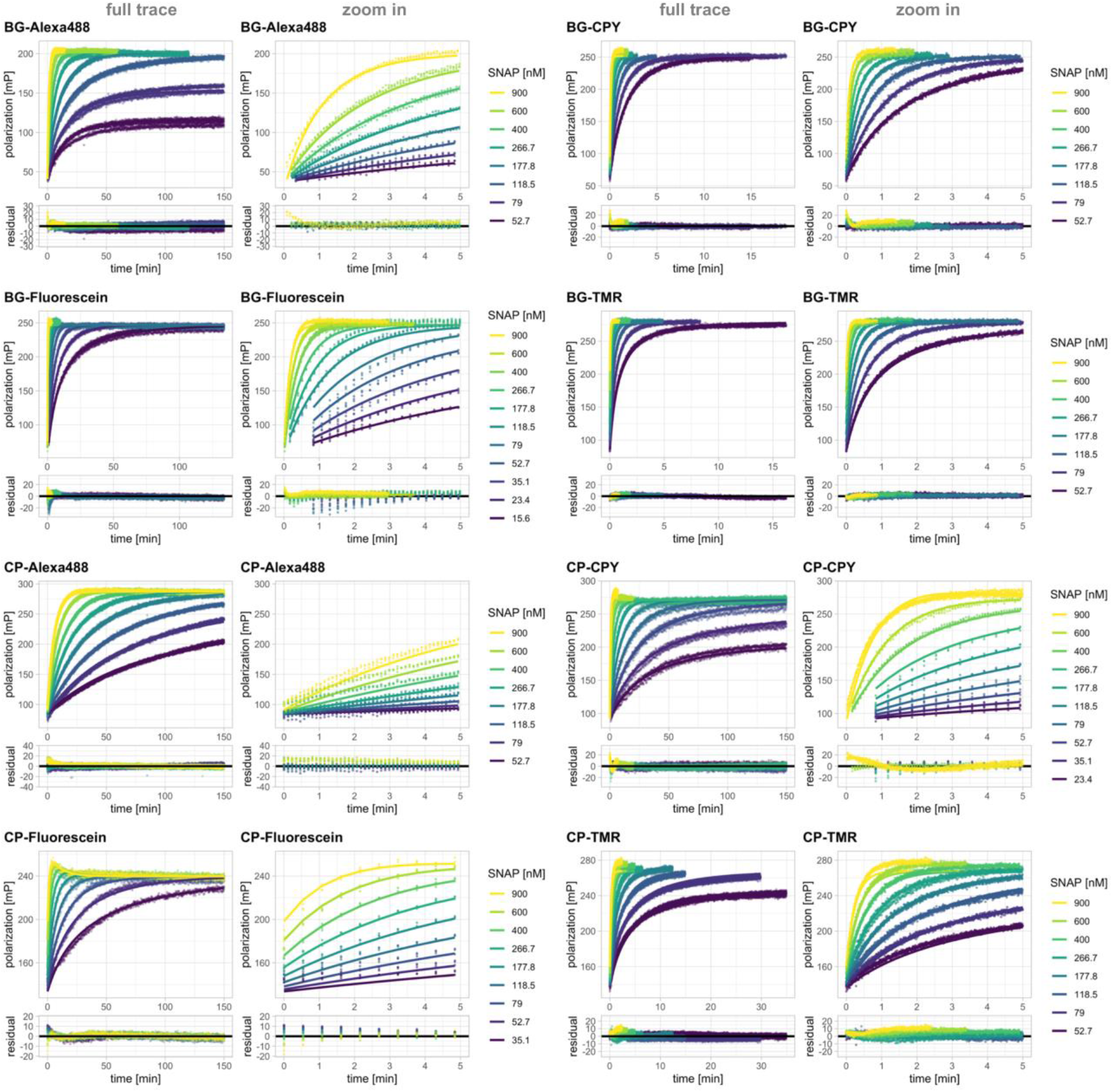
Labeling kinetics of SNAP with fluorescent BG and CP substrates. Full fluorescence polarization traces (points) and predications of fits based on model 1 or 1.2 (lines) along with zoom on the initial 5 minutes are represented on the top panels. Most substrates were fitted to model 1 except CP-Fluorescein and CP-CPY, which showed an additional phase (model 1.2). Residuals from the fits are depicted in the bottom panels. Kinetics were recorded by following fluorescence polarization changes over time using a plate reader. Labeling was performed at different concentrations of SNAP protein. Substrate concentrations were aimed at 20 nM based on the dyes extinction coefficient but fitted in the model since significant deviations from the expected stoichiometry were observed. For structures of BG and CP substrates see **Fig. S1.**

**Figure S14:**
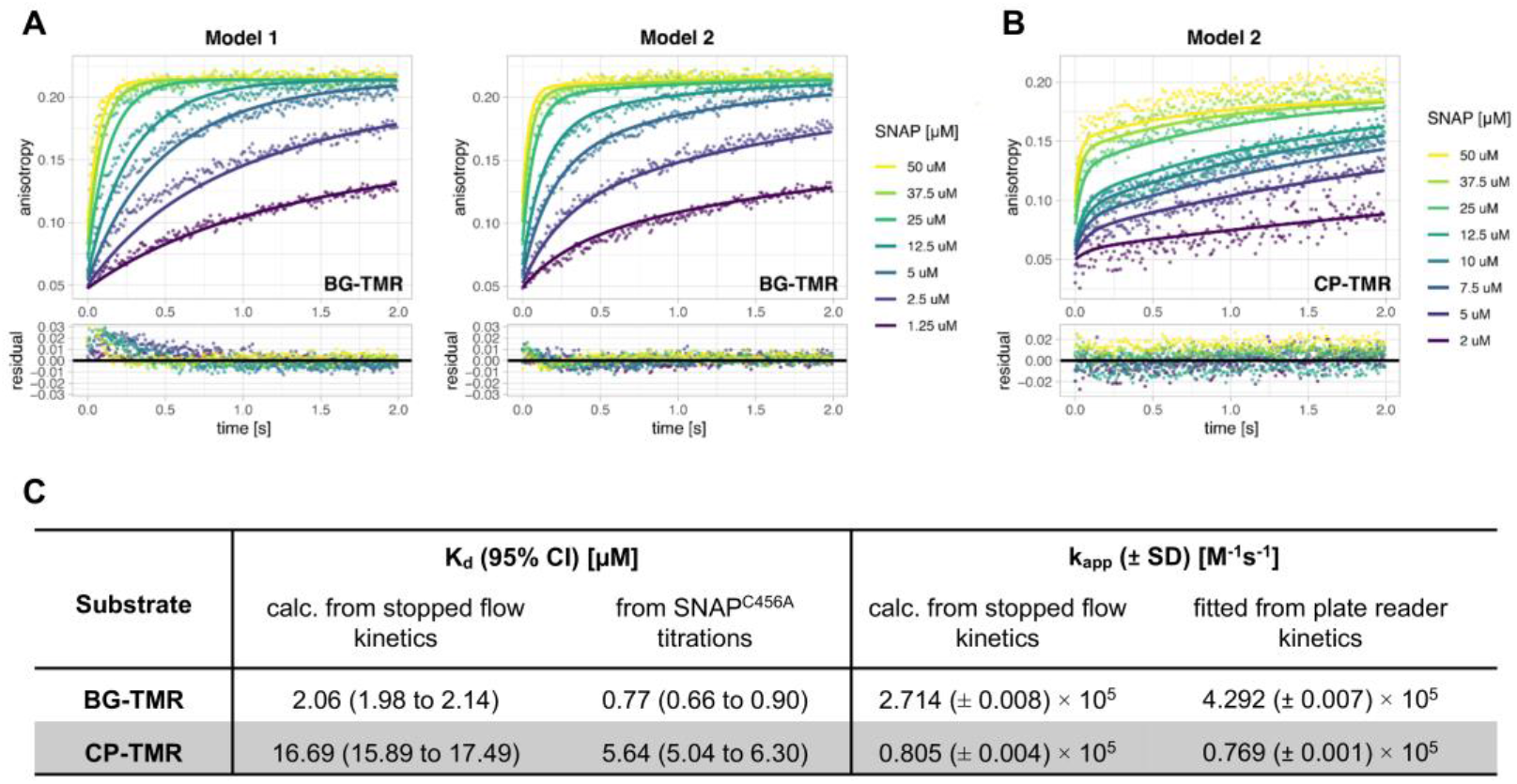
Labeling kinetics of SNAP measured by stopped flow fluorescence anisotropy. **A**. Comparative data analysis of SNAP labeling kinetics with BG-TMR. Anisotropy traces (points) and predications of fits based on either model 1 or model 2 (lines) of the labeling reaction between SNAP and BG-TMR are represented in the top panels. Residuals from the fits are depicted in the bottom panels. Labeling was performed at different concentrations of SNAP protein and a constant substrate concentration of 1 μM. Model 2 describes the data better than the simplified model 1. (for model description see **Fig. 1**). **B.** Kinetic traces of SNAP labeling with CP-TMR represented as previously explained and fit with model 2. For structures of BG and CP substrates see **Fig. S1. C.** K_d_ and k_app_ values calculated from parameters obtained by fitting model 2 to stopped flow anisotropy data (K_d_ = k_−1_/k_1_, k_app_ = k_1_*k_2_/(k_−1_+k_2_)) compared to values directly fitted to fluorescence polarization assay with SNAP^C145A^ (K_d_) and plate reader kinetics at lower SNAP concentrations fitted with model 1 (k_app_).

**Figure S15:**
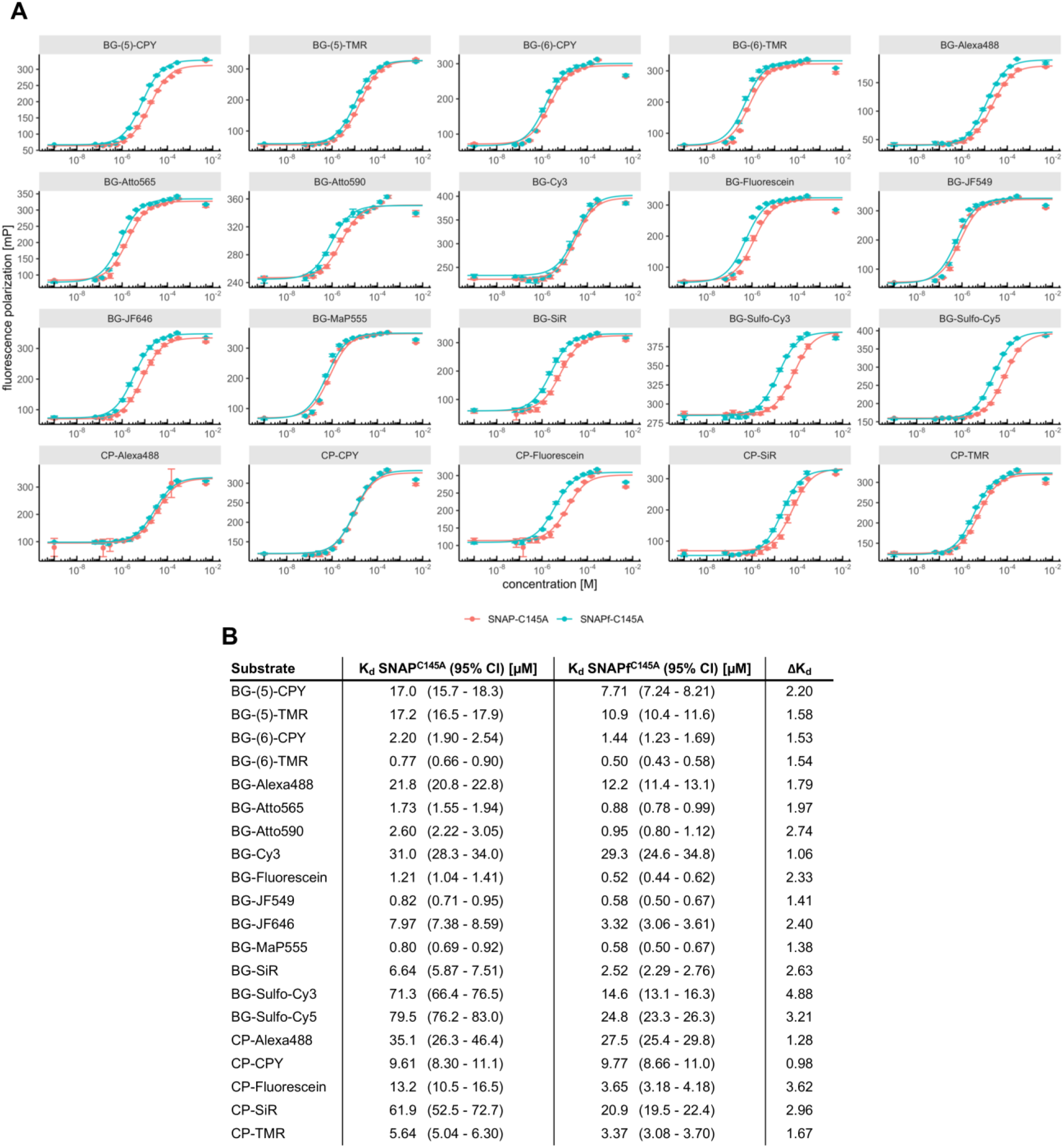
Comparison of fluorophore substrate affinities between the dead mutants SNAP^C145A^ and SNAPf^C145A^. **A**. Titration curves obtained for the dead mutants SNAP^C145A^ and SNAPf^C145A^ measured via fluorescence polarization. The FP value of each dye fully bound to native SNAP/SNAPf was added at c = 0.005 M to improve fitting of the upper plateau. (See corresponding methods section for more details). **B**. Table summarizing fitted K_d_ values with 95% confidence intervals. For structures of BG and CP substrates see **Fig. S1.**

**Figure S16:**
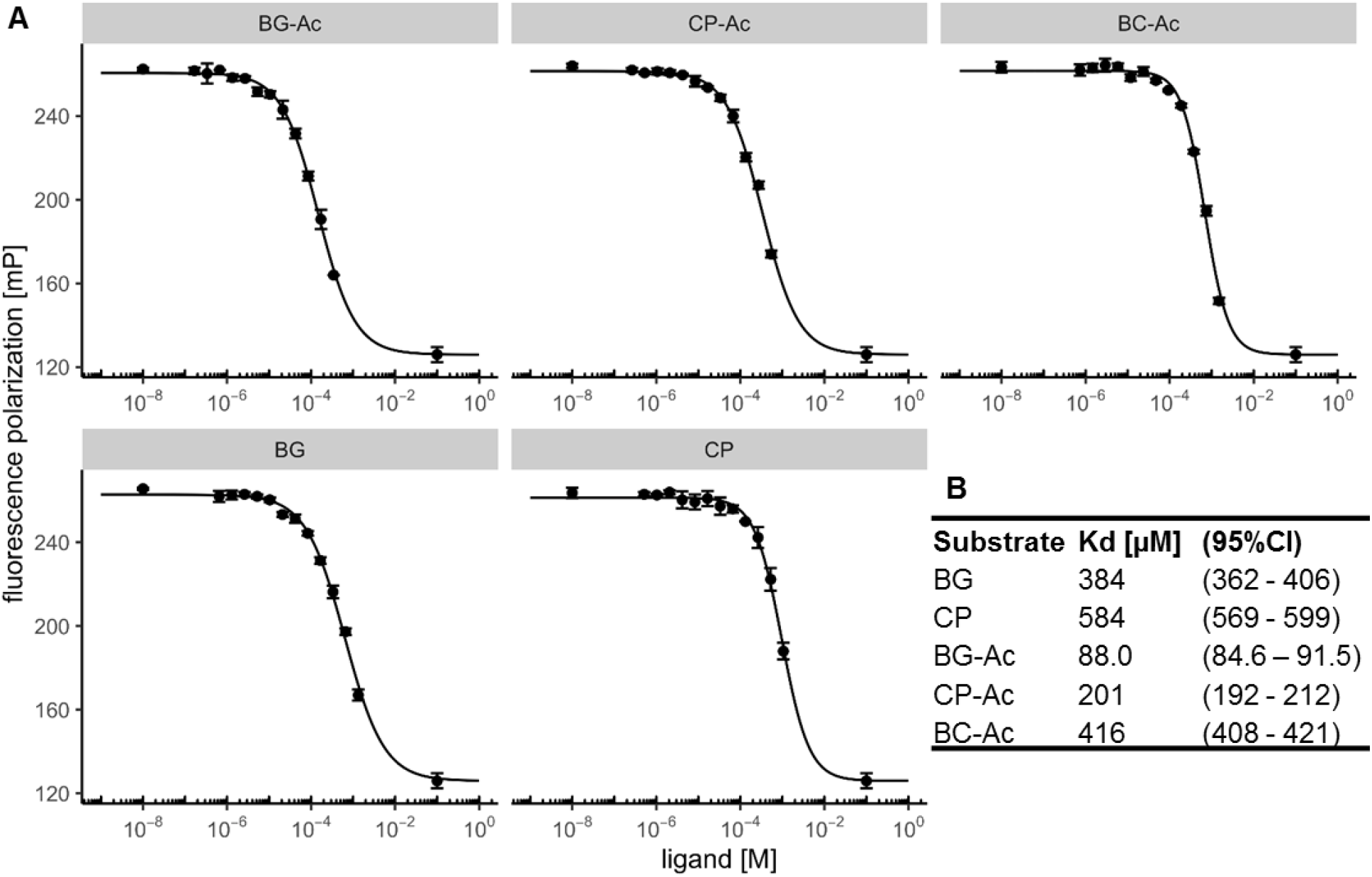
Comparison of non-derivatized core substrate affinities with the dead mutant SNAP^C145^. **A**. Titration curves obtained for the dead mutant SNAP^C145A^ measured via competitive fluorescence polarization. The FP value of free dye was added at c = 0.1 M to improve fitting of the lower plateau. (See corresponding methods section for more details) **B**. Table summarizing fitted K_d_ values with 95% confidence intervals. For structures of substrates see **Fig. S1.**

**Figure S17:**
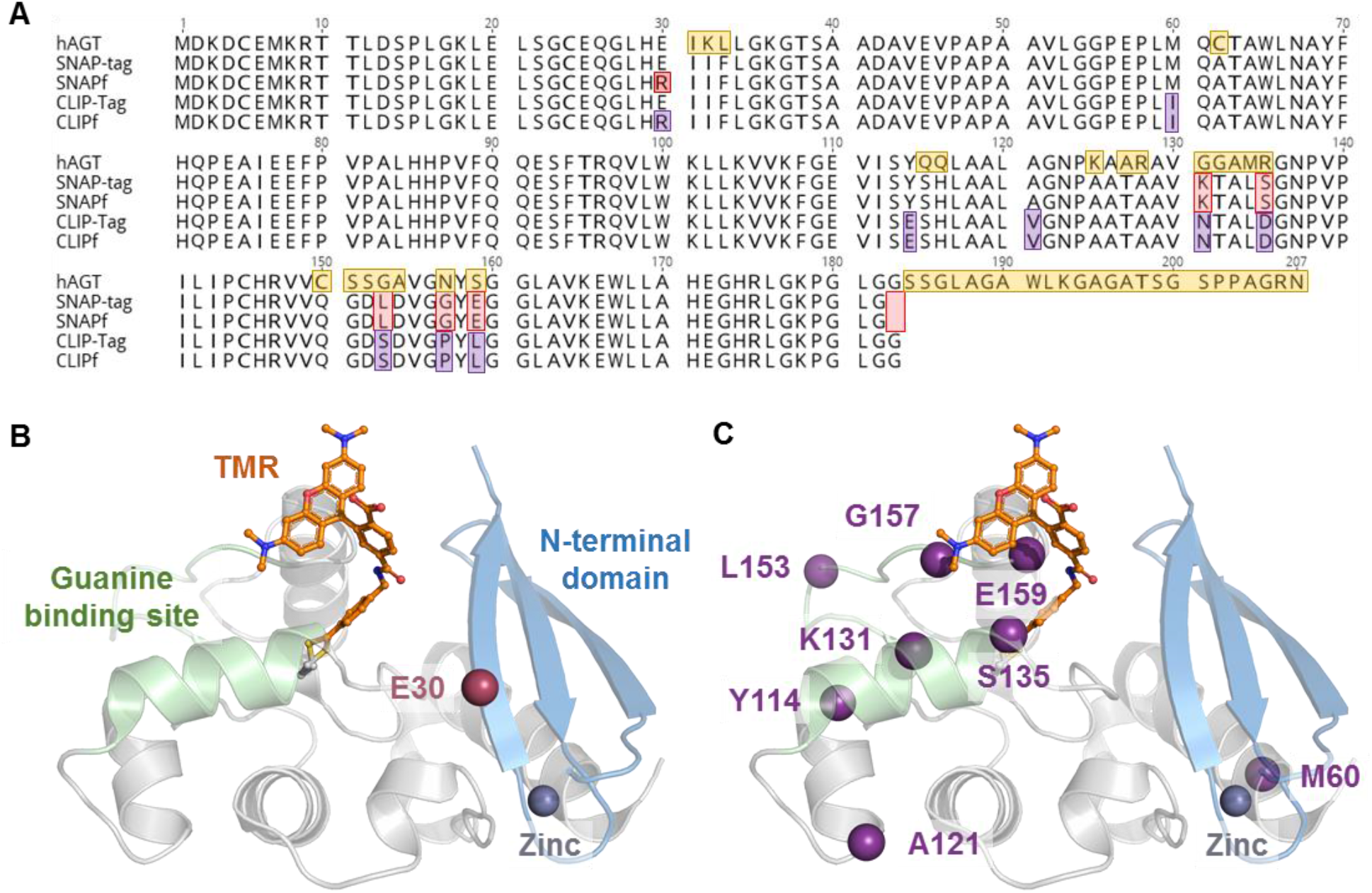
Sequence alignment and structural comparison between SNAP and CLIP variants. **A.** Sequence alignment of hAGT, SNAP, SNAPf, CLIP and CLIPf. Differences are highlighted in yellow, red and violet in the hAGT, SNAP(f) and CLIP(f) sequences, respectively. **B.** Crystal structure of SNAP labeled with TMR. SNAP is represented as grey cartoon despite for the BG binding site and the N-terminal domain that are represented in green and blue, respectively. The catalytic cysteine is represented as grey sticks and the benzyl-TMR as orange sticks. The residue E30 which is mutated to an arginine (R) in SNAPf is highlighted as a red sphere. **C.** Crystal structure of SNAP labeled with TMR with α-carbons of the residues that differ between SNAP and CLIP represented as purple spheres.

**Figure S18:**
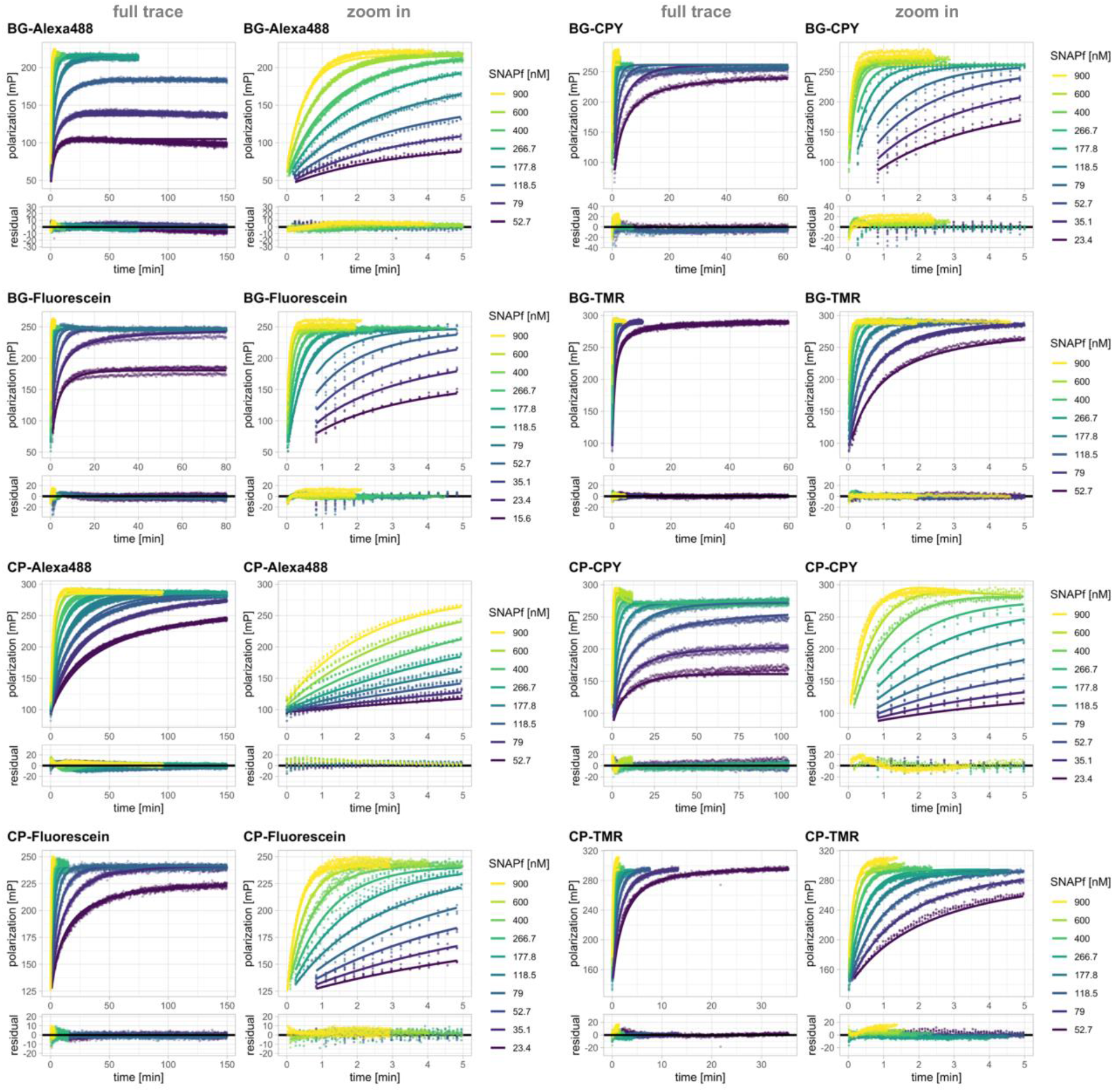
Labeling kinetics of SNAPf with fluorescent BG and CP substrates. Full fluorescence polarization traces (points) and predications of fits based on model 1 or 1.2 (lines) along with zoom on the initial 5 minutes are represented on the top panels. All substrates were fitted to model 1 except CP-CPY, which showed an additional phase (model 1.2). Residuals from the fits are depicted in the bottom panels. Kinetics were recorded by following fluorescence polarization changes over time using a plate reader. Labeling was performed at different concentrations of SNAPf protein. Substrate concentrations were aimed at 20 nM based on the dyes extinction coefficient but fitted in the model since significant deviations from the expected stoichiometry were observed. For structures of BG and CP substrates see **Fig. S1.**

**Figure S19:**
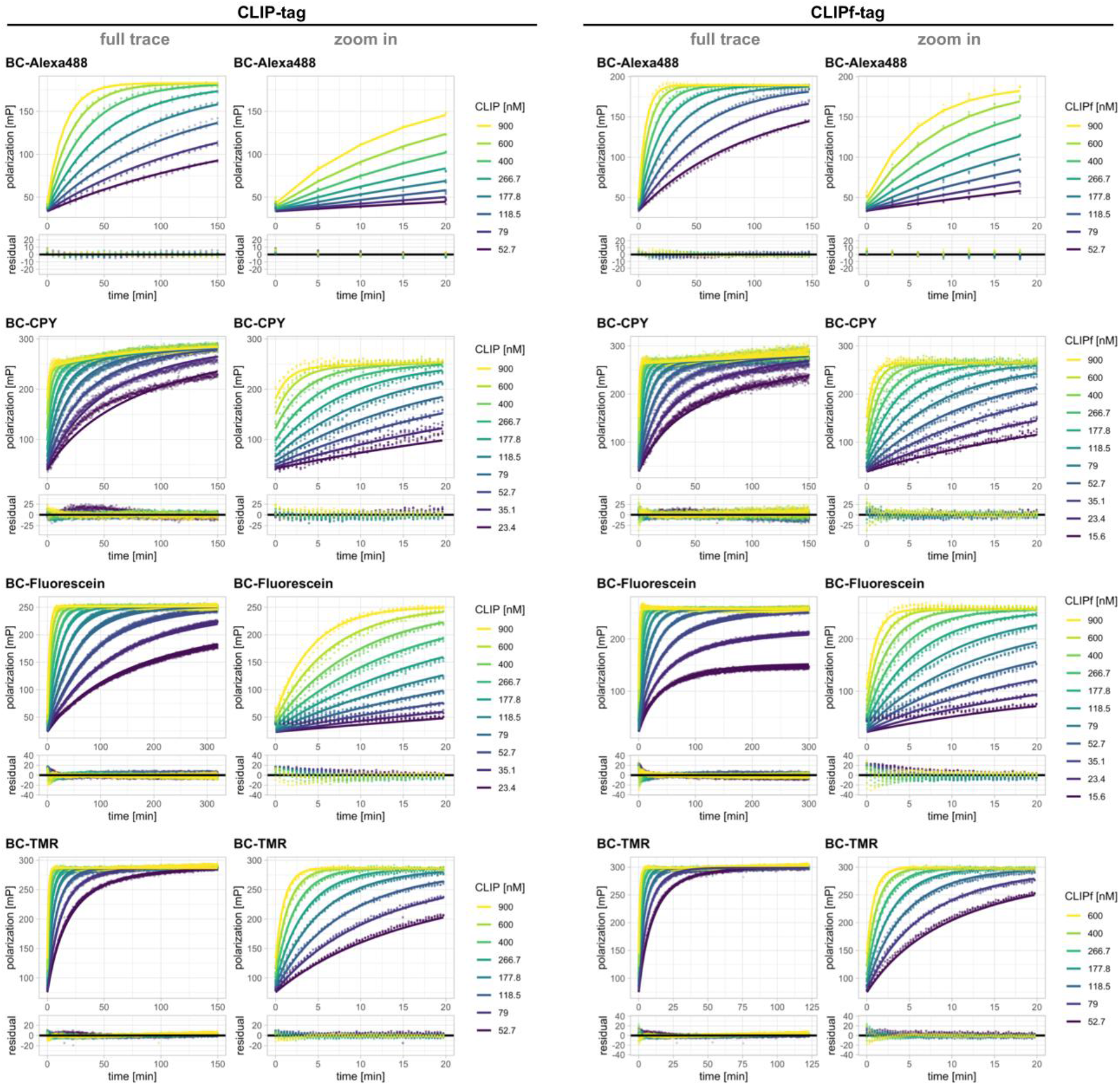
Labeling kinetics of CLIP and CLIPf with fluorescent BC substrates. Full fluorescence polarization traces (points) and predications of fits based on model 1 or 1.2 (lines) along with zoom on the initial 20 minutes are represented on the top panels. All substrates were fitted to model 1 except BC-CPY, which showed an additional phase (model 1.2). Residuals from the fits are depicted in the bottom panels. Kinetics were recorded by following fluorescence polarization changes over time using a plate reader. Labeling was performed at different concentrations of CLIP and CLIPf protein. Substrate concentrations were aimed at 20 nM based on the dyes extinction coefficient but fitted in the model since significant deviations from the expected stoichiometry were observed. For structures of BC substrates see **Fig. S1**.

**Figure S20:**
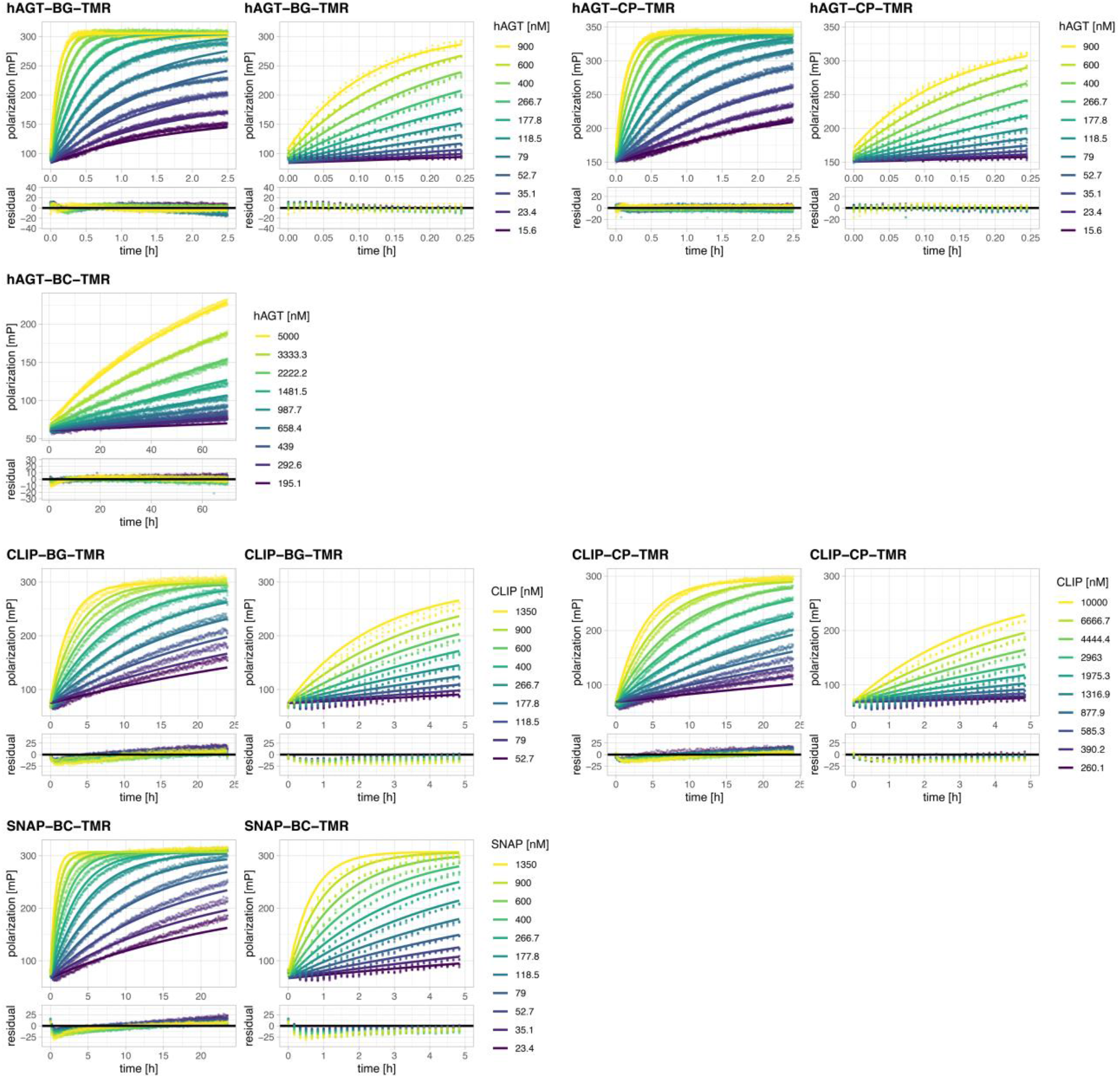
Labeling kinetics of hAGT, SNAP and CLIP with the non-respective BG-, CP- and BC-TMR substrates. Full fluorescence polarization traces (points) and predications of fits based on model 1 along with zoom on the initial part (except for BC-TMR and hAGT) are represented on the top panels. Residuals from the fits are depicted in the bottom panels. Kinetics were recorded by following fluorescence polarization changes over time using a plate reader. Labeling was performed at different concentrations of hAGT, SNAP and CLIP proteins. Substrate concentrations were aimed at 20 nM based on the dyes extinction coefficient but fitted in the model since significant deviations from the expected stoichiometry were observed. For structures of BG, CP and BC substrates see **Fig. S1**.

**Figure S21:**
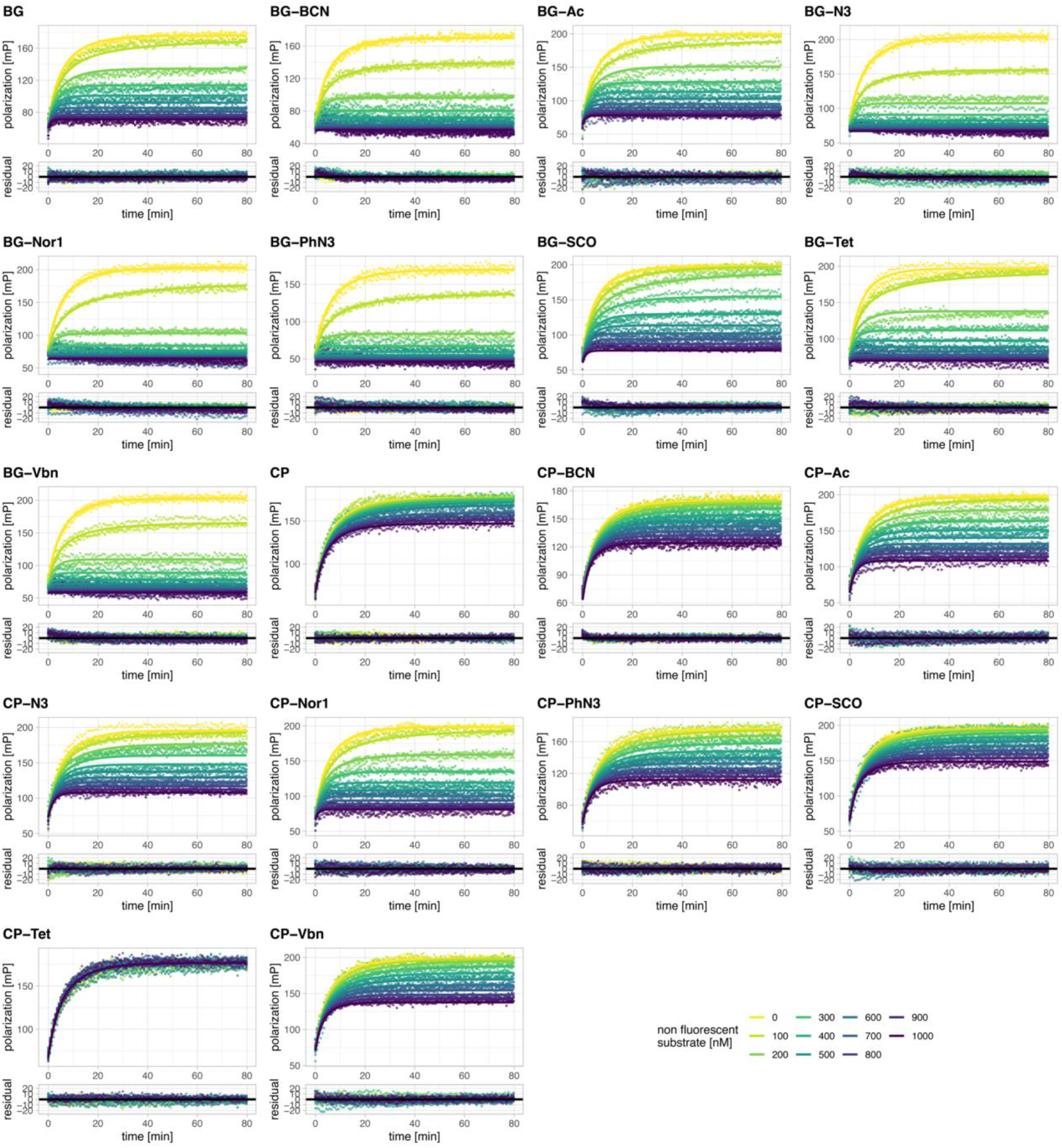
Labeling kinetics of SNAP with non-fluorescent BG and CP substrates. Fluorescence polarization traces (points) of kinetic competition assays and predications of fits based on a simple competitive model (lines, see methods section for details) of SNAP labeling with BG-Alexa488 in the presence of different concentrations of non-fluorescent BG/CP substrates are represented on the top panels. Residuals from the fits are depicted in the bottom panels. Kinetics were recorded by following fluorescence polarization changes over time using a plate reader. For structures of BG and CP substrates see **Fig. S1.**

**Figure S22:**
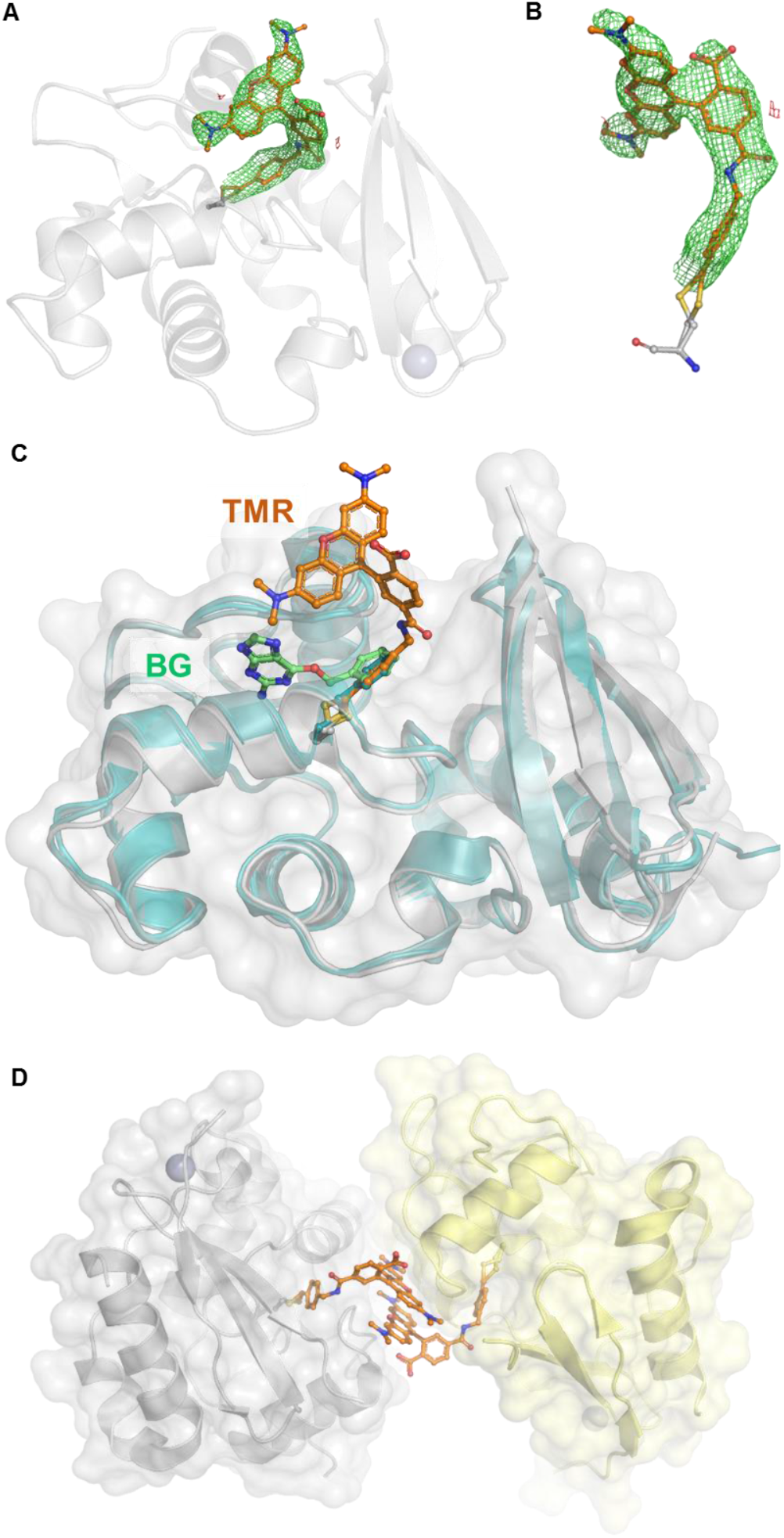
Validation and analysis of the SNAP-TMR X-ray structure. **A.** Omit-map of the TMR ligand of the SNAP-TMR structure. The protein is represented as grey cartoon, TMR fluorophore-substrate as orange sticks and the catalytic cysteine as grey sticks. **B.** Zoom on the isolated labeled catalytic cysteine of SNAP-TMR. Omit-map contoured at 3 σ, represented as green and red mesh for missing and extra density, respectively. **C**. Comparison of the SNAP structure with available SNAP structures. SNAP-TMR is represented as previously explained. Apo SNAP (PDB ID 3KZY), benzylated SNAP PDB ID 3L00) and the BG bound dead mutant SNAP^C145A^ (PDB ID 3KZZ) are represented as cartoon with different shades of blue-green. No major structural differences are observed with SNAP-TMR. **D.** Benzyl-TMR constraints by the crystal packing. Two monomers of SNAP-TMR are represented as grey and yellow cartoons. The conformation of the benzyl-TMR (orange sticks) of both monomers is constrained by the other monomer. Symmetry mates were generated within 4 Å radius and selected to highlight the packing constraints.

**Figure S23:**
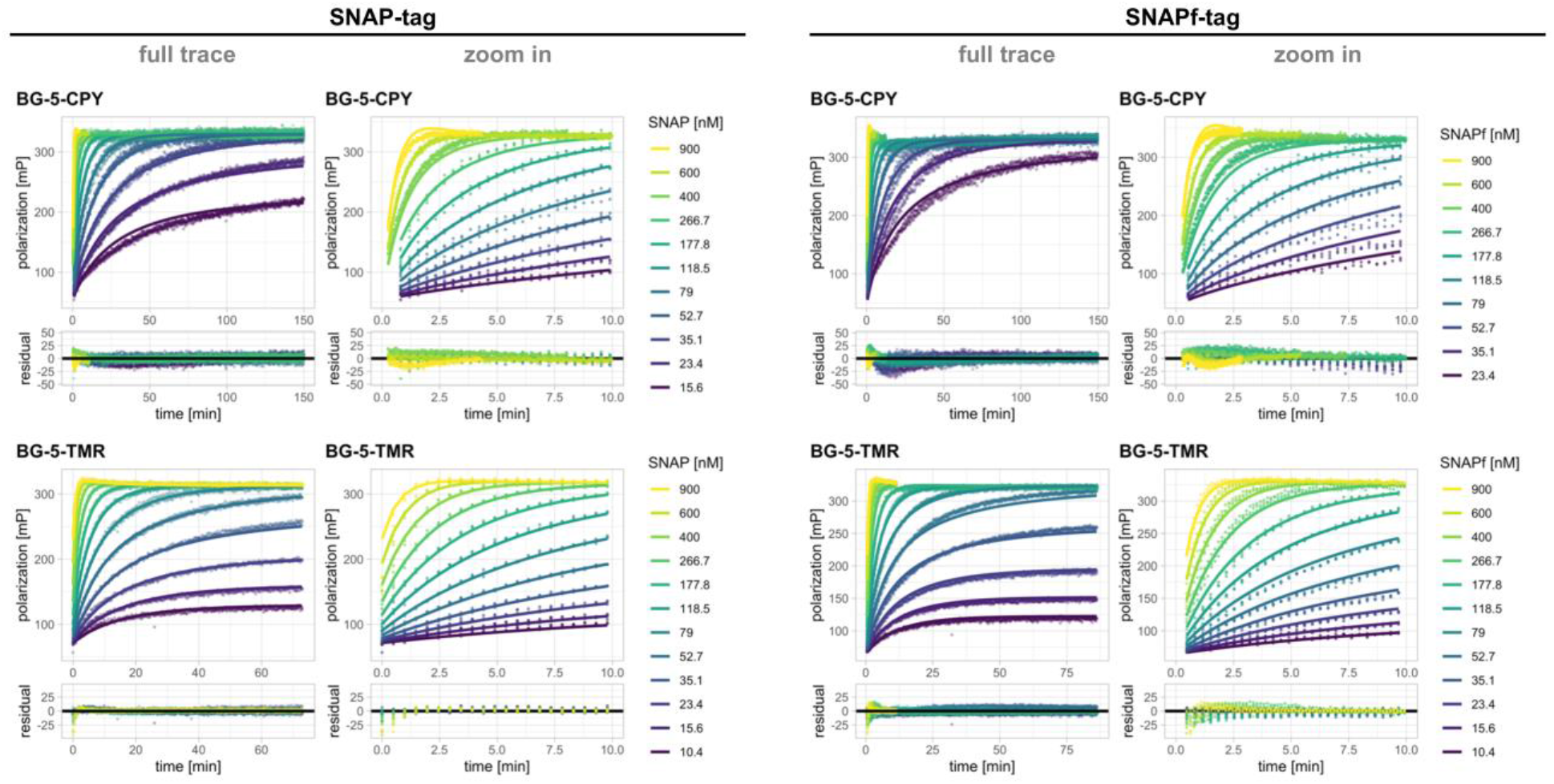
Labeling kinetics of SNAP and SNAPf with BG-5-TMR and BG-5-CPY. Full fluorescence polarization traces (points) and predications of fits based on model 1.2 (lines) along with zoom on the initial 10 minutes are represented on the top panels. Residuals from the fits are depicted in the bottom panels. Kinetics were recorded by following fluorescence polarization changes over time using a plate reader. Labeling was performed at different concentrations of SNAP and SNAPf protein. Substrate concentrations were aimed at 20 nM based on the dyes extinction coefficient but fitted in the model since significant deviations from the expected stoichiometry were observed. For structures of BG substrates see **Fig. S1**.

**Figure S24:**
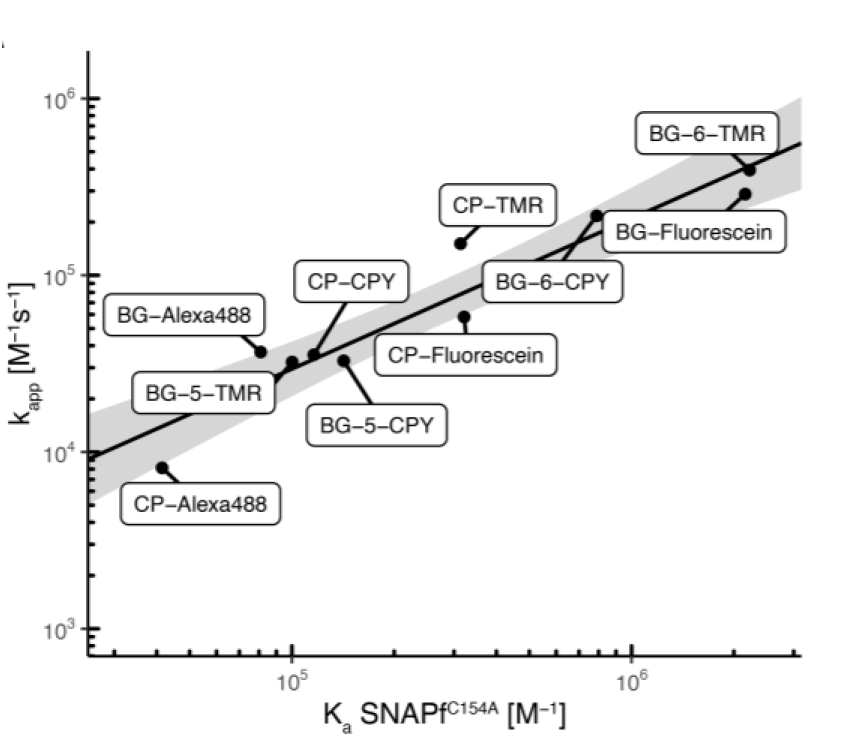
SNAPf kinetic and affinity correlations. Correlation between SNAPf labeling kinetics (k_app_) and affinity (K_a_ = 1/K_d_) for different fluorophore substrates. Affinities were obtained with the catalytically dead mutant SNAPf^C145A^. Log transformed values were fitted to a linear model (black line, log(k_app_) = 0.2568 + log(K_a_) * 1.0697, 95% confidence bands in grey, depicting the area in which the true regression line lies with 95% confidence). The linear correlation in logarithmic space suggests that the K_d_ of fluorescent SNAP substrates towards SNAPf^C145A^ could represent a valid proxy to estimate their K_app_ towards native SNAPf.

**Figure S25:**
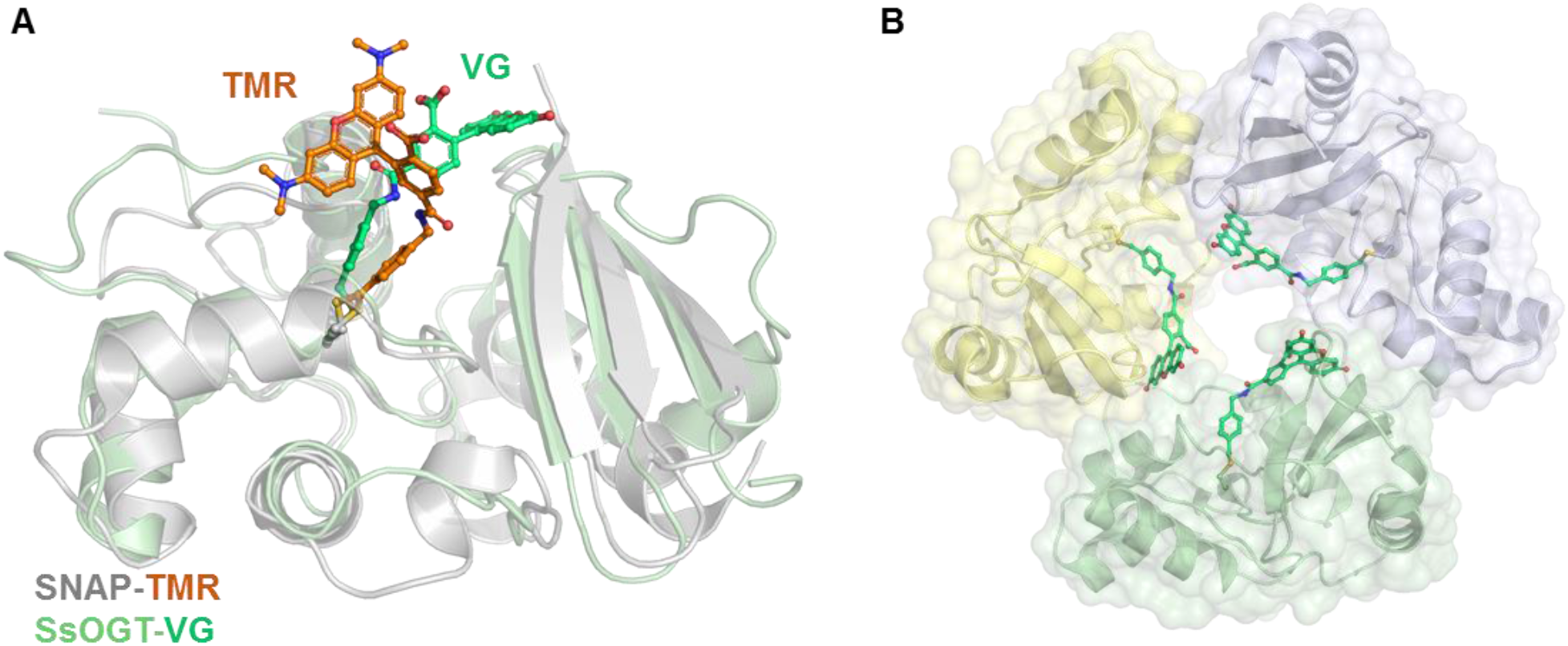
*Ss*OGT-H^5^-VistaGreen alternative fluorophore conformation. **A.** Structural alignment of SNAP-TMR with *Ss*OGT-H^5^-VG structure (PDB ID 6GA0) (17). **B**. Benzyl-VG constraints by the crystal packing. Three monomers of *Ss*OGT-H^5^-VG are represented as blue, green and yellow cartoons. The conformation of the benzyl-VG (green sticks) of all monomers is constrained by the neighboring monomer. Symmetry mates were generated within 4 Å radius and selected to highlight the packing constraints.

**Figure S26:**
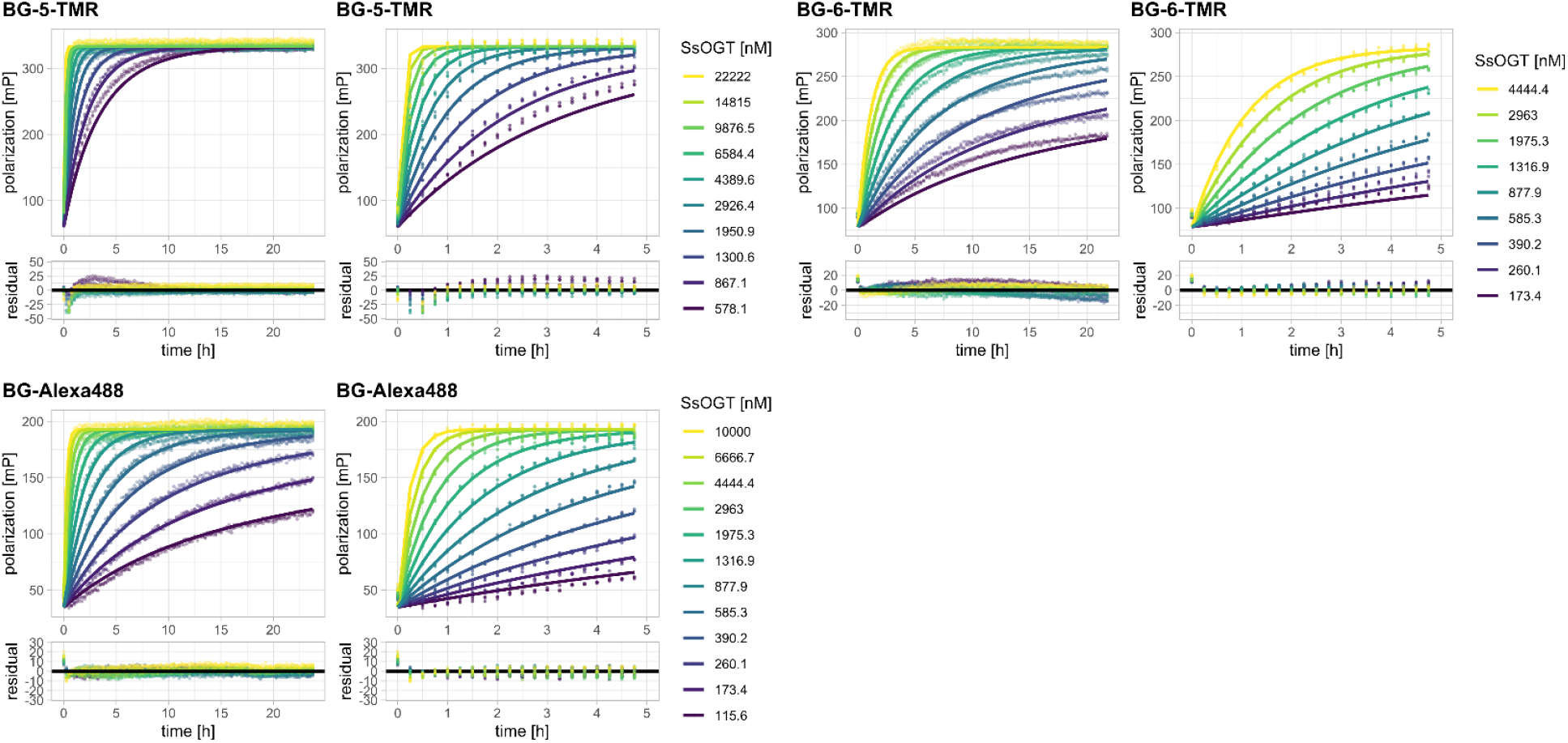
Labeling kinetics of *Ss*OGT-H^5^ with BG-Alexa488 and BG-TMR. Full fluorescence polarization traces (points) and predications of fits based on model 1 (lines) along with zoom on the initial 5 hours are represented on the top panels. Residuals from the fits are depicted in the bottom panels. Kinetics were recorded by following fluorescence polarization changes over time using a plate reader. Labeling was performed at different concentrations of *Ss*OGT-H^5^ protein. Substrate concentrations were aimed at 20 nM based on the dyes extinction coefficient but fitted in the model since significant deviations from the expected stoichiometry were observed. For structures of BG substrates see **Fig. S1**.

**Table S1:** Kinetic parameters of HT7 labeling with fluorescent CA substrates.

| Substrate | $k_1 (\pm \text{S.D.}) [\text{M}^{-1} \text{s}^{-1}]$ | $k_{-1} (\pm \text{S.D.}) [\text{s}^{-1}]$ | $k_2 (\pm \text{S.D.}) [\text{s}^{-1}]$ | $k_{\text{app}} (\pm \text{S.D.}) [\text{M}^{-1} \text{s}^{-1}]$ |
| --- | --- | --- | --- | --- |
| CA-TMR | $7.84 (\pm 0.76) \times 10^7$ | $2.56 (\pm 0.38) \times 10^1$ | $8.06 (\pm 0.29)$ | $1.88 (\pm 0.01) \times 10^7$ |
| CA-JF549 | $1.60 (\pm 0.16) \times 10^8$ | $9.83 (\pm 2.04) \times 10^1$ | $1.82 (\pm 0.07) \times 10^1$ | $1.66 (\pm 0.01) \times 10^7$ |
| CA-JF669 | $2.35 (\pm 0.75) \times 10^7$ | $3.39 (\pm 1.49) \times 10^1$ | $6.94 (\pm 0.47)$ | $4.03 (\pm 0.02) \times 10^6$ |
| CA-CPY | $1.67 (\pm 0.067) \times 10^8$ | $7.60 (\pm 0.98)$ | $9.86 (\pm 0.73)$ | $9.44 (\pm 0.18) \times 10^7$ |
| CA-LIVE580 | $1.74 (\pm 0.05) \times 10^8$ | $1.75 (\pm 0.28)$ | $6.77 (\pm 0.77)$ | $1.39 (\pm 0.03) \times 10^8$ |
| CA-TMR-biotin | $3.69 (\pm 0.25) \times 10^7$ | $2.10 (\pm 0.25) \times 10^1$ | $8.24 (\pm 0.28)$ | $1.04 (\pm 0.01) \times 10^7$ |

Data analyzed using model 2.

**Table S2:** Comparison k_app_ of HT7 labeling kinetics analyzed using models 1 and 2.

| Substrate | $k_{\text{app}} (\pm \text{S.D.}) [\text{M}^{-1} \text{s}^{-1}]$ | |
| --- | --- | --- |
|  | Model 1 | Model 2 |
| CA-TMR | $1.79 (\pm 0.01) \times 10^7$ | $1.88 (\pm 0.01) \times 10^7$ |
| CA-JF549 | $1.46 (\pm 0.01) \times 10^7$ | $1.66 (\pm 0.01) \times 10^7$ |
| CA-JF669 | $3.95 (\pm 0.02) \times 10^6$ | $4.03 (\pm 0.02) \times 10^6$ |
| CA-CPY | $1.10 (\pm 0.02) \times 10^8$ | $9.44 (\pm 0.18) \times 10^7$ |
| CA-LIVE580 | $1.58 (\pm 0.02) \times 10^8$ | $1.39 (\pm 0.03) \times 10^8$ |
| CA-TMR-biotin | $9.00 (\pm 0.04) \times 10^6$ | $1.04 (\pm 0.01) \times 10^7$ |

**Table S3:** Comparison of HT7 and HOB labeling kinetics with fluorescent CA substrates.

| Protein | Substrate | $k_1 (\pm \text{S.D.}) [\text{M}^{-1} \text{s}^{-1}]$ | $k_{-1} (\pm \text{S.D.}) [\text{s}^{-1}]$ | $k_2 (\pm \text{S.D.}) [\text{s}^{-1}]$ | $k_{\text{app}} (\pm \text{S.D.}) [\text{M}^{-1} \text{s}^{-1}]$ |
| --- | --- | --- | --- | --- | --- |
| HT7 | CA-TMR | $7.84 (\pm 0.76) \times 10^7$ | $2.56 (\pm 0.38) \times 10^1$ | $8.06 (\pm 0.29)$ | $1.88 (\pm 0.01) \times 10^7$ |
| | CA-Alexa488 | - | - | - | $2.57 (\pm 0.01) \times 10^4$ |
| HOB | CA-TMR | $4.15 (\pm 0.26) \times 10^7$ | $1.83 (\pm 0.17) \times 10^1$ | $5.05 (\pm 0.13)$ | $8.99 (\pm 0.04) \times 10^6$ |
| | CA-Alexa488 | - | - | - | $8.04 (\pm 0.02) \times 10^7$ |

**Table S4:**
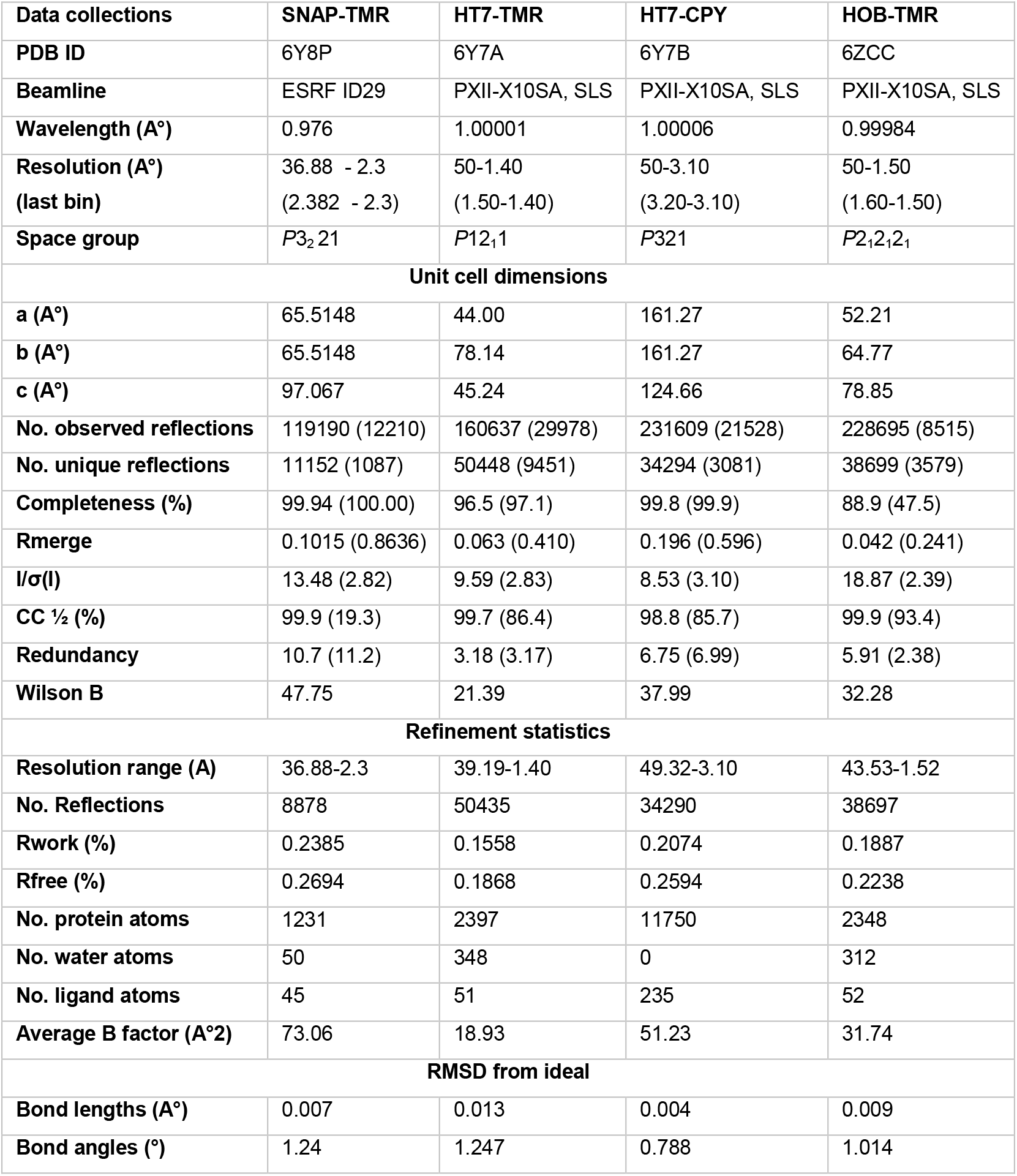
Data collection and refinement statistics the X-ray crystal structures.

| Data collections | SNAP-TMR | HT7-TMR | HT7-CPY | HOB-TMR |
| --- | --- | --- | --- | --- |
| PDB ID | 6Y8P | 6Y7A | 6Y7B | 6ZCC |
| Beamline | ESRF ID29 | PXII-X10SA, SLS | PXII-X10SA, SLS | PXII-X10SA, SLS |
| Wavelength (Å°) | 0.976 | 1.00001 | 1.00006 | 0.99984 |
| Resolution (Å°)<br>(last bin) | 36.88 - 2.3<br>(2.382 - 2.3) | 50-1.40<br>(1.50-1.40) | 50-3.10<br>(3.20-3.10) | 50-1.50<br>(1.60-1.50) |
| Space group | *P*3<sub>2</sub>21 | *P*12<sub>1</sub>1 | *P*321 | *P*2<sub>1</sub>2<sub>1</sub>2<sub>1</sub> |
| Unit cell dimensions |  |  |  |  |
| a (Å°) | 65.5148 | 44.00 | 161.27 | 52.21 |
| b (Å°) | 65.5148 | 78.14 | 161.27 | 64.77 |
| c (Å°) | 97.067 | 45.24 | 124.66 | 78.85 |
| No. observed reflections | 119190 (12210) | 160637 (29978) | 231609 (21528) | 228695 (8515) |
| No. unique reflections | 11152 (1087) | 50448 (9451) | 34294 (3081) | 38699 (3579) |
| Completeness (%) | 99.94 (100.00) | 96.5 (97.1) | 99.8 (99.9) | 88.9 (47.5) |
| Rmerge | 0.1015 (0.8636) | 0.063 (0.410) | 0.196 (0.596) | 0.042 (0.241) |
| I/σ(I) | 13.48 (2.82) | 9.59 (2.83) | 8.53 (3.10) | 18.87 (2.39) |
| CC ½ (%) | 99.9 (19.3) | 99.7 (86.4) | 98.8 (85.7) | 99.9 (93.4) |
| Redundancy | 10.7 (11.2) | 3.18 (3.17) | 6.75 (6.99) | 5.91 (2.38) |
| Wilson B | 47.75 | 21.39 | 37.99 | 32.28 |
| Refinement statistics |  |  |  |  |
| Resolution range (Å) | 36.88-2.3 | 39.19-1.40 | 49.32-3.10 | 43.53-1.52 |
| No. Reflections | 8878 | 50435 | 34290 | 38697 |
| Rwork (%) | 0.2385 | 0.1558 | 0.2074 | 0.1887 |
| Rfree (%) | 0.2694 | 0.1868 | 0.2594 | 0.2238 |
| No. protein atoms | 1231 | 2397 | 11750 | 2348 |
| No. water atoms | 50 | 348 | 0 | 312 |
| No. ligand atoms | 45 | 51 | 235 | 52 |
| Average B factor (Å°2) | 73.06 | 18.93 | 51.23 | 31.74 |
| RMSD from ideal |  |  |  |  |
| Bond lengths (Å°) | 0.007 | 0.013 | 0.004 | 0.009 |
| Bond angles (°) | 1.24 | 1.247 | 0.788 | 1.014 |

**Table S5:** Kinetic parameters of SNAP and CLIP labeling with fluorescent substrates analyzed using model 1.2.

| Substrate | $k_{app} (\pm \text{S.D.}) [\text{s}^{-1}\text{M}^{-1}]$ | $k_3 (\pm \text{S.D.}) [\text{s}^{-1}]$ | $k_{app} (\pm \text{S.D.}) [\text{s}^{-1}\text{M}^{-1}]$ | $k_3 (\pm \text{S.D.}) [\text{s}^{-1}]$ |
| --- | --- | --- | --- | --- |
| SNAP CP-Fluorescein | $1.42 (\pm 0.01) \times 10^4$ | $1.61 (\pm 0.04) \times 10^{-3}$ | - | - |
| SNAP CP-CPY | $1.59 (\pm 0.01) \times 10^4$ | $1.26 (\pm 0.01) \times 10^{-2}$ | $3.55 (\pm 0.02) \times 10^4$ | $6.22 (\pm 0.13) \times 10^{-3}$ |
| CLIP BC-CPY | $1.26 (\pm 0.01) \times 10^4$ | $2.16 (\pm 0.09) \times 10^{-4}$ | $2.65 (\pm 0.01) \times 10^4$ | $9.02 (\pm 0.48) \times 10^{-7}$ |

**Table S6:**
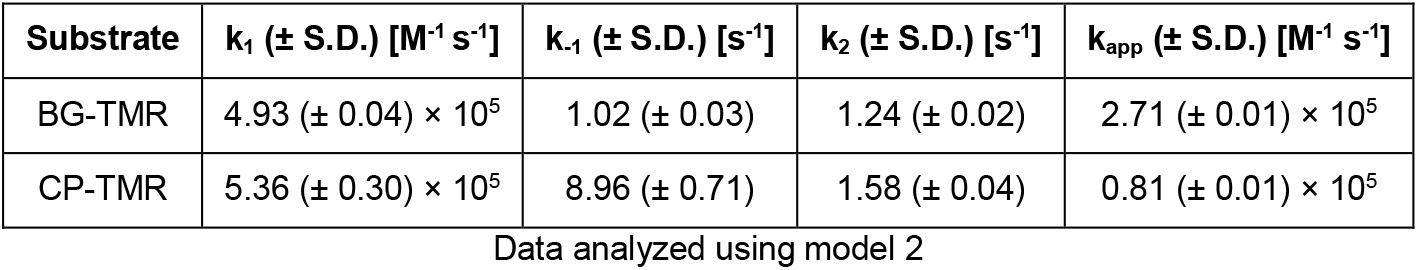
Kinetic parameters of SNAP labeling with TMR substrates measured via stopped flow.

**Table S7:** Comparison of SNAP/CLIP with SNAPf/CLIPf labeling kinetics with fluorescent substrates.

| Substrate | | $k_{app} (\pm \text{S.D.}) [\text{s}^{-1}\text{M}^{-1}]$ | |
| --- | --- | --- | --- |
|  |  | Original | Fast variant |
| SNAP BG substrates | BG-Alexa488 | $1.22 (\pm 0.01) \times 10^4$ | $3.68 (\pm 0.64) \times 10^4$ |
| | BG-Fluorescein | $1.17 (\pm 0.01) \times 10^5$ | $2.88 (\pm 0.01) \times 10^5$ |
| | BG-CPY | $2.17 (\pm 0.01) \times 10^5$ | $2.17 (\pm 0.02) \times 10^5$ |
| | BG-TMR | $4.29 (\pm 0.01) \times 10^5$ | $3.94 (\pm 0.01) \times 10^5$ |
| SNAP CP substrates | CP-Alexa488 | $3.12 (\pm 0.003) \times 10^3$ | $8.13 (\pm 0.01) \times 10^3$ |
| | CP-Fluorescein | $1.42 (\pm 0.01) \times 10^4 (*)$ | $5.81 (\pm 0.01) \times 10^4$ |
| | CP-CPY* | $1.59 (\pm 0.01) \times 10^4 (*)$ | $3.55 (\pm 0.02) \times 10^4 (*)$ |
| | CP-TMR | $7.69 (\pm 0.01) \times 10^4$ | $1.51 (\pm 0.01) \times 10^5$ |
| CLIP BC substrates | BC-Alexa488 | $1.26 (\pm 0.01) \times 10^3$ | $3.10 (\pm 0.02) \times 10^3$ |
| | BC-Fluorescein | $4.36 (\pm 0.01) \times 10^3$ | $1.62 (\pm 0.01) \times 10^4$ |
| | BC-TMR | $1.85 (\pm 0.01) \times 10^4$ | $3.37 (\pm 0.01) \times 10^4$ |
| | BC-CPY* | $1.26 (\pm 0.01) \times 10^4 (*)$ | $2.65 (\pm 0.01) \times 10^4 (*)$ |

Data analyzed using model 1 or 1.2 (\*) which included an additional phase (see Table S5).

**Table S8:** Comparison of SNAP labeling kinetics with 5- and 6-fluorophores.

| Substrate | SNAP |  | SNAPf |  |
| --- | --- | --- | --- | --- |
| | $k_{app} (\pm \text{S.D.}) [\text{s}^{-1}\text{M}^{-1}]$ | $k_3 (\pm \text{S.D.}) [\text{s}^{-1}]$ | $k_{app} (\pm \text{S.D.}) [\text{s}^{-1}\text{M}^{-1}]$ | $k_3 (\pm \text{S.D.}) [\text{s}^{-1}]$ |
| BG-6-TMR | $4.29 (\pm 0.01) \times 10^5$ | - | $3.94 (\pm 0.01) \times 10^5$ | - |
| BG-5-TMR (*) | $2.67 (\pm 0.01) \times 10^4$ | $1.53 (\pm 0.12) \times 10^{-3}$ | $3.23 (\pm 0.01) \times 10^4$ | $2.18 (\pm 0.18) \times 10^{-3}$ |
| BG-6-CPY | $2.17 (\pm 0.01) \times 10^5$ | - | $2.17 (\pm 0.02) \times 10^5$ | - |
| BG-5-CPY (*) | $2.51 (\pm 0.01) \times 10^4$ | $2.11 (\pm 0.04) \times 10^{-2}$ | $3.28 (\pm 0.01) \times 10^4$ | $1.42 (\pm 0.03) \times 10^{-2}$ |

Data analyzed using model 1 or 1.2 (\*) which included an additional phase.

**Table S9:** Kinetic parameters of *Ss* OGT-H^5^ labeling.

| Substrate | $k_{app} (\pm \text{S.D.}) [\text{s}^{-1}\text{M}^{-1}]$ |
| --- | --- |
| BG-6-TMR | $6.78 (\pm 0.67) \times 10^1$ |
| BG-5-TMR | $1.45 (\pm 0.92) \times 10^2$ |
| BG-6-Alexa488 | $1.24 (\pm 0.01) \times 10^2$ |

Data analyzed using model 1.

### Protein sequences

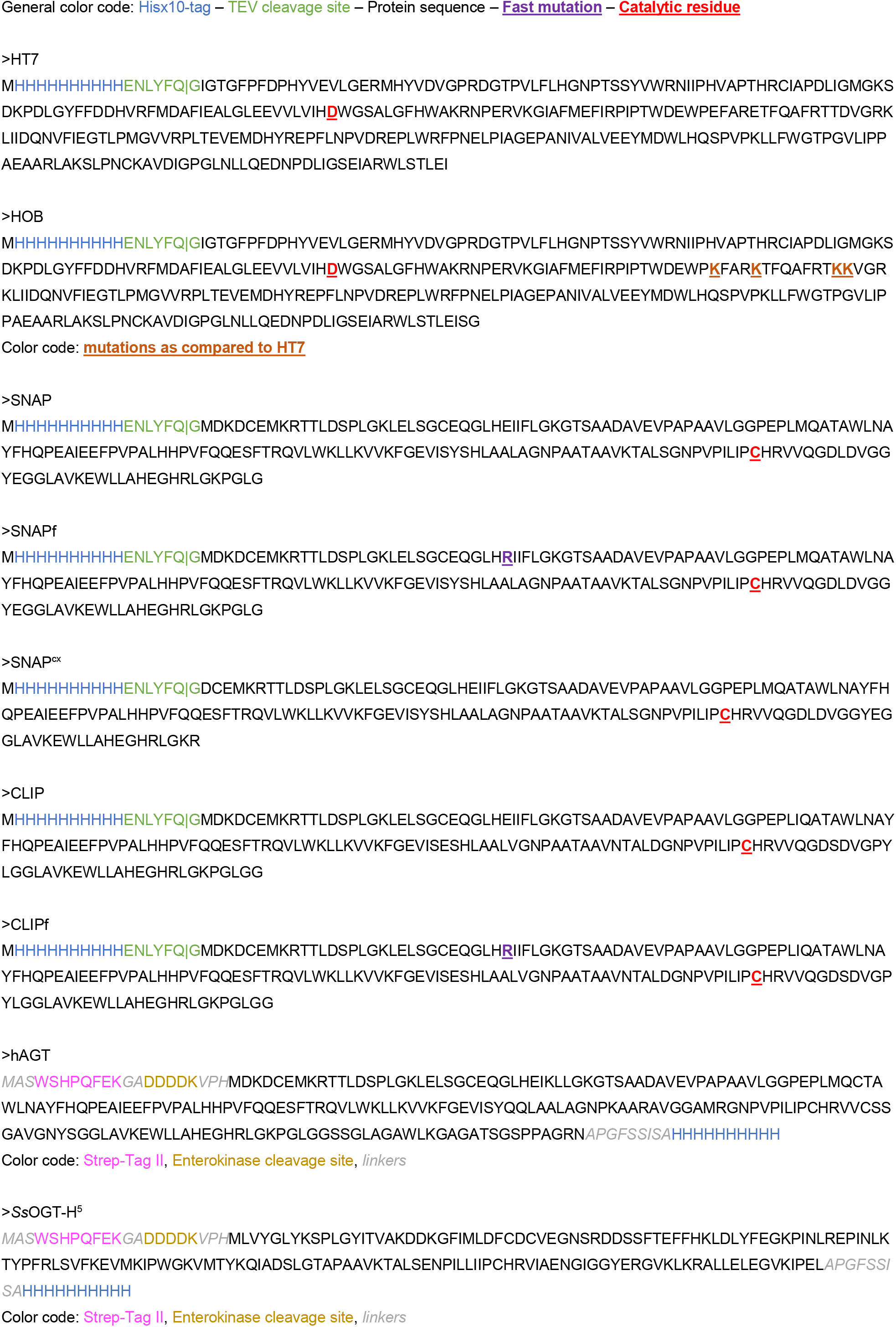

### Example DynaFit scripts

#### HT7 stopped flow labeling kinetics model 2

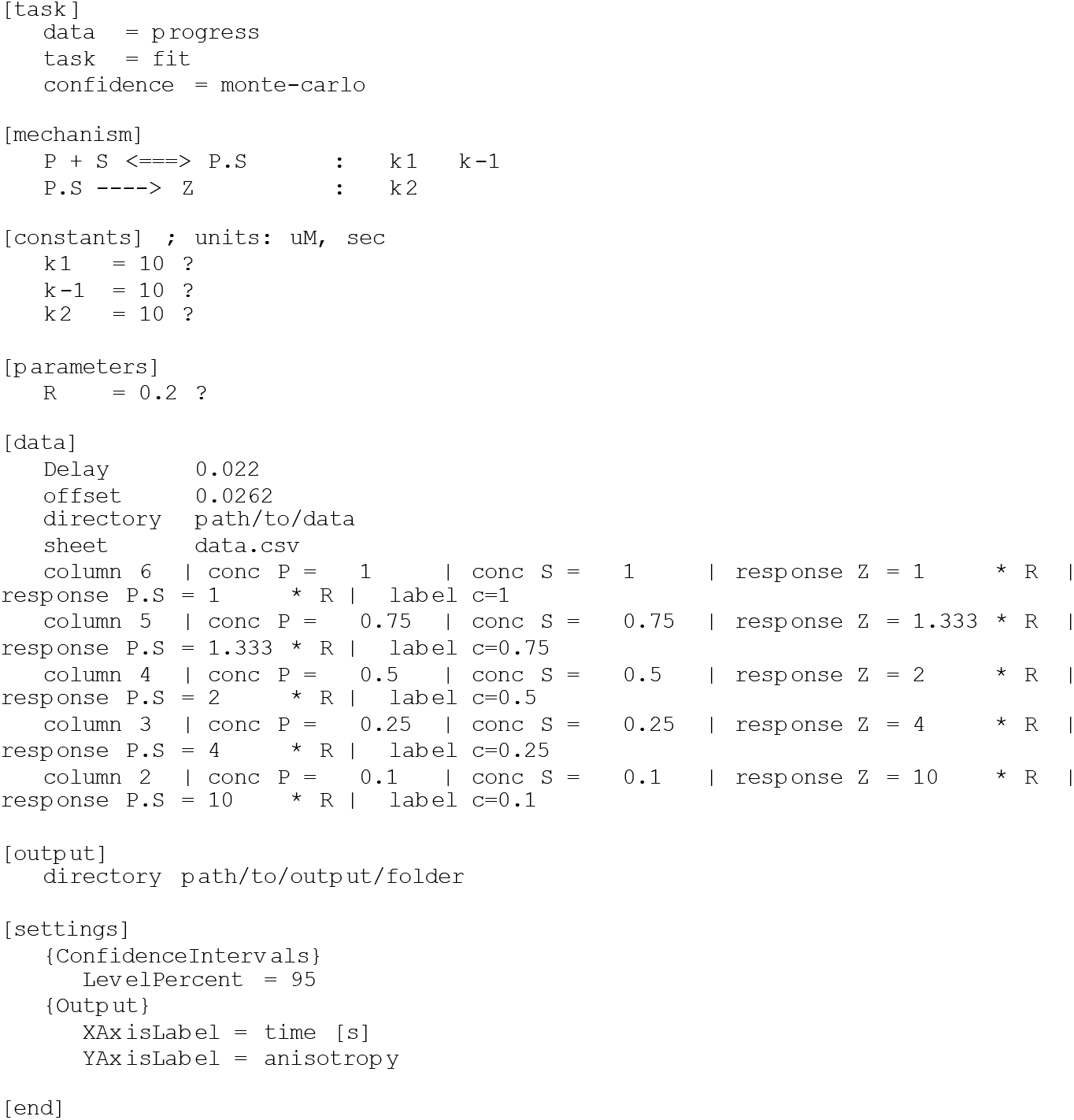

#### SNAP stopped flow labeling kinetics model 2

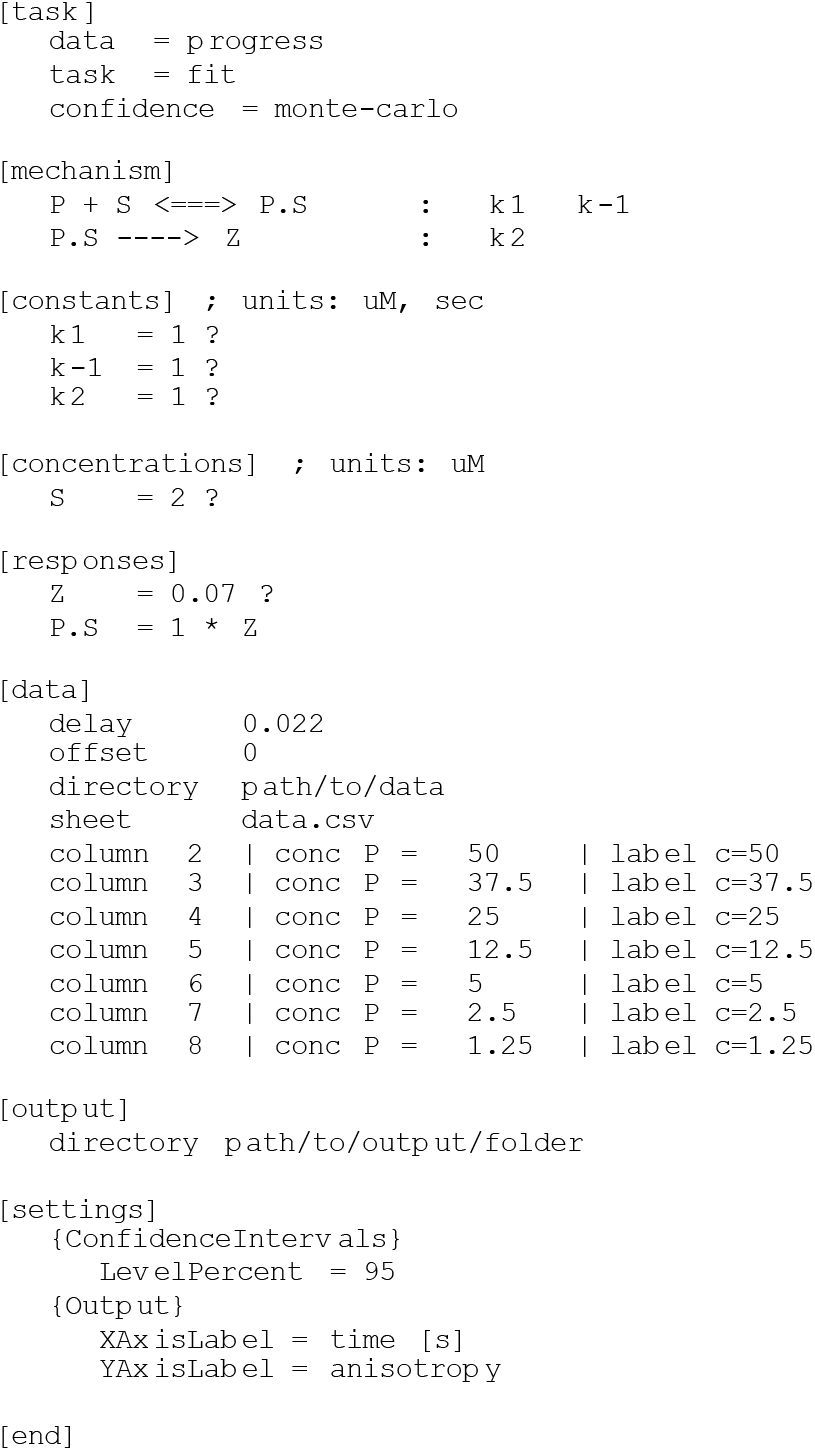

#### HT7 microplate reader labeling kinetics model 1

(Time series of each condition were not averaged before DynaFit analysis since the TECAN plate reader has small inconsistencies in measurement intervals)

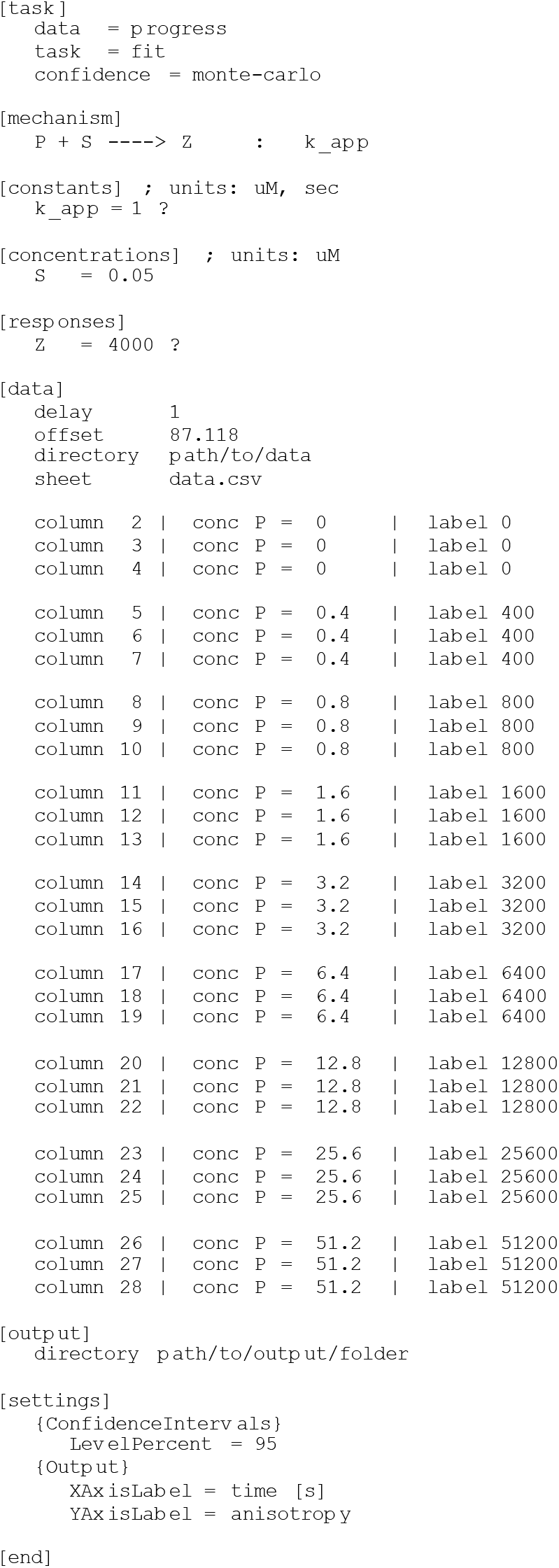

#### SNAP-/CLIP microplate reader labeling kinetics model 1

(Time series of each condition were not averaged before DynaFit analysis since the TECAN plate reader has small inconsistencies in measurement intervals)

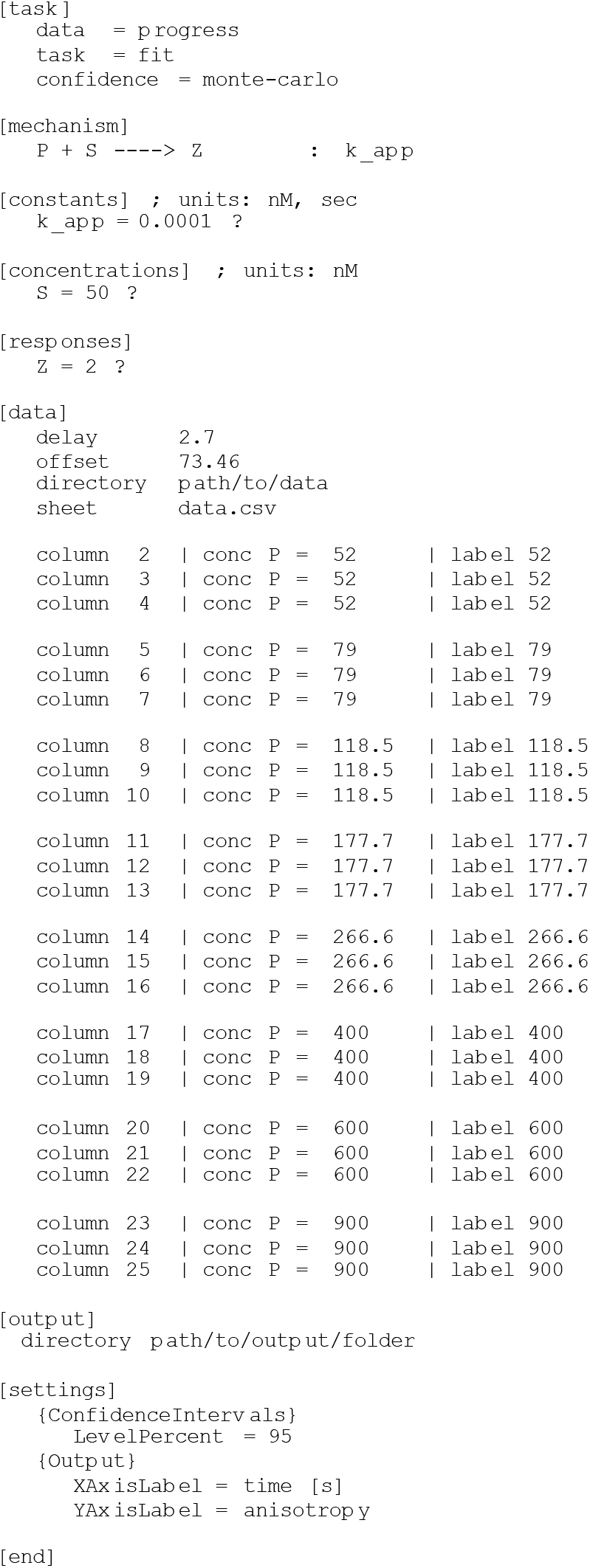

#### SNAP-/CLIP microplate reader labeling kinetics model 1.2

(Time series of each condition were not averaged before DynaFit analysis since the TECAN plate reader has small inconsistencies in measurement intervals)

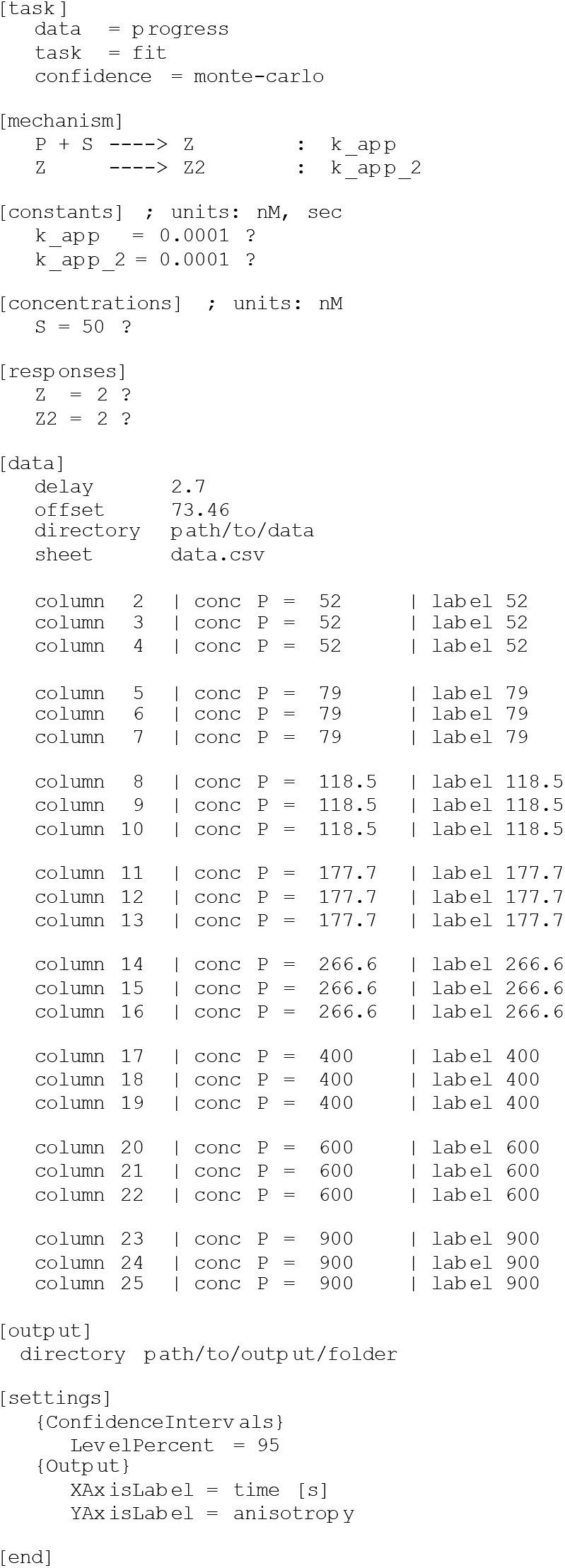

